# Laminin 111 triggers cell quiescence and long-term survival by inducing IQGAP1-mediated cytosolic scaffolding of ERK and BAD inactivation

**DOI:** 10.1101/2024.08.01.606017

**Authors:** Chunhong Yu, Sigita Malijauskaite, Claudia Hinze, Marco Franzoni, Séamus Hickey, Lynnette Marcar, Sew Yeu Peak-Chew, Adam Cryar, Charlie Bain, Jane Marsden, Joanna M. Allardyce, Ana Maria Mendes-Pereira, Harvey T. McMahon, Konstantinos Thalassinos, Kieran McGourty, Emmanuel Boucrot

**Affiliations:** Institute of Structural and Molecular Biology, University College London & Birkbeck College, London, UK; Bernal Institute, University of Limerick, Analog Devices Building, Limerick, Ireland; School of Allied Health, University of Limerick, Limerick, Ireland; MRC Laboratory of Molecular Biology, Cambridge, UK; School of Medical Sciences and Engineering, Beihang University, Beijing, China; SSPC, University of Limerick, Limerick, Ireland; Health Research Institute, University of Limerick, Limerick, Ireland; Limerick Digital Cancer Research Centre, University of Limerick, Limerick, Ireland

## Abstract

In an adult human body, only a minority (∼1%) of cells are dividing; all others are either quiescent, senescent or terminally differentiated. Cellular quiescence, also called G0, is a reversible non-proliferative state in which cells, such as adult stem cells, exist until stimuli trigger their re-entry into the cell cycle. Quiescent cells are known to reside within microenvironment niches of specific extracellular matrix (ECM) composition, but the molecular mechanisms that control their entry and maintenance into G0 and their long-term survival are poorly understood. Here, using a reproducible and homogenous *in vitro* model of quiescence, *ex vivo* tissue histology, phosphoproteomics, and molecular cell biological assays, we revealed that Laminin 111 was sufficient to trigger i) reversible cell cycle exit into G0; ii) sustained and elevated MAPK/ERK signaling; and iii) long-term survival. We found that ERK was activated through the Rap1-BRAF-MEK arm underneath Laminin-binding Integrin α3β1. Activated pERK was scaffolded into the cytoplasm by IQGAP1, thereby blocking its translocation into the nucleus and the activation of proliferative transcription factors. Instead, cytoplasmic pERK inhibited pro-apoptotic protein BAD, which mediated the survival of quiescent cells even in absence of mitogen stimuli. Importantly, we confirmed that pERK was elevated and retained in the cytoplasm of Lgr5^+^ stem cells when they were located within Laminin α1-positive niches in porcine intestine. These findings uncovered a molecular mechanism that may explain how quiescent cell pools, such as dormant adult stem cells, can survive many years despite low mitogen stimuli and be resistant to apoptotic challenges, including chemotherapy.

**HIGHLIGHTS:**

- Laminin 111 is sufficient to induce cellular quiescence (G0) and long-term survival.
- Laminin 111 triggers the sustained and elevated activation of ERK during G0.
- ERK is activated not by growth factor receptors but through the Rap1-BRAF-MEK arm underneath Laminin-binding Integrin α3β1.
- Active, phosphorylated ERK (pERK) is scaffolded by IQGAP1, which prevents it from translocating into the nucleus and activating proliferative transcription factors.
- Instead, cytoplasmic pERK mediates the phosphorylation, and thus inhibition, of BAD, thereby raising the threshold at which G0 cells enter apoptosis.

## INTRODUCTION

A typical human adult body is composed of about 4 x 10^13^ cells, of which less than 1% are proliferating to support tissue and organ homeostasis^1,2^. All other cells have exited the cell cycle momentarily or permanently, and are either quiescent, terminally differentiated or senescent^2^. Terminally differentiated cells have irreversibly exited the cell cycle into specialized lineages such as neurons or muscle cells^3^. Cellular senescence is the irreversible growth arrest response and permanent exit from the cell cycle caused by persistent stress (*e.g.* DNA damage, oncogene activation, telomere shortening), which prevents the replication of old or damaged cells^4^. Cellular quiescence, also called ‘G0’, is the temporary and reversible exit from the cell cycle, a cell cycle state that cells can enter when conditions supporting proliferation are not met^5–7^. As such, cells enter quiescence due to a lack of nutrients, mitogenic stimuli, space (cell-cell contact inhibition) or mechanical support (cell isolation in suspension)^5,8^. Unlike terminally differentiated and senescent cells, quiescence is defined by the cardinal feature of restarting proliferation upon stimulation. Quiescent cells, which include adult stem cell or immune cell pools, persist for long periods (up to several years) in a dormant state but can re-enter the cell cycle during normal tissue turnover or in response to exogenous events (wound healing, immune response, or elevated growth factor stimulation)^5,7^. Thus, the ability of cells to switch between quiescence and proliferation is critical for tissue homeostasis and for responding to disease states.

Quiescent cells are defined by having a diploid (‘2N’) genome, low levels of cell cycle progression markers (*e.g.,* Ki67, PCNA, CyclinD1 or phosphorylated Retinoblastoma protein, hereafter Rb), elevated levels of cyclin-dependent kinase (CDK) inhibitors p21^Waf1/Cip1^, p27^Kip1^ and p57^Kip2^, and decreased levels of mRNA^5,7,9^. Exit from the cell cycle occurs as a bifurcation from G1 upon the inhibition of CyclinD/CDK4/6 and Cyclin E/CDK2/ complexes by elevated p21, p27 and p57, which leads among other molecular events to the hypo-phosphorylation of Rb, which then binds to E2F, repressing cell-cycle gene expression^8^. Quiescent cells also have low mTOR activity, *de-novo* protein synthesis, elevated autophagy, a smaller cell size and an increased volume ratio of nucleus to cytoplasm^10–12^.

While quiescence is the most common cell state among all cellular organisms, our understanding of its molecular mechanisms has lagged behind that of actively dividing cells. This has been caused, in part, by the heterogeneity of *in vitro* culture systems of quiescence that use (in isolation or combinations thereof): nutrient starvation, mitogen deprivation, contact inhibition or loss of adhesion. Although each approach leads to cell cycle exit, they each involve different cellular programs such as mitogen-dependent reduction in proliferation signaling cascades during mitogen removal or mechano-transduction regulatory mechanisms and tight junction signaling after contact inhibition. Interestingly, cell model systems that relied on varying the durations of either contact inhibition or mitogen stimuli, resulted in the generation of heterogenous populations of both deep and shallow quiescent cells - with increased duration being correlated with increased depth of quiescence.

Quiescent cells are known to be able to survive for up to many years, even in the absence of mitogenic stimuli^13^. The lack of mitogenic stimuli decreases Akt kinase activity and increases PTEN phosphatase activity, both of which depress mTOR signaling^6,14^. Lower mTOR activity reduces *de-novo* protein synthesis and relieves its inhibition on autophagy, thereby increasing the recycling of existing proteins and organelles to support long- term survival^15,16^. *In vivo*, the microenvironment surrounding cells is key in maintaining G0: specific extracellular matrix (ECM) composition defines specialized environments, described as niches, in which quiescent cells reside for extended periods of time until they are stimulated to proliferate and differentiate, enabling tissue homeostasis^17–19^. However, the detailed molecular mechanisms of long-term survival during G0 are still unclear and many important questions remain unanswered: What signal transduction pathway(s) mediate ECM-induced long-term survival? Is the mechanism specific to G0, or is it also present in growing cells? Is ECM sufficient, or does it require the concurrent absence of mitogen signaling? Does ECM-Integrin signaling dominate or is simultaneous cell-cell contact signaling required?

Here, using a reproducible *in vitro* model of quiescence, phosphoproteomics, molecular cell biological assays using recombinant ECM and *ex vivo* tissue histology, we revealed that Laminin 111 (‘LM111’) was sufficient to trigger both a reversible cell cycle exit into G0 and long-term survival. It does so through paradoxical sustained and elevated MAPK/ERK signaling though the non-canonical Rap1-BRAF-MEK arm underneath Integrin α3β1. Importantly, activated phosphorylated ERK1 and ERK2 (hence force ‘pERK’) was scaffolded in the cytoplasm by IQGAP1, thereby blocking its ability to translocate into the nucleus to activate proliferative transcription factors. Instead, cytoplasmic pERK phosphorylated and inhibited the pro-apoptotic protein BAD, thereby mediating the long-term survival of quiescent cells.

## RESULTS

### Establishment of an *in vitro* model of cellular quiescence

The best characterized *in vitro* model of quiescence uses human primary neonatal dermal fibroblasts^10,12^ but, for our purpose of studying the molecular mechanisms of ECM-induced long-term survival during G0, we found it to have limitations. First, scaling the protocol to produce enough material for biochemical assays (especially phosphoproteomics and immunoprecipitations) was challenging. Second, genetic manipulations *after* cells entered quiescence was difficult: primary cells are refractory to liposome-based transfection, electroporation of monolayers was not feasible, and CRISPR-Cas9 gene editing likely inefficient (DNA repair by homologous recombination and non-homologous end joining is altered during quiescence^7^) and clone selection and validation are challenging on monolayers. Third, we were concerned that genetic diversity between cell donors, and even heterogeneity from batch to batch, might introduce some level of variability and reduce the reproducibility between experiments.

To address these challenges, we improved upon the existing protocol by introducing the following three changes: i) we used retinal pigment epithelial cells immortalized with the human Telomere Reverse Transferase (hTERT-RPE1), a karyotypically normal, stable, and diploid human cell line that retains its primary characteristics for several hundred division cycles *in vitro* ^20,21^, ii) we used strict serum-free medium to avoid confounding effects of growth factor signaling and iii) we combined contact-inhibition and mitogen withdrawal, and extended the protocol until stabilization of quiescence markers (**Figure S1**).

Our tissue culture protocol^22^was to replate at low density (2.5 10^4^ cells/cm^2^), cells from exponentially growing stocks, and let them proliferate in full serum medium for 7 days (renewing the medium every other day), until they grew from single, isolated cells cultures into homogeneous monolayers (**Figure S1A**). Cells were then maintained for another 10 days in a strict serum-free medium, renewed every other day, thereby removing mitogenic stimuli from growth factors, but not challenging cells with metabolic starvation - culture medium had still excess levels of glucose, amino acids, and other essential nutrients (**Figure S1A**). By day 17, less than 1% of the cells had DNA profile greater than 2n (**Figure 1A**), confirming that the monolayers did not contain significant numbers of cells still progressing through the cell cycle (cells in S, G2 and M all have >2n DNA). To measure protein and phosphorylation levels in G0 extracts, we normalized extracts to correct for the smaller size (about 20% smaller) and lower protein content (about 40% less per cell) of quiescent cells (**Figure S1B-D**). Throughout this study, immunoblots were loaded with equal protein amounts, as determined using GAPDH as a loading control, as we found that it was stable per μg of protein in both growing and G0 cells (**Figure S1E**). Complementing biochemical assays, high-throughput fluorescent microscopy coupled with automated image analysis allowed for the quantitation of high numbers of cells (typically >3,000 growing and >10,000 quiescent cells per condition) and robust statistics throughout this study (*see STAR Methods*).

**Figure 1.**
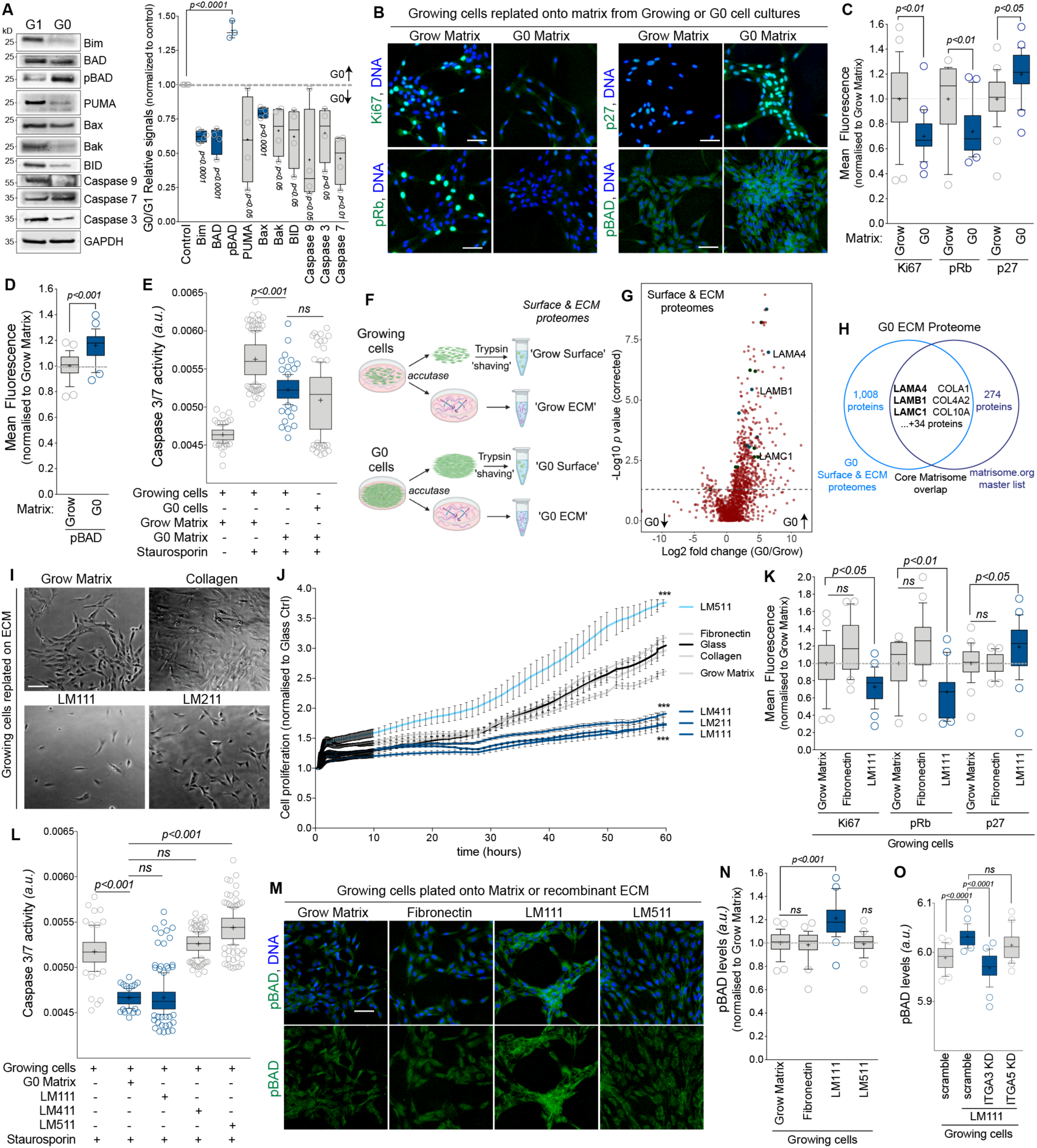
Laminin 111 is sufficient to induce cell cycle exit and long-term survival. (A) *Left*, representative immunoblots showing the levels in apoptosis proteins Bim, BAD, phosphorylated Ser112 BAD (‘pBAD’), PUMA, Bax, Bak, BID, Caspase 9, Caspase 7 and Caspase 7 in G1 or G0 cell extracts (isolated by cell sorting, *see STAR Methods*). GAPDH served as loading control. *Right*, G0/G1 relative signals, normalized to GAPDH of >3 independent immunoblots as in (*Left*). (B) Representative fluorescence microscopy images of growing cells replated onto tissue culture dishes coated by the ECM naturally secreted by growing (‘Grow Matrix’) or quiescent (‘G0 Matrix’) cells previously grown on them. Cells were immunostained for endogenous Ki67, phosphorylated Ser807/811 Retinoblastoma protein (‘pRb’), p27^kip1^ (‘p27’), or phosphorylated Ser112 BAD (‘pBAD’) and DNA (blue). (C) Mean fluorescence of Ki67, pRb or p27 in cells grown as in (B). (D) Mean fluorescence of pBAD in cells grown as in (B). (E) Caspase 3/7 activity in G0 cells or growing cells replated onto Grow or G0 Matrix, challenged or not with 2 μM staurosporin for 6h, as indicated. (F) Schematic of Surface and ECM proteomes. Growing or G0 cells were detached using accutase, then their cell surface proteins were ‘shaved’ using Trypsin and collected. ECM and proteins left on tissue culture dishes were collected and lysed in 6M urea. Proteins were identified and quantified using label-free quantitative mass spectrometry (LC-HDMS^E^). (G) Volcano plot of -Log10 of *p* values versus Log2 of fold changes in protein levels detected by label-free quantitative mass spectrometry (LC-HDMS^E^) from Surface and ECM fractions from Growing or G0 cells. Proteins relevant to this study were annotated. (H) Venn diagram showing the overlap between the proteins detected in (F) and the core human matrisome ^24^. (I) Representative bright field images of growing cells replated onto tissue culture pre-coated with purified Collagen or recombinant Laminin 111 (LM111) or 211 (LM211), as indicated, and grown for 3 days in full-serum medium. (J) Cell proliferation index (normalized to no coating control) of growing cells replated onto Grow Matrix or the indicated recombinant ECM and measured using a xCELLigence RTCA DP E-plate system. (K) Mean fluorescence of endogenous Ki67, pRb or p27 in growing cells replated onto Grow Matrix, purified Fibronectin or recombinant LM111 and grown for 6 days in SF medium. (L) Caspase 3/7 activity in G0 cells or growing cells replated onto G0 Matrix or recombinant LM111, LM411 or LM511, challenged or not with 2 μM staurosporin for 6h, as indicated. (M) Representative fluorescence microcopy images of growing cells replated onto Grow Matrix, purified Fibronectin or recombinant LM111 or LM511 in full serum media for 24 hr before being cultured for a further for 6 days in SF media, and immunostained for endogenous pBAD (green) and DNA (blue). (N) Mean fluorescence of endogenous pBAD in cells gown as in (M). (O) Mean fluorescence of endogenous pBAD in growing cells depleted for Integrin α3 (ITGA3 KD), or Integrin α5 (ITGA5 KD), or treated with control siRNA oligos (scramble), and cultured for 6 days in SF media with or without recombinant LM111. In B, I and M, the scale bars represent 50 μm. In A, C, D, E, K, L, N and O, crosses show the mean, horizontal lines show the median and whiskers show the 10 to 90^th^ percentiles of values. In J, bars show the mean and error bars represent SEM of 3 independent experiments. *P* values were calculated by one-way ANOVA (Tukey and 95% CI for multiple comparison testing), *\*\*\*P<0.001, ****P<0.0001*

Very strong reductions in the proliferative markers Cyclin D1, PCNA, Ki67 and phosphorylated Ser807/811 Retinoblastoma protein (hereafter ‘pRb’) and an increase in p27^Kip1^ (hereafter, ‘p27’), confirmed that our tissue culture protocol produced homogenous populations of quiescent cells (**Figure S1F-I**). Time course experiments showed that Cyclin D1, Ki67 and pRb levels dropped during the first 15 days after cell plating and remained low for the following 10 days (**Figure S1H**). However, p27 levels were already maximal at day 7, well before the aforementioned markers reached their lowest, and thus also before cells entered what we consider stable quiescence (**Figure S1H**). Following this, assays were performed on day 17 or later of the protocol, a stage at which we estimated our G0 cells samples were homogeneously in quiescence.

Finally, we checked that our *in vitro* quiescence tissue culture protocol did not trigger senescence. Consistent with their increased lysosomal activity and gradient in senescence biomarkers^23^ the quiescent cells had higher β−Galatosidase (βGal) levels than growing cells (**Figure S2A-B**). However, βGal levels in G0 cells increased further after treatments with the senescence inducers H2O2 and Doxorubicin. In addition, G0 cells were less positive for p21^Waf1/Cip1^ and phosphorylated γHAX foci, both of which were elevated when senescence was induced chemically (**Figure S2C-F**).

Altogether, this established that our *in vitro* model of quiescence was suitable for producing consistent and homogeneous samples to study molecular mechanisms of long-term survival during G0.

### ECM from G0 cells is sufficient to induce cell cycle exit and long-term survival

In our *in vitro* quiescence cell model, we noticed that the removal of growth factors induced extensive cell death if it occurred before cells reached confluency (**Figure S3A**). Once they had formed a monolayer and entered G0, cells survived the starvation for at least 12 days without detectable cell death (**Figure S3A**). Quiescent cells were also resistant to 6h incubation with 2 μM of the broad kinase inhibitor staurosporine, a challenge that induced apoptosis in growing cells (**Figure S3B**). Cell quiescence conferred an elevated resistance, but not absolute protection, as greater challenges such as 15h incubations eventually triggered the apoptosis of G0 cells (**Figure S3B**). Cell survival during G0 was corroborated by the decrease in pro-apoptotic proteins Bax, Bak, BID and downstream effector Caspase 3, 7 and 9, as well as an increase in the inhibitory phosphorylation of the pro-apoptotic regulator BAD (Ser112, ‘pBAD’) (**Figure 1A**). Thus, in addition to mediating cell cycle exit, cell-cell contact appeared to increase survival capacity. However, cell-cell contact also triggers the synthesis and secretion of specific ECM^12^, which may instead be the origin of such anti-apoptosis signaling.

Therefore, to decouple the contribution of ECM from that of cell-cell contacts, exponentially growing cells were plated at low density onto tissue culture dishes in which either growing or G0 cells were previously cultured. After cell removal, we tested dishes retaining ECM naturally secreted from either growing (‘Grow Matrix’) or G0 cells (‘G0 Matrix’) (**Figure S3C**). In the absence of mitogen stimuli and minimal cell-cell contacts, sparse growing cells plated onto ‘G0 Matrix’ showed markedly reduced proliferation and exited the cell cycle, as measured by a reduction of Ki67 and pRB and a rise of p27 (**Figure 1B** and **S3D-E**). Importantly, despite the strict absence of serum, growing cells plated onto G0 Matrix had elevated pBAD levels and were protected against apoptosis to the same levels as long-term quiescent cultures (**Figure 1C-E** and **S3E**). This revealed that the ECM deposited by quiescent cells was sufficient to induce both cell cycle exit and long-term survival.

### Laminin 111 is sufficient to trigger both cell cycle exit and long-term survival

To identify proteins enriched in G0 Matrix, growing or quiescent cells were detached from the plates on which they were cultivated, and both the ECM left on the plates, and cell surface proteins (shaved off using trypsin) were collected (**Figure 1F** and **S4**). Protein abundances measured by label-free quantitative mass spectrometry (LC- HDMS^E^) identified 1,482 proteins, including 1,008 enriched in samples from quiescent cells (**Figure 1G**, **S5** and **Supplementary Table 1**). Amongst them, 40 proteins were part of the core human matrisome^24^, with the notable presence of some Collagen subtypes and all three chains of LM411 (**Figure 1G-H**, **S5F S5** and **Supplementary Table 1**). Collagen and Laminins are major components of the ECM mediating adhesion, survival and proliferation of many cell types. There are 28 different types of Collagen, and Laminins form a family of 16 heterotrimeric glycoproteins (composed of α, β and γ chains).

When replated onto recombinant ECM, growing cells stopped proliferating on LM111, LM211 and LM411, but not on Fibronectin nor, importantly, on LM511 (**Figure 1I-J** and **S6A-B**). However, only LM111 was sufficient to induce growing cells to exit the cell cycle and enter a G0-like state with low Cyclin D1, Ki67 and pRb, and elevated p27 levels (**Figure 1K** and **S6C-G**). Likewise, if either LM111, LM211 or LM411 could protect sparse growing cells from mitogen-removal cell death, only LM11 protected against acute apoptosis challenge (**Figure 1L**). Importantly, the protection conferred by LM111 was of the same magnitude than that of G0 Matrix, and was accompanied by a similar elevation in anti-apoptotic pBAD (**Figure 1M-N**). Finally, LM111-mediated elevation of pBAD was dependent on the Laminin-binding Integrin α3, but not Fibronectin-binding Integrin α5 (**Figure 1O** and **S6H-I**). Altogether, this set of experiments revealed that, even in the absence of cell-cell contacts, Laminin 111 was sufficient to trigger both cell cycle exit and long-term survival in growing cells.

### MAPK/ERK signaling is elevated during G0

To understand the signaling events that could mediate the long-term survival of G0 cells, we performed SILAC quantitative proteome and phosphoproteome comparing G0 and G1 cells (**Figure S4**). We used G1 cells isolated by cell sorting as we reasoned that, given the juxtaposition of G1 to G0, phosphorylations occurring during the rest of the cell cycle (S, G2 and M) were less relevant to G0. To record legitimate changes in phosphorylation, G0/G1 variations were corrected with their corresponding G0/G1 variations in protein levels. Overall, the relative changes of 8,586 phosphorylation sites from 6,982 proteins between G0 and G1 cells were quantified (**Figure 2A**, **S7-10** and **Supplementary Table 2**).

**Figure 2.**
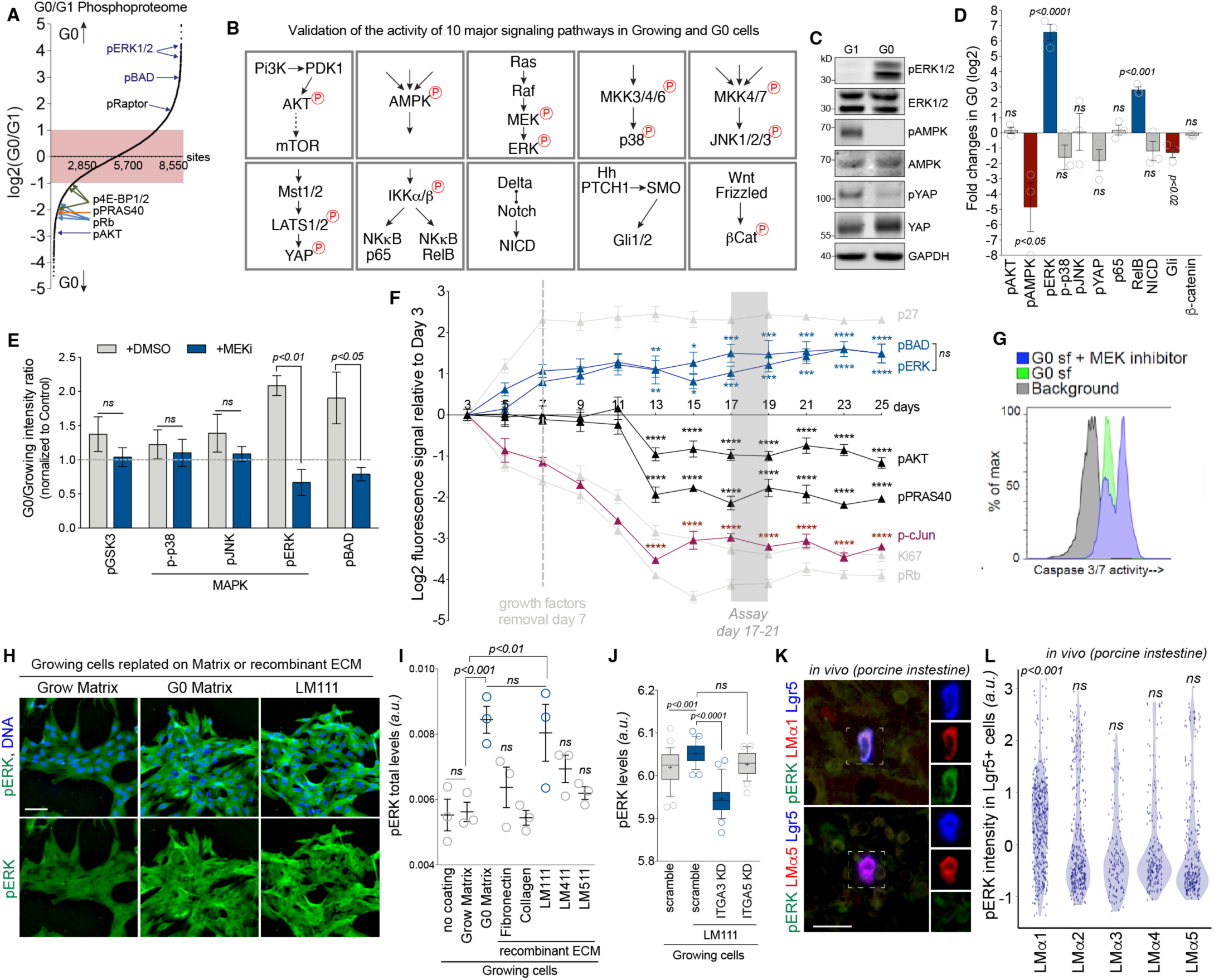
Active ERK is elevated during G0 and mediates long-term survival. (A) ‘S’ plot showing the relative abundance in 8,586 phosphorylation sites measured by SILAC mass spectrometry, and ranked by Log2 normalized ratio (G0/G1). The pink shaded area highlights the range between -1 to +1 ratio, beyond which variations were considered significant (*see STAR Methods*). (B) Schematic of the 10 major signaling pathways investigated in Supplementary Fig. S11. (C) Representative immunoblots showing the levels in phosphorylated ERK, AMPK and YAP (pERK, pAMPK and pYAP, respectively), and their total protein levels in G1 or G0 cell extracts (isolated by cell sorting, *see STAR Methods*). Full data set and quantitation are in Supplementary Fig. S11 and uncropped gels are displayed in Supplementary Fig. S16-18. (D) Fold changes (Log2) between G0 and G1 of 10 major signaling pathways measured by immunoblots from 3 independent experiments. Phosphorylations were corrected by GAPDH loading and their corresponding total protein levels. Full data set and quantitation are in Supplementary Fig. S11 and uncropped gels are displayed in Supplementary Fig. S16-18. (E) G0/Growing intensity ratios of 3 independent experiments measuring the indicated phosphorylations using PathScan Intracellular Signaling Array Kit, from control (left) and cells treated with 2 μM of PD0325901 (MEKi) for 6 hours (right). (F) Mean fluorescence (Log2) form 3 independent experiments of the indicated endogenous immunostainings of quiescent cell cultures measured every 2 days from day 3 over a 25 day period. (G) Flow cytometry profiles of Caspase 3/7 activity in G0 cells grown in serum-free medium (G0 sf) with or without MEKi. (H) Representative fluorescence microcopy images of growing cells replated onto Grow or G0 Matrix or recombinant LM111 and cultured for 6 days in SF media, and immunostained for endogenous pERK (green) and DNA (blue). (I) Mean fluorescence of pERK levels from 3 independent experiments using growing cells replated onto the indicated ECM and grown and immunolabeled as in (H). (J) Mean fluorescence of endogenous pERK in growing cells depleted for Integrin α3 (ITGA3 KD), or Integrin α5 (ITGA5 KD), or treated with control siRNA oligos (scramble), and grown or not onto recombinant LM111. (K) Representative fluorescence microcopy images of porcine intestine sections immunostained for endogenous pERK (green), Laminin alpha chain 1 or 5 (LMα1 or LMα5, red) and the stem cell marker Lgr5 (blue). (L) Mean fluorescence of endogenous pERK levels in Lgr5^+^ stem cells colocalizing with Laminin α1 to 5 (LMα1-5) using 6 porcine intestine histology sections per condition from three different pigs. In H and K, the scale bars represent 50 μm. In D, E and I, bars show the mean and error bars represent SEM of 3 independent experiments. *P* values calculated by one-way ANOVA (Tukey and 95% CI for multiple comparison testing), *\*P<0.05, **P<0.001, ***P<0.001, ****P<0.0001*

Surprisingly, the activatory sites ERK1 and ERK2, Thr202/Tyr204 and Thr185/Tyr187 respectively (‘pERK’) were amongst the phosphorylations most elevated in G0 (**Figure 2A**, **S7D**, **S10** and **Supplementary Table 2**). This was unexpected because the MAPK/ERK kinase signal transduction cascade is typically activated by growth factors (*e.g.* insulin or EGF), which were absent from the G0 cell cultures. In addition, robust activation of this pathway is well characterized as having oncogenic potential associated with dysregulated proliferation. To validate this and gain a more comprehensive picture of the signal transduction happening in quiescent cells, we measured the activity of 10 major signaling pathways using Immunoblotting of G0 and G1 cell extracts (**Figure 2B-D** and **S11**). For each pathway, we probed key phosphorylation sites of the main kinases, normalized by their total proteins levels, or levels of the main effectors. While the p38, JNK, Hippo, and Wnt pathways were not significantly changed, the AMPK and Hedgehog pathways were depressed during G0 (**Figure 2B-D** and **S11**). On the other hand, while the p65-mediated NF-κB pathway was unchanged, the RelB-dependent NF-κB arm was increased, and, indeed, ERK was the most elevated pathway we measured during G0 (**Figure 2D** and **S11**). The strong increase in pERK in G0 cells was not only unexpected from the low mitogen stimuli, but also because MAPK/ERK increase in pERK in G0 cells was not only unexpected from the low mitogen stimuli, but also because MAPK/ERK triggers, amongst other functions, the activation of transcription factors driving cell cycle progression and proliferation, which are known to be inhibited during quiescence^25^. Importantly, MAPK/ERK has also a pro-survival function by phosphorylating, and thereby inhibiting, pro-apoptotic factors^25^. Thus, we tested next whether pERK was inhibiting BAD and mediating long-term survival during G0.

### pERK phosphorylates BAD to support long-term survival during G0

Inhibiting the upstream kinase MEK with PD0325901 (hereafter ‘MEKi’) not only abrogated pERK elevation during G0, but also abolished pBAD, establishing a functional hierarchy (**Figure 2E**). Time-course experiments revealed that pERK and pBAD levels rose in lockstep and maintained constant and elevated levels for at least 25 days, similar to the rise of p27 and drops in Ki67 and pRb (**Figure 2F**). Importantly, the viability of G0 cells was mostly abolished by the addition of MEKi (**Figure 2G**), which showed that ERK signaling and BAD inhibition were at the center of an essential survival axis in the long-term survival of quiescent cells.

The growth factor removal step of our quiescence protocol (Day 7) induced an expected decrease in pAKT and downstream effector pPRAS40, but did not affect pERK and pBAD elevation (**Figure 2F**). Interestingly, levels of activated transcription factor cJun (‘p-cJun’), a downstream target of pERK and major mediator of cell cycle progression^25^, dropped during deep quiescence establishment (**Figure 2F**), suggesting that MAPK/ERK signaling may be biased towards pro-survival during G0.

Importantly, the elevation in pERK seen in quiescent cells was reproduced in sparse growing cells either plated onto G0 Matrix or onto recombinant LM111, and was mediated by Integrin α3 (**Figure 2H-J** and **S12**). This established that pERK mediated the elevation of pBAD induced underneath LM111 and Integrin α3. Furthermore, we validated these findings *in vivo* in Lgr5^+^ stem cells, a cell population which is predominantly quiescent, and which resides in the crypts/glandular regions of porcine intestines^26^. Lgr5^+^ stem cells found to have enriched Lamin α1 in their vicinity had elevated pERK levels when compared to α2-5 chains (**Figure 2K-L**). Altogether, these data established that quiescent cell ECM, which included LM111, induced the sustained elevation of pERK to mediate anti-apoptotic signaling and long-term survival.

### Rap1-BRAF-MEK signaling cascade mediate pERK activation during G0

Next, we investigated the signal transduction cascade between LM111-Integrin α3β1 at the plasma membrane and pERK. Parallel signaling arms involve Ras or Rap1, Raf proteins (ARAF, BRAF or CRAF) and MEK1/2 (**Figure 3A**). First, while levels of GTP-Ras were similar, significantly more active GTP-Rap1 was immunoprecipitated from G0 than from G1 extracts (**Figure 3B**). Second, only active, phosphorylated BRAF form (Ser445, ‘pBRAF’) was elevated in G0 (**Figure 3C**). Total levels of ARAF were decreased in G0, but active phosphorylated forms of ARAF (Ser299, ‘pARAF’) and CRAF (Ser338, ‘pCRAF’) were similar (**Figure 3C**). Third, a BRAF-specific inhibitor (GDC- 0879, ‘BRAFi’) abolished pERK in G0 cells to the same extent as MEKi (**Figure 3D-E**). However, as expected from the known BRAF-CRAF cross-talk^27^, a CRAF-specific inhibitor (GW5074, ‘CRAFi’) had the paradoxical effect of hyperactivating pERK in G0 cells (**Figure 3D-E**). The effect was the opposite to growing cells, where it was BRAFi which increased pERK levels, suggesting that CRAF activity was dominant in growing cells, whereas BRAF mediated most of the activation of ERK during quiescence. Finally, and consistent with their action on pERK, the inhibition of BRAF but not CRAF, blocked pBAD elevation during G0 and removed the protection of G0 cells against apoptosis (**Figure 3F-I**).

**Figure 3.**
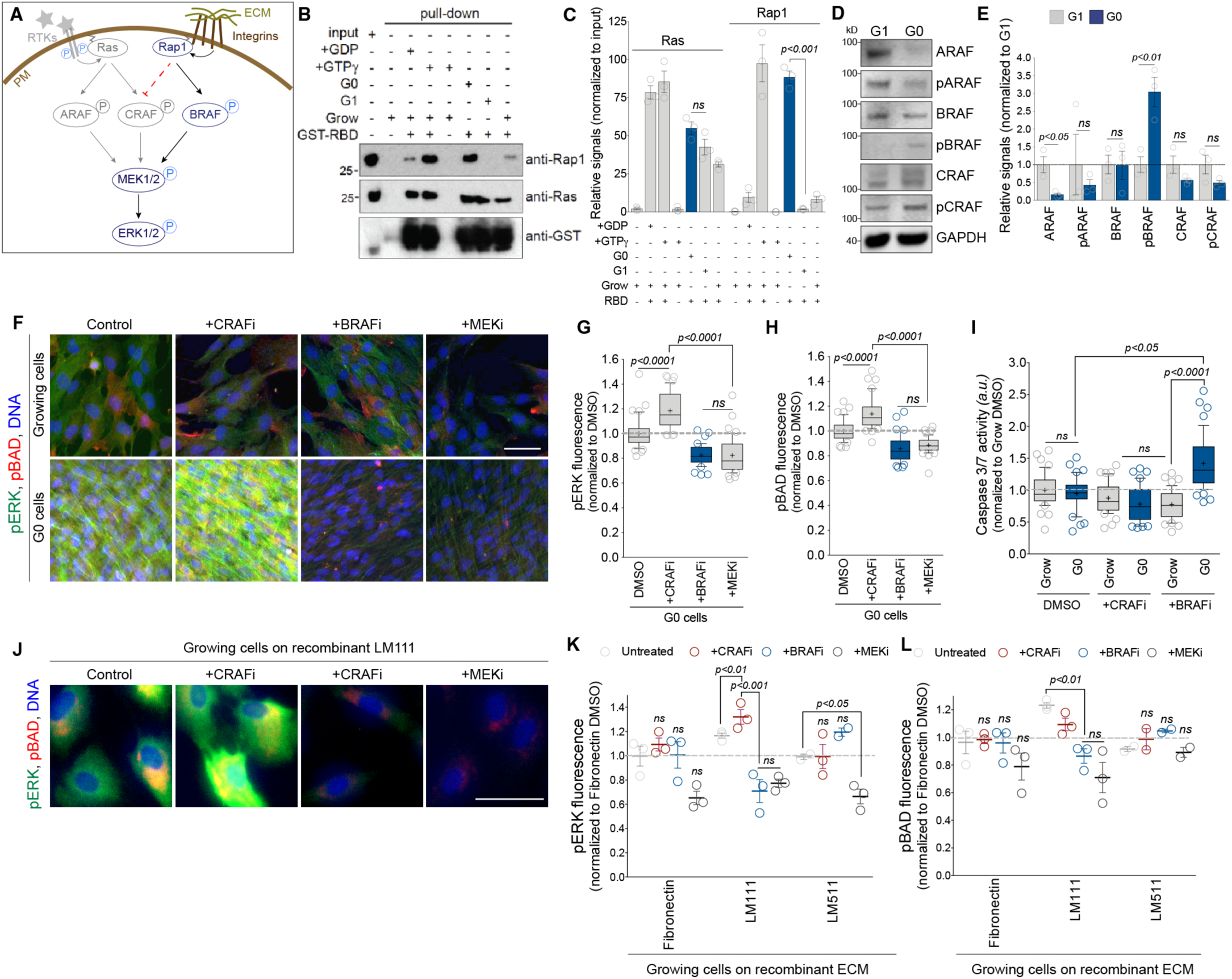
Rap1/BRAF/MEK cascade activates ERK during G0. (A) Schematic of the different arms of the MAPK/ERK pathway. Growth factors activate receptor tyrosine kinase (RTKs), which in turn trigger Ras-ARAF-MEK1/2-ERK1/2 or Ras-CRAF-MEK1/2-ERK1/2 signaling cascades. Extracellular matrix (ECM) binding to Integrins typically trigger the Rap1-BRAF-MEK1/2-ERK1/2 cascade. (B) Representative immunoblots of active GTP-Ras and GTP-Rap1 immunoprecipitation assay using G1 or G0 cell extracts (isolated by cell sorting, *see STAR Methods*). (C) Relative signals normalized to inputs of 3 independent GTP-Ras and GTP-Rap1 immunoprecipitation assays performed as in (B). (D) Representative immunoblots showing the levels in phosphorylated ARAF, BRAF and CRAF (pARAF, pBRAF and pCRAF, respectively), and their total protein levels in G1 or G0 cell extracts (isolated by cell sorting, *see STAR Methods*). Uncropped gels are displayed in Supplementary Fig. S16-18. (E) Relative signals, adjusted to GAPDH loading and normalized to G1, of 3 independent immunoblots as in (of 3 independent immunoblots as in (D). (F) Representative fluorescence microcopy images of growing or G0 cells treated with 2 μM of PD0325901 (MEKi), 2.5 μM of GDC-0879 (BRAFi), or 2 μM of ZM336372 (CRAFi), and immunostained for endogenous pERK (green), pBAD (red) and DNA (blue). (G) Mean fluorescence of pERK expressed as percentage changes relative to untreated control samples, from 3 independent experiments done as in (F). (H) Mean fluorescence of pBAD expressed as percentage changes relative to untreated control samples, from 3 independent experiments done as in (F). (I) Mean fluorescence of Caspase 3/7 expressed as percentage changes relative to untreated control samples, from 3 independent experiments. (J) Representative fluorescence microcopy images of growing cells replated onto recombinant LM111 for 6 days in SF media and treated with MEKi, BRAFi, or CRAFi for the last 6 hr, fixed and immunostained for endogenous pERK (green), pBAD (red) and DNA (blue). (K) Mean fluorescence of pERK expressed as percentage changes relative to untreated control samples, from 3 independent experiments done as in (J) and using purified Fibronectin or recombinant LM111, LM211, LM411 or LM511, as indicated. (L) Mean fluorescence of pBAD expressed as percentage changes relative to untreated control samples, from 3 independent experiments done as in (J) and using purified Fibronectin or recombinant LM111, LM211, LM411 or LM511, as indicated. In F and J, the scale bars represent 50 and 20 μm, respectively. In C, E, G, H, I, K, and L, bars show the mean and error bars represent SEM of 3 independent experiments. *P* values calculated by one-way ANOVA (C, E, G, H and I), or two-way ANOVA (C, E, G, H and I). *\*P<0.05, **P<0.001, ***P<0.001, ****P<0.0001*

When BRAFi and CRAFi were tested in growing cells replated onto recombinant ECM, only LM111 and LM411 could fully recapitulate the phenotypes recorded in quiescent cells (**Figure 3J-L**). Cells on Fibronectin and LM511 were mostly insensitive to both BRAFi and CRAFi, and cells on LM211 displayed a hyperactivation of pERK and pBAD upon BRAFi, but were unaffected by CRAFi (**Figure 3J-L**).

Altogether, this established that during G0, the non-canonical Rap1-BRAF-MEK signaling branch activated pERK and pBAD to mediate long-term survival.

### pERK is sequestered into the cytoplasm during G0

To understand how ERK signaling could be biased away from cell cycle progression and towards pBAD-mediated pro-survival during G0, we measured the translocation of pERK into the nucleus and its activation of downstream transcription factors. As expected, the stimulation of growing cells with 50 ng/mL EGF for 30 min induced: i) the translocation of pERK into the nucleus, and ii) the phosphorylation of cJun, cFos, Creb, Msk1 and Elk1 (**Figure 4A-B**). However, the stimulation of quiescent cells with EGF elevated pERK levels further still, but, surprisingly, without significant increases in pERK nuclear translocation (**Figure 4A-C**). This was confirmed by the absence of phosphorylated downstream transcription factors p-cJun, p-cFos, pCreb and pMsk1 in the nucleus of G0 cells (**Figure 4A-C**). Because Elk1 can also be phosphorylated by the p38 and JNK pathways, the increase in pElk1 in quiescent cells showed that not all transcription is blocked in response to growth factor stimulation (**Figure 4C**). It also suggested that the exclusion of pERK from the nucleus was unlikely to be caused by a broad inhibition in nuclear translocation during G0.

**Figure 4.**
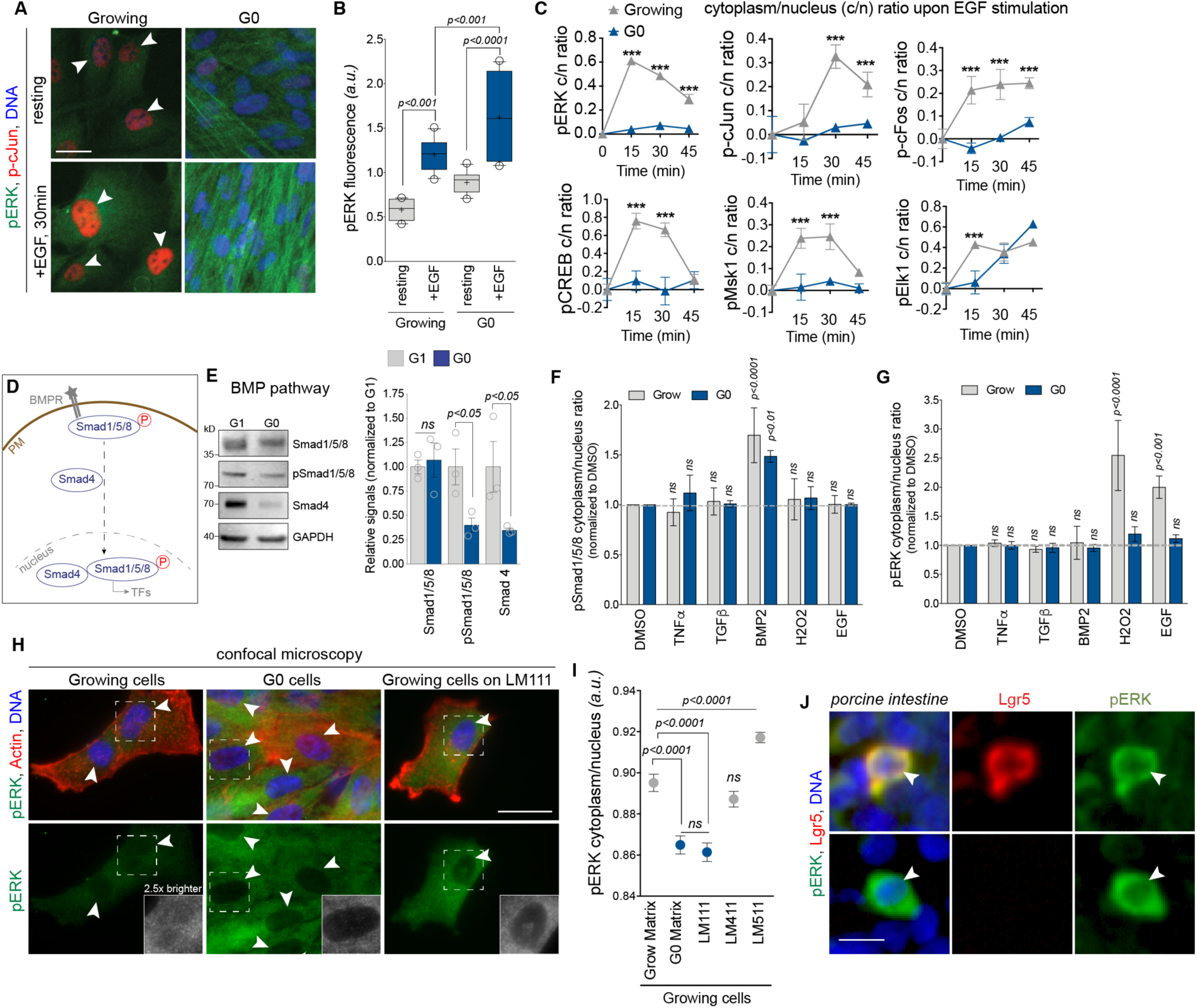
Phosphorylated ERK is sequestered into the cytoplasm during G0. (A) Representative fluorescence microcopy images of growing or G0 cells stimulated or not with 50 ng/mL EGF for 30 min and immunostained for endogenous pERK (green), p-cJun (red) and DNA (blue). (B) Mean fluorescence of pERK from 3 independent experiments performed as in (A). (C) Mean fluorescence from 3 independent experiments of cytoplasmic to nuclear (c/n) ratios normalized to t0 of pERK, p-cJun, p-cFos, pCreb, pMsk1/2 or pElk1 in Growing or G0 cells stimulated with 50 ng/mL EGF for the indicated times. (D) Schematic of Bone Morphogenic Protein (BMP) signaling pathway. BMP binds to BMP receptors (BMPR) to phosphorylates Smad1/5/8, which in turn assemble with Smad4 and translocate into the nucleus where the complex activate transcription factors (TFs). (E) *Left*, representative immunoblots showing the levels in phosphorylated Samd1/5/8 or total protein levels of Smad1/5/8 or Smad4 in G1 or G0 cell extracts (isolated by cell sorting, *see STAR Methods*). Uncropped gels are displayed in Supplementary Fig. S16-18. *Right,* Relative signals, adjusted to GAPDH loading and normalized to G1, of 3 independent immunoblots as in (*Left*). (F) Mean fluorescence of pSmad1/5/8 nuclear versus cytoplasm levels, normalized to DMSO, in Growing or G0 cells, stimulated for 40 min with 20 ng/mL TNFα, 5 ng/mL TGFβ, 10 ng/mL BMP2, 400 μM H2O2 or 50 ng/mL EGF. (G) Mean fluorescence of pERK nuclear versus cytoplasm levels, normalized to DMSO, in Growing or G0 cells treated as in (F). (H) Representative fluorescence confocal microcopy images of G0 cells, or Growing cells replated or not on recombinant LM111 and immunostained for endogenous pERK (green), actin (red) and DNA (blue). (I) Mean fluorescence from 3 independent experiments of cytoplasmic to nucleus ratios normalized of pERK in G0 cells or Growing cells replated and cultured in SF conditions for 6 days on recombinant LM111, LM411 or LM511. (J) Representative fluorescence microcopy images of porcine intestine sections immunostained for endogenous pERK (green), stem-cell marker Lgr5 (red) and DNA (blue). In A, H and J, the scale bars represent 20 μm. In B, E, F, and G, bars show the mean and error bars represent SEM of 3 independent experiments. *P* values calculated by one-way ANOVA (B, C, E, and I), or two-way ANOVA (F and G). *\*P<0.05, **P<0.001, ***P<0.001, ****P<0.0001*

To test this, we compared the translocation of phosphorylated Smad1, 5 and 8 (‘pSmad1/5/8’), the effector complex downstream of BMP pathways, to that of pERK. Even though the total cellular level of pSmad1/5/8 was significantly reduced in resting G0 compared to growing cells, its translocation into the nucleus in response to BMP2 was similar in both cell states (**Figure 4D-F**). As expected, the stimulation of cells with unrelated growth factors TNFα, TGFβ or EGF did not induce the significant nuclear translocation of pSmad1/5/8 in neither G0 nor growing cells. Yet, nuclear translocation of pERK was only elevated in growing, and not in quiescent cells upon EGF stimulation, as well as upon exposure to hydrogen peroxide (H_2_O_2_), a potent trigger of pERK translocation into the nucleus (**Figure 4G**). This confirmed that, while signal transduction effectors can translocate into the nucleus of G0 cells, pERK did not and was retained into the cytoplasm. Indeed, confocal microscopy confirmed that, despite its elevated levels during G0, pERK was excluded from the nucleus and accumulated instead in the cytosol (**Figure 4H**). Strikingly, replating growing cells onto recombinant LM111, but not onto LM411 or LM511, was sufficient to reproduce the phenotype (**Figure 4H-I**), and cytoplasmic pERK was observed in many cells, including Lgr5^+^ stem cells, at the base of villi of porcine intestines (**Figure 4J** and **S13-14**).

### IQGAP1 scaffolds pERK in the cytoplasm to mediate G0 survival

As the nuclear import was found to be functioning during G0, we tested the hypothesis that protein scaffolds could be holding pERK into the cytosol. We tested IQGAP1, as it is a known BRAF/MEK/ERK scaffold and was reported in quiescent cells^28,29^. Indeed, immunoprecipitations with an anti-BRAF antibody isolated an endogenous pBRAF- pERK-IQGAP1 complex in a significantly greater abundance in quiescent than growing cells (**Figure 5A** and **S16a**). As a control, we used 14-3-3ζ, as i) it is another ERK scaffold; ii) like IQGAP1, it was expressed at similar levels in growing and G0 cells (**Fig. S15A**); and iii) it promotes ERK-mediated anti-apoptosis signaling in growing cellsxing^30^.

**Figure 5.**
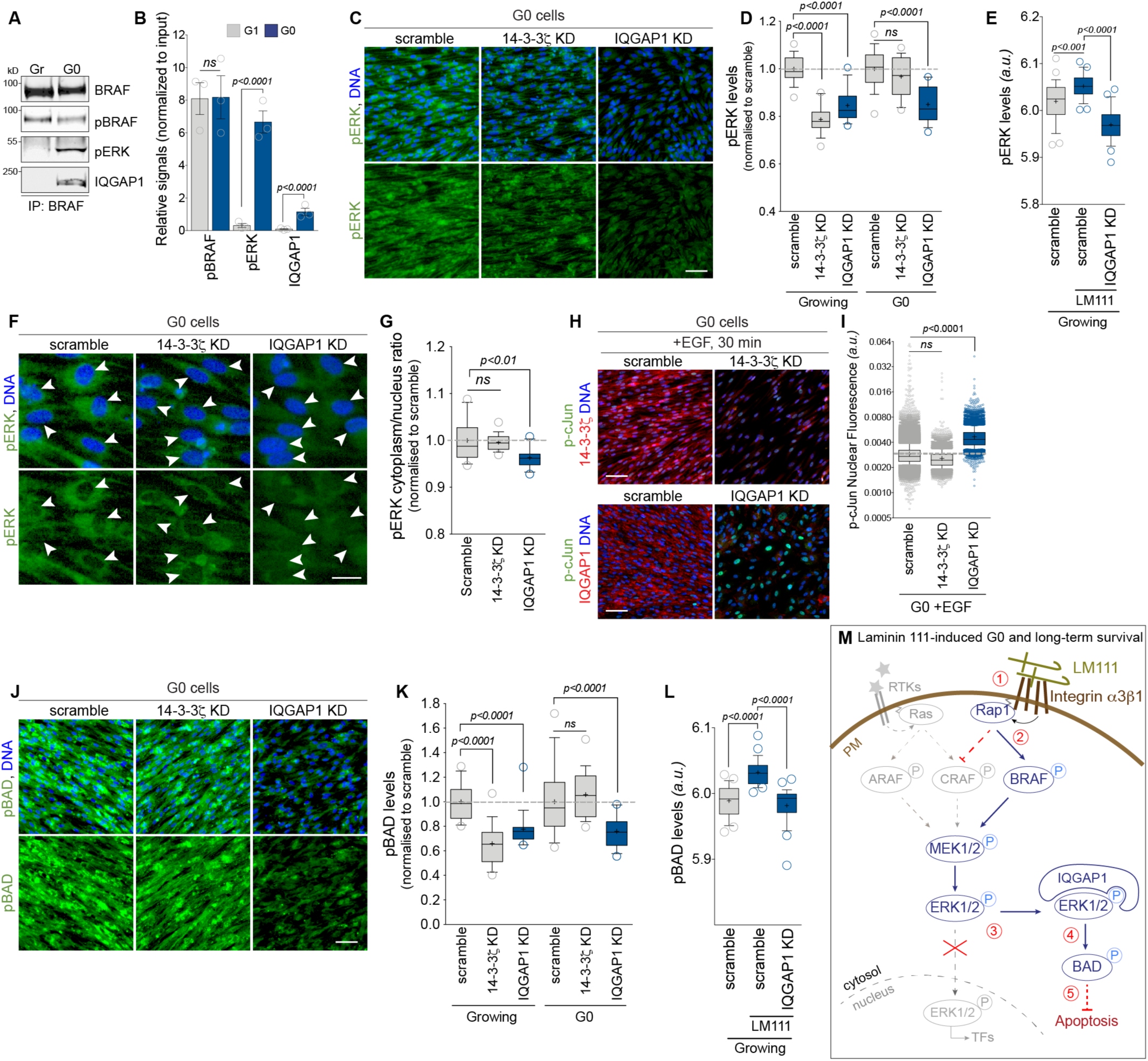
IQGAP1 scaffolds ERK into the cytoplasm during G0 to mediate BAD phosphorylation. (A) Representative immunoblots of immunoprecipitation assays using anti-BRAF antibody and Growing (Gr) or G0 cell extracts and immunostained for endogenous pBRAF, pERK or IQGAP1. (B) Relative signals normalized to inputs of 3 independent immunoprecipitation assays performed as in (A). (C) Representative fluorescence microcopy images of G0 cells depleted of 14-3-3ζ (14-3-3ζ KD) or IQGAP1 (IQGAP1 KD), or treated with control siRNA oligos (scramble), and immunostained for endogenous pERK (green) and DNA (blue). (D) Mean fluorescence of pERK, normalized to scramble, in Growing or G0 cells treated and immunostained as in (C). (E) Mean fluorescence of pERK in Growing cells depleted of IQGAP1 (IQGAP1 KD), or treated with control siRNA oligos (scramble) and replated or not onto recombinant LM111 and cultured in SF media for 6 days. (F) Representative fluorescence microcopy images of G0 cells treated and immunostained as in (C), but the images focused on the exclusion or no of pERK from the nucleus (arrowheads). (G) Mean fluorescence of pERK cytoplasmic to nucleus ratios normalized to scramble in G0 cells treated and immunostained as in (C). (H) Representative fluorescence microcopy images of G0 cells depleted of 14-3-3ζ (14-3-3ζ KD) or IQGAP1 (IQGAP1 KD), or treated with control siRNA oligos (scramble), and stimulated with 50ng/mL EGF for 30 min, and immunostained for endogenous p-cJun (green), 14-3-3ζ or IQGAP1 (red) and DNA (blue). (I) Mean fluorescence of p-cJun in the nucleus of cells treated and immunostained as in (H). (J) Representative fluorescence microcopy images of G0 cells depleted of 14-3-3ζ (14-3-3ζ KD) or IQGAP1 (IQGAP1 KD), or treated with control siRNA oligos (scramble), and immunostained for endogenous pBAD (green) and DNA (blue). (K) Mean fluorescence of pBAD, normalized to scramble, in Growing or G0 cells treated and immunostained as in (J). (L) Mean fluorescence of pBAD in Growing cells depleted of IQGAP1 (IQGAP1 KD), or treated with control siRNA oligos (scramble) and replated or not onto recombinant LM111 and cultured for 6 days in SF media. (M) Summary model. During cellular quiescence, growth factor and receptor tyrosine kinase (RTKs) activity is low, which depress the canonical Ras-ARAF/CRAF arm (grey) of the MAPK/ERK pathway. Instead, 1) LM111 (and potentially other ECM) binds to Integrin α3β1 (and potentially other Integrins), 2) Rap1 activates BRAF, which phosphorylates MEK1/2, which in turn phosphorylates ERK1/2, 3) unlike in growing cells, pERK1/2 does not translocate into the nucleus, and does not activate cell proliferation transcription factors (TFs), 4) instead, pERK remains in the cytoplasm because it is scaffolded by IQGAP1 (and potentially other scaffold proteins). 5) There, pERK phosphorylates, and thus inhibit, BAD, thereby depressing apoptosis and supporting long-term survival of G0 cells. In C, F, H and J, the scale bars represent 50 μm. In B, bars show the mean and error bars represent SEM of 3 independent experiments. *P* values calculated by one-way ANOVA (B, C, E, and I), or two-way ANOVA (F and G). *\*P<0.05, **P<0.001, ***P<0.001, ****P<0.0001*

As expected from their known functions as ERK scaffolds, the depletion of either 14-3-3ζ or IQGAP1 in growing cells reduced ERK activation (**Figure 5B**). In G0 cells, however, only the depletion of IQGAP1 reduced pERK levels and triggered pERK translocation into the nucleus (**Figure 5B-G**). The release of pERK and its nuclear translocation upon IQGAP1 knock-down was confirmed by the restoration in the phosphorylation of cJun in the nucleus of G0 cells (**Figure 5H**). Yet, the depletion of 14-3-3ζ did not relieve the G0-specific inhibition in p-cJun, even upon EGF stimulation (**Figure 5H**). Mirroring these nuclear events, pBAD levels in the cytosol of G0 cells were strongly reduced by IQGAP1 but not 14-3-3ζ KD (**Figure 5J-K**). Finally, depleting IQGAP1 was sufficient to reverse the elevations in pERK and pBAD induced by recombinant LM111 in growing cells (**Figure 5E and L**).

Altogether, we established that during G0, IQGAP1 scaffolded pERK into the cytoplasm, thereby abrogating its translocation into the nucleus and blocking its activation of proliferative transcription factors. This restricts ERK activity to cytoplasmic targets, such as BAD, and supports the long-term survival of quiescent cells.

## DISCUSSION

Based on these data, we propose the following model: during cellular quiescence, growth factor and receptor tyrosine kinase (RTKs) activity is low, which depresses the canonical Ras-ARAF/CRAF arm of the MAPK/ERK pathway. Instead, LM111 and Integrin α3β1 activate the following cascade (**Figure 5M**, step 1): GTP-Rap1 activates BRAF, which phosphorylates MEK, which in turn phosphorylates ERK (**Figure 5M**, step 2). Unlike in growing cells, active pERK does not translocate into the nucleus, and does not activate cell proliferation transcription factors such as cJun, cFos, CREB or Msk1 (**Figure 5M**, step 3). Instead, pERK remains in the cytoplasm because it is scaffolded by IQGAP1 (**Figure 5M**, step 4). There, pERK inhibits BAD, thereby depressing apoptosis and supporting the long-term survival of G0 cells (**Figure 5M**, step 5).

Specific ECM compositions of stem cell niches and of cell microenvironments are known to mediate cell-cycle exit and dormancy^31,32^. Quiescent cells themselves have upregulated the production and secretion of ECM, either to sustain their own quiescence or that of neighboring cells^12,32^. In this study, LM111 was associated with intestinal Lgr5^+^ stem cells that have elevated pERK and was the most potent and consistent at recapitulating G0- like traits in sparse growing RPE1 cells. However, it will undoubtedly not be the sole ECM with such activities. Our data suggest that at least LM211 and LM411 may also have some role in inducing cell cycle exit and long-term survival during quiescence.

The contribution of specific Collagen subtypes will also likely be important - Collagen IV was a strong hit in our mass spectrometry analysis of the ECM secreted by G0 RPE cells, and it is a known interactor and spatial organizer of several Laminins, and a prominent component of the niches of several quiescent stem cells ^32,33^. Other Collagen subtypes may be involved, such as COL17A1 which mediates the dormancy of Lgr5^+^ cancer stem cells^19^. Depending on the cell type, various Collagen and Laminins subtypes may be secreted, perhaps in precise stoichiometries. The potential importance of ECM combinations may be suggested from experiments done using Matrigel, a LM111- and Collagen IV-rich 3D gel extracted from Engelbreth-Holm-Swarm tumors in mice^34^. Normal mammary epithelial cells grown in Matrigel form quiescent acinar structures within 48h in insulin-and hydrocortisone-containing medium, albeit that they have reduced pERK levels^31,35^, which was opposite to our results with pure recombinant LM111 under strict serum-free conditions. However, a proteomic analysis of Matrigel showed that it is a complex ECM mix of 1,851 unique proteins, and although LM111 was the most abundant, Laminin α3, α4, α 5 and β2 chains were detected as well^36^. The consistently opposite phenotypes induced by recombinant LM511 in our experiments suggest that Laminin α5 might counter the effects of LM111, possibly even at sub-stoichiometry. Further, a nuanced picture must also be considered when identifying receptors sensing quiescence-inducing ECM: RPE1 cells in this study relied on Integrin α3β1 and not on Fibronectin-binding α5β1, but other Laminin-binding Integrins such as α6β1, α6β4 or α7β1 may be important as well.

Interestingly, malignant cells resist Matrigel-induced quiescence, probably because of their elevated expression in the ECM-degrading enzymes Matrix-Metalloproteinases (MMP)^37,38^. In particular, oncogenic Ras-mediated canonical CRAF/MEK-ERK signaling increased MMP-9 expression, which degraded LM111 and mediated the escape of malignant mammary epithelial cells from the quiescence-inducing effect of Matrigel^39^. Consistently, LM111 levels are reduced in several stages of breast cancer^40^.

The finding that LM111 was a potent inducer of long-term survival of quiescent cells through the BRAF arm of the ERK pathway may be linked to the known side effects of chemotherapies targeting BRAF, perhaps through a vulnerability of dormant stem cells that may rely on wild-type BRAF for their long-term survival. For example, Sorafenib, the first FDA-approved BRAF inhibitor, caused the development of skin lesions ^27^. Although mutations in ARAF, CRAF and MEK1/2 are rarely reported in human tumors, BRAF is mutated in the majority of malignant melanoma or papillary thyroid carcinoma patients, as well as in some colorectal carcinomas^27^. All mutations increase its kinase activity, such as the prevalent V600E substitution, and inhibitors targeting BRAF will equally affect both mutant and wild-type forms. Moreover, our results suggest that high BRAF activity on itself was likely not sufficient to drive oncogenic proliferation if downstream ERK is sequestered in the cytosol and does not activate cell-cycle transcription factors in the nucleus.

Thus, a key regulation in the mechanism proposed here is likely to be at the level of the scaffolding of pERK and its retention in the cytoplasm. This phenotype, which we confirmed in Lgr5^+^ stem cells within the crypt region of porcine intestine, is at the heart of the biased signaling of ERK, away from nuclear cell-cycle inducing transcription factors and towards cytoplasmic apoptotic targets, including BAD. As we did not detect changes in total levels of IQGAP1 during quiescence, a simple titration effect can probably be ruled out, and post-translational regulations, such as phosphorylation, are more likely. Moreover, other scaffolds forming different pools of ERK are likely involved. In growing cells, about 20 ERK scaffolds have been reported and the single deletion of each of them was enough to release ERK to translocate into the nucleus^41^. This suggests several separate and parallel pools, as the release of few copies of ERK will be enough to trigger a robust phosphorylation of nuclear targets - a single kinase molecule can phosphorylate many substrates in series. During quiescence, inactive ERK is maintained in the cytoplasm by MEK1, PEA-15 and by the Phosphatase MKP3 since they all contain nuclear export signals, and shuttle ERK back into the cytoplasm in a Ran-GTP-dependent, and Exportin-1-dependent manner ^42–44^. On the other hand, LM111 decreases the level of nuclear actin by upregulating its export by Exportin-6 ^45^.

However, the mechanism for the nuclear exclusion of pERK induced by LM111 reported here appears to be different, as the depletion of IQGAP1 was sufficient to release it, indicating a sustained retention by the scaffold in the cytoplasm instead of an active expulsion from the nucleus.

Overall, the mechanism proposed here links the effects of specific ECM, including LM111, to an intracellular signaling using the Rap1/BRAF/MEK/ERK cascade biased towards anti-apoptotic regulation in the cytoplasm without activation of cell-cycle genes in the nucleus. Importantly, both are required: the serum starvation of sparse growing cells induces a rebound in pERK levels, around 48h after an initial drop, and its retention in the cytoplasm ^46^. But this is not sufficient to block apoptosis and such cells eventually die within days of serum starvation. This is likely because growing cells did not have the trigger to upregulate and secrete ECM supporting quiescence and survival, a step that requires contact-inhibition^12^. By bypassing this step and replating growing cell directly onto recombinant LM111 under strict growth factor-free conditions, we could reveal signal transduction events without the confounding effects of cell-cell contact signaling, or residual extrinsic growth factors or culture supplements. This clearly established that the long-term survival of quiescent cells was directly mediated by specific ECM, such as the ones found in stem cell niches. This may explain how quiescent cell pools, such as dormant adult stem cells, could survive many years despite low mitogen stimuli and be resistant to apoptotic challenges, including chemotherapy.

## ACKNOWLEDGMENTS

We thank the members of the McGourty and Boucrot labs for helpful comments. The G2-Si ion mobility mass spectrometer was purchased with a grant from the Wellcome Trust (104913/Z/14/ZBM) to K.T; K.McG was a recipient of a Enterprise Ireland Innovation Partnership (EI IPP IP 2018 0766) and a HRI Seed Award 2018, and E.B. was a BBSRC David Phillips Research Fellow (BB/R01551X), a Lister Institute Research Fellow and a recipient of a BBSRC Pathfinder grant (BB/R01552X) and of a Birkbeck / Wellcome Trust Institutional Strategic Support Fund (ISSF2) Career Development Award.

## AUTHOR CONTRIBUTIONS

C.Y., C.H, L.M., K.McG. and E.B. performed biochemical assays and analysis; C.Y., C.H, C.B., K.McG. and E.B. performed cell biology experiments and analysis; C.Y., C.H., S.M., S.H., M.F, L.M, J.A and K.McG. performed image acquisition and analysis; S.Y.P-C. performed SILAC mass spectrometry under the supervision of H.T.McM., A.C. performed label-free mass spectrometry under the supervision of K.T.; S.M., J.M., A.M.M-P. and K.McG. performed bioinformatics analysis; M.F., S.H. S.M. and K.McG. performed *ex-vivo* sample preparation, image acquisition and analysis; K.McG. and E.B. designed the research and supervised the project. E.B. assembled the figures and wrote the manuscript with input from all the other authors.

## STAR METHODS

### Cell culture and induction of cellular quiescence

Exponentially growing *Human,* normal and diploid, telomerase reverse transcriptase (hTERT)-immortalized retinal pigmented epithelial (RPE-1) cells (ATCC CRL- 4000, called ‘RPE1’ in this study) cells were cultured in ‘Full Medium’ (DMEM:F12 HAM 1:1v/v (Sigma D6421), 0.25% Sodium bicarbonate w/v (Sigma Aldrich S8761), 1mM GlutaMAX-I (Thermo Fisher Scientific A1286001), 1X antibiotic-antimycotic (Thermo Fisher Scientific 15240062), and 10% Fetal Bovine Serum (FBS; Thermo Fisher Scientific 11570506). Exponentially growing RPE1 cells were passaged three times a week by 10 min incubation in EDTA-based cell dissociation buffer (Millipore S-014-B) at 37°C, resuspension in Full Medium and re-seeding at a 1:5 to 1:10 ratio.

Detailed protocol to induce proliferative quiescence was published elsewhere^22^. Briefly, RPE1 cells were counted with a CASY Cell Counter and Analyzer Model TT (Roche Applied Science) and seeded at a density of 2.5 x 10^4^ cells/cm^2^ and grown in Full Medium for 7 days to reach confluence. Full Medium was then exchanged for ‘Serum-free Medium’ (DMEM:F12 HAM 1:1v/v, 0.25% Sodium bicarbonate w/v, 1mM GlutaMAX-I, 1X antibiotic-antimycotic), in which the cells were maintained for at least 10 days to induce long-term quiescence (**Figure S1A**). Both Full Medium and ‘Serum-free Medium’ were renewed every three days. For detachment of the quiescent monolayer, cells were rinsed with phosphate-buffered saline (PBS) and incubated in 0.05% trypsin 0.02% EDTA (Sigma-Aldrich 59428C) or, to preserve cell surface receptors, in Accutase solution (Innovative Cell Technologies AT106-500), both for 5-10 min at 37°C. Serum-free Medium was added to dilute the Trypsin/EDTA or Accutase solution and cell clumps were dissociated by gentle aspiration. Cells were centrifuged for 5 min at 1000*g* and resuspended in FBS-free medium to obtain single-cell solutions. All the cells were maintained at 37°C, 5% CO2. Cells were regularly tested for mycoplasma contamination by nuclei staining with DNA-intercalating fluorescent dyes followed by high resolution confocal microscopy or by PCR with mycoplasma DNA-specific primers.

### Extracellular matrix coating of tissue culture surface

Human recombinant ECM were diluted in sterile 1X DPBS^++^ (Thermofisher 14040133, PBS containing 100mg/L Ca^2+^, 100 mg/L Mg^2+^) and used at the following concentration to coat tissue culture plates: Fibronectin (Sigma ECM001 1.6 µg/cm^2^), Collagen (Sigma C2249 1.8 µg/cm^2^), Laminin-111 (Biolamina LN111-0501, 1.6 µg/cm^2^), Laminin-211 (Biolamina LN211-0501, 1.6 µg/cm^2^), Laminin-411 (Biolamina LN411-0501, 1.6 µg/cm^2^) and Laminin-511 (Biolamina LN511-0502, 1.6 µg/cm^2^). Coating was allowed to proceed at 4°C o/n. Cells were treated with accutase (Accutase, ThermoFisher) before the excess coating solution was aspirated off the tissue culture plates, ensuring that the surface did not dry, and cells plated onto the surface in full serum medium.

### Senescence evaluation

Levels of senescence in Growing and G0cells were evaluated by treating cells under G0 or growing conditions with either solvent control, 50 nM doxorubicin for 4 days or 500 µM H_2_O_2_ for 2 hr followed by fixation and staining with senescence reporters. A Senescence β-Galactosidase Staining Kit (Cell Signaling 9860) was used according to the manufacturer’s instructions and the resultant signal was acquired on an ImageXpress microscope and quantified using automated image analysis in cell profiler. In addition, immunostaining against nuclear p21^WAF1/CIP1^ (12D1) (Cell Signaling Technology 2947) and nuclear phosphor-Histone H2A.X (Ser139) (Cell Signaling Technology 2947) were undertaken and imaged on an ImageXpress microscope with the resultant signal quantified using CellProfiler image analysis software.

### Trypan Blue Cell Viability Assay

A Trypan Blue Stock Solution of 0.4% was added to media in a 1:1 vol:vol as indicated in the manufacturers protocol. This mixture was then added to the indicated culture conditions of RPE1 cells for 5 minutes before being examined under a standard transmitted light microscope. At least 5 fields of view per condition at a 20x magnification were counted with all trypan blue positive and negative cells being recorded. Results are express as the percentage trypan blue positive cells, blue, from total cells normalized to 100% per condition. Triplicate biological repeats were undertaken.

### Caspase 3/7 apoptosis assay

Caspase 3/7 activity in cells was assessed using Magic Red fluorescent kit (Bio-Rad AbD Serotec ICT936) added to live cells at half the concentration indicated per manufacturer’s instructions and fluorescent signal was allowed to develop at 37°C +5% CO_2_ for 4 hr after which cells were fixed in 4% PFA and co-stained with DAPI nuclear label. Cells were then imaged on an ImageXpress microscope with the resultant signal quantified using CellProfiler image analysis software or FlowJo 8.8.6 (FlowJo LLC), as appropriate.

### Cell adhesion, survival and proliferation assays using xCELLigence RTCA DP systems

The degree of adhesion, proliferation or cells death was monitored in real time using the xCELLigence RTCA DP E-plate system (ACEA, CA, USA). 5 × 10^3^ cells, either integrin antibody block or untreated as indicated were seeded into each well coated with indicated matrix composition as above. The impedance value of each well were measured by the xCELLigence system every 30 seconds for the first 10 min for 72 h and expressed as a cell index value (CI). Wells were analyzed in duplicate with 3 independent biological repeats undertaken. Data is displayed as mean cell index over three experiments +/− SEM or as the mean cell index between duplicates across time for an individual experiment, +/- SD.

### Small compound inhibitors and ligands

The following small compound inhibitors (amongst the best-reported inhibitors for each kinase^47^) were used: GDC-0879 (called BRAFi in this study, Tocris 4453) used at 2.5 μM, ZM 336372 (called CRAFi in this study, Santa Cruz sc-200639) used at 2 μM, PD0325901 (called MEKi in this study, Tocris 4192) used at 5 μM, SCH772984 (called ERKi in this study, Selleckhem S7101) used at 5 μM, Staurosporin (Sigma Aldrich 19-123-MG) used at 2 μM, Doxorubicin (called Dox in this study, LKT Labs D5794) used at 50 nM, hydrogen peroxide (H2O2, Sigma Aldrich H1009) used at 0.4 mM, *Human* Epidermal Growth Factor (EGF, Sigma Aldrich E9644) used at 50 ng/mL, *Human* Tumour Necrosis Factor alpha (TNFα, Millipore GF442) used at 20 ng/mL, *Human* Transforming Growth Factor beta1 (TGFβ, R&D 240-B-002/CF) used at 5 ng/mL, and *Human* Bone Morphogenetic protein 2 (BMP2, R&D 355-BEC) used at 10 ng/mL.

### siRNA suppression of gene expression in growing and quiescent cells

The following siRNA oligos were used: 14-3-3ζ (Thermo Fisher Scientific Stealth HSS107094; 2 oligos against *human YWHAZ*), IQGAP-1 (Thermo Fisher Scientific Stealth HSS137296; 2 oligos against *human IQGAP1*), Integrin α3 (Thermo Fisher Scientific Stealth HSS107726; 2 oligos against *human ITGA3*) and Integrin α5 (Thermo Fisher Scientific Stealth HSS106727; 2 oligos against *human ITGA5*). Control siRNA used were Invitrogen Stealth control (scrambled) oligo 138782.

Exponentially growing cells seeded at 7x 10^3^ cells/cm^2^ in 96 well plates (Greiner Bio-One 655892), were transfected twice (on day 1 and 2, each followed by medium exchanges 24 h later) with Lipofectamine RNAi MAX (Thermo Fisher Scientific 13778150, 0.125 μL per well) complexed for 20 min at room temperature with each indicated siRNA (1.35 pmol per well) diluted in OptiMEM (Thermo Fisher Scientific 31985062, 5 μL per well) and analyzed 3 to 4 days after the first transfection.

Quiescent cells were seeded at 1.4x 10^4^ cells/cm^2^ in 96 well plates (Greiner Bio-One 655892), transfected twice (on day 5 and 7, each followed by medium exchanges 24 h later) with Lipofectamine RNAi MAX (0.425 μL per well) complexed for 20 min at room temperature with each indicated siRNA (2 pmol per well) diluted in OptiMEM (5 μL per well) and analyzed 10 to 12 days after the first transfection. RNAi knock-down efficiency was verified by immunofluorescence counter-staining. The use of validated pools of siRNA targeting the same genes increased the knock-down efficiency and specificity.

### Antibodies

The following primary antibodies were used for immunoblotting (IB), immunoprecipitation (IP) or immunostaining (IF, both immunofluorescence and flow cytometry): anti-Cyclin D1 (*rabbit* monoclonal 92G2, Cell Signaling Technology 2978) used at 1:1,000 dilution for IB, 1:100 for IF, anti-PCNA (*mouse* monoclonal PC10, Cell Signaling Technology 2586) used at 1:2,000 dilution, anti-Ki67 (*rabbit* polyclonal, AbCam ab15580) used at 0.5 μg/mL for IB, anti-phosphorylated Ser807/811 retinoblastoma, called ‘pRb’ in this study (*rabbit* monoclonal D20B12, Cell Signaling Technology 8516) used at 1μg/mL (1:400 dilution), anti-phosphorylated Ser139 Histone H2A.X, called ‘p-γH2AX’ in this study (*rabbit* monoclonal 20E3, Cell Signaling Technology 97186) used at 1:100 dilution, anti-p21^Waf1/Cip1^, called ‘p21’ in this study (*rabbit* monoclonal 12D1, Cell Signaling Technology 2947) used at 1:100 dilution, anti-p27 Kip1, called ‘p27’ in this study (*rabbit* monoclonal D69C12, Cell Signaling Technology 3686 and *mouse* monoclonal SX53G8.5, Cell Signaling Technology 3698) used at 1:100 dilution and 1:1,000 dilution, respectively, anti-Lgr5-647 (*rat* monoclonal 4D11F8 Alexa Fluor 647, BD Biosciences BD-562912) used at 1:100 dilution, anti-Caspase-3 (*rabbit* polyclonal, Cell Signaling Technology 9662) used at 1:1,000 dilution, anti-Caspase-7 (*rabbit* polyclonal, Cell Signaling Technology 9492) used at 1:1,000 dilution), anti-Caspase-9 (*rabbit* polyclonal, Cell Signaling Technology 9502) used at 1:1,000 dilution, anti-Bim (*rabbit* monoclonal C34C5, Cell Signaling Technology 9502) used at 1:1000 dilution, anti-BID (*rabbit* polyclonal, Cell Signaling Technology 2002) used at 1:1,000 dilution, anti-Bik (*rabbit* polyclonal, Cell Signaling Technology 4592) used at 1:1,000 dilution, anti-Bax (*rabbit* monoclonal D2E11, Cell Signaling Technology 5023) used 1:1,000 dilution), anti-Bak (*rabbit* monoclonal D2D3, Cell Signaling Technology 6947) used at 1:1,000 dilution, anti-Puma (*rabbit* polyclonal, Cell Signaling Technology 4976) used at 1:1,000 dilution, anti-BAD (*rabbit* monoclonal D24A9, Cell Signaling Technology 9239) used at 1:1,000 dilution, anti-phosphorylated Ser112 BAD, called ‘pBAD’ in this study (*rabbit* monoclonal 40A9, Cell Signaling Technology 5284) used at 1:100 dilution, anti-Integrin α3 (*mouse* monoclonal J1432, GeneTex GTX11767 and *mouse* monoclonal P1B5, Santa Cruz sc-13545) used 1:400 and 1:50 dilution (blocking experiemnts), respectively, anti-Integrin α4 (*rabbit* polyclonal, Cell Signaling Technology 4600) used at 1:100 dilution), anti-Integrin α5 (*rabbit* polyclonal, Cell Signaling Technology 4705 or *mouse* monoclonal P1D6, Santa Cruz sc-13547) used at 1:100 dilution, anti-Integrin α11 (*mouse* monoclonal F5, Santa Cruz sc-390091) used at 1:100 dilution, anti-Integrin αV (*rabbit* polyclonal, Cell Signaling Technology 4711) used at 1:1,000 dilution, anti-Integrin β1 (*rabbit* polyclonal, Cell Signaling Technology 4706 or *rat* monoclonal 9EG7, BD Pharmigen 553715) used at 1:1,000 dilution, anti-Integrin β3 (*rabbit* polyclonal, Cell Signaling Technology 4702) used at 1:1,000 dilution, anti-Integrin β4 (*mouse* monoclonal 3E1, Millipore MAB1964 or *rabbit* polyclonal, Cell Signaling Technology 4707) used at 1:1,000 dilution, anti-Integrin β5 (*rabbit* polyclonal, Cell Signaling Technology 4708) used at 1:1,000 dilution, anti-CD151 (*mouse* monoclonal 50-6, Biolegend 350402) used at 1:100 dilution, anti-AKT (*rabbit* polyclonal, Cell Signaling Technology 9272) used at 1:100 dilution, anti-phosphorylated Ser473 AKT, called ‘pAKT’ in this study (*rabbit* monoclonal D9E, Cell Signaling Technology 4060) used at 1:100 dilution, anti-PRAS40 (*rabbit* monoclonal D23C7, Cell Signaling Technology 2691) used at 1:1,000 dilution, anti-phosphorylated Thr246 PRAS40, called ‘pPRAS40’ in this study (*rabbit* monoclonal C77D7, Cell Signaling Technology 2997) used at 1:800 dilution, anti-p70 S6 Kinase, called ‘p70S6K’ in this study (*rabbit* monoclonal 49D7, Cell Signaling Technology 2708) used at 1:1,000 dilution, anti-phosphorylated Ser371 p70 S6 Kinase, called ‘p-p70S6K’ in this study (*rabbit* polyclonal, Cell Signaling Technology 9208) used at 1:1,000 dilution), anti-phosphorylated Thr389 p70 S6 Kinase, called ‘p-p70S6K’ in this study (*rabbit* monoclonal 108D2, Cell Signaling Technology 9234) used at 1:1,000 dilution), anti-4E-BP1 (*rabbit* monoclonal 53H11, Cell Signaling Technology 9644 or *rabbit* polyclonal, Cell Signaling Technology 2845) used at 1:1,000 dilution, anti-phosphorylated Thr37/46 4E-BP1, called ‘p4EBP Thr37/46’ in this study (*rabbit* monoclonal 236B4, Cell Signaling Technology 2855) used at 1:1,000 dilution, anti-phosphorylated Thr70 4E-BP1, called ‘p4EBP Thr70’ in this study (*rabbit* polyclonal, Cell Signaling Technology 9455) used at 1:1,000 dilution, anti-LC3B (*rabbit* monoclonal D11, Cell Signaling Technology 3868) used at X μg/mL (1:1,000 dilution), anti-A-Raf, called ‘ARAF in this study (*rabbit* polyclonal, Cell Signaling Technology 4432) used at 1:1,000 dilution, anti-phosphorylated Ser299 A-Raf, called ‘pARAF’ in this study (*rabbit* polyclonal, Cell Signaling Technology 4431) used at 1:1,000 dilution, anti-B-Raf, called ‘BRAF in this study (*rabbit* monoclonal 55C6, Cell Signaling Technology 9433) used at 1:1,000 dilution, anti-phosphorylated Ser445 B-Raf, called ‘pBRAF’ in this study (*rabbit* polyclonal, Cell Signaling Technology 2696) used at 1:1,000 dilution, anti-c-Raf, called ‘CRAF in this study (*mouse* monoclonal D5X6R, Cell Signaling Technology 12552) used at 1:1,000 dilution), anti-phosphorylated Ser338 c-Raf, called ‘pCRAF’ in this study (*rabbit* monoclonal 56A6, Cell Signaling Technology 9427) used at 1:1,000 dilution, anti-MEK1/2, called ‘MEK’ in this study (*mouse* monoclonal L38C12, Cell Signaling Technology 4694) used at 1:1,000 dilution, anti-phosphorylated Ser217/221 MEK1/2, called ‘pMEK’ in this study (*rabbit* monoclonal 41G9, Cell Signaling Technology 9154) used at 1:1000 dilution, anti-p44/42 MAPK (ERK1/2), called ‘ERK’ in this study (*rabbit* polyclonal, Cell Signaling Technology 9102 and *mouse* monoclonal MK1, Santa Cruz sc-135900) used at 1:1,000 dilution, anti-phosphorylated Thr202/Tyr204 p44/42 MAPK (ERK1/2), called ‘pERK’ in this study (*rabbit* monoclonal D13.14.4E, Cell Signaling Technology 4370 and *mouse* monoclonal pT202/pY204.22A, Santa Cruz sc-136521) used at 1:100 dilution, anti-phosphorylated Ser73 c-Jun, called ‘p-cJun’ in this study (*rabbit* monoclonal D47G9, Cell Signaling Technology 3270) used at 1:600 dilution, anti-phosphorylated Ser32 c-Fos, called ‘p-cFos’ in this study (*rabbit* monoclonal D82C12, Cell Signaling Technology 5348) used at 1:400 dilution, anti-phosphorylated Ser133 CREB, called ‘pCREB’ in this study (*rabbit* monoclonal 87G3, Cell Signaling Technology 9198) used at 1:600 dilution, anti-phosphorylated Thr581 MSK1, called ‘pMSK1’ in this study (*rabbit* polyclonal, Cell Signaling Technology 9595) used at 1:1,000 dilution), anti-phosphorylated S383 Elk1, called ‘pElk1’ in this study (*rabbit* monoclonal, Cell Signaling Technology 9181) used at:100 dilution, anti- AMPKα, called ‘AMPK’ in this study (*rabbit* polyclonal, Cell Signaling Technology 2532) used at 1:1,000 dilution, anti-phosphorylated Thr172 AMPKα, called ‘pAMPK’ in this study (*rabbit* monoclonal 40H9, Cell Signaling Technology 2535) used at 1:1,000 dilution, anti-p38 MAPK (*rabbit* polyclonal, Cell Signaling Technology 9212) used at 1:1,000 dilution, anti-phosphorylated Thr180/Tyr182 p38 MAKP, called ‘p-p38 MAPK’ in this study (*rabbit* monoclonal D3F9, Cell Signaling Technology 4511) used at 1:1,000 dilution), anti-SAPK/JNK (JNK1, JNK2 and JNK3), called ‘pJNK’ in this study (*rabbit* polyclonal, Cell Signaling Technology 9252) used at 1:1,000 dilution), anti-phosphorylated Thr183/Tyr185 p46/p54 SAPK/JNK, called ‘pJNK’ in this study (*rabbit* monoclonal G9, Cell Signaling Technology 9255) used at 1:2,000 dilution, anti-Mst1/2/STK3-4 called ‘Mst1/2’ in this study (*rabbit* polyclonal, Bethyl Laboratories A300-466A) used at 1:1000 dilution, anti-phosphorylated Thr183/Thr180 Mst1/2, called ‘pMst1/2’ in this study (*rabbit* polyclonal, Cell Signaling Technology 3681) used at 1:1,000 dilution), anti- YAP1, (*mouse* monoclonal 63.7, Santa Cruz sc-101199) used at 1:500 dilution, anti-phosphorylated Ser127 YAP1, called ‘pYAP1’ in this study (*rabbit* monoclonal D9W2I, Cell Signaling Technology 13008) used at 1:1,000 dilution, anti-SMAD1, (*rabbit* monoclonal D59D7, Cell Signaling Technology 6944) used at 1:1,000 dilution, anti-phosphorylated Ser463/465 SMAD1, Ser463/465 SMAD5, Ser465/467 SMAD9 (aka SMAD8) called ‘pSMAD1/5/8’ in this study (*rabbit* monoclonal D5B10, Cell Signaling Technology 13820) used at 1:200 dilution, anti- SMAD2/SMAD3, (*rabbit* monoclonal D7G7, Cell Signaling Technology 8685) used at 1:100 dilution, anti-phosphorylated Ser465/467 SMAD2, called ‘pSMAD2’ in this study (*rabbit* monoclonal 138D4, Cell Signaling Technology 3108) used at 1:1,000 dilution), anti-SMAD4 (*rabbit* polyclonal, Cell Signaling Technology 9515) used at 1:400 dilution, anti-NF-κB p65, called ‘p65’ in this study (*rabbit* monoclonal D14E12, Cell Signaling Technology 8242) used at 1:1,000 dilution), anti- NF-κB p52/p100, called ‘p52/p100’ in this study (*rabbit* monoclonal 18D10, Cell Signaling Technology 3017) used at 1:1,000 dilution, anti-RelB (*rabbit* monoclonal C1E4, Cell Signaling Technology 4922) used at 1:1,000 dilution, anti-Notch 1, called ‘Notch’ in this study (*rabbit* monoclonal D6F11, Cell Signaling Technology 4380) used at 1:1,000 dilution, anti-cleaved Notch1 (Val1744), called ‘NICD’ in this study (*rabbit* monoclonal D3B8, Cell Signaling Technology 4147) used at 1:1,000 dilution, anti-GLI1 (*rabbit* monoclonal C68H3, Cell Signaling Technology 3538) used at 1:1,000 dilution, anti-acetylated Tubulin, (*mouse* monoclonal 6- 11B1, Sigma-Aldrich T7451), anti-αTubulin (*mouse* monoclonal TUB2.1, AbCam ab11308) used at 1:2,000 dilution, anti-GSKα/β (*rabbit* monoclonal D75D3, Cell Signaling Technology 5676) used at 1:400 dilution, anti-phosphorylated Ser9 GSK3β, called ‘pGSK3’ in this study (*rabbit* monoclonal D85E12, Cell Signaling Technology 5558) used at 1:400 dilution, anti-β-catenin (*rabbit* monoclonal D10A8, Cell Signaling Technology 8480) used at 1:1,000 dilution, anti-14-3-3ζ (*rabbit* polyclonal C-16, Santa Cruz sc-1019) used at 1:100 dilution, anti-IQGAP1 (*rabbit* polyclonal, Bethyl Laboratories A301-950A) used at 1:100 dilution, and anti-GAPDH (*mouse* monoclonal 0411, Santa Cruz sc-47724 and *rabbit* monoclonal 14C10, Cell Signaling Technology 2118) used at 0.007μg/mL (IB) or 1:2,000 dilution (IF).The following secondary antibodies were used for flow cytometry (all used at 1:1,000 dilution) and fluorescent microscopy (all used at 1:500 dilution): AlexaFluor 488 *goat* anti-*mouse* IgG (Thermo Fisher Scientific A-11001), AlexaFluor 488 *goat* anti-*rabbit* IgG, (Thermo Fisher Scientific A-11008), AlexaFluor 488 *goat* anti-*rat* IgG, (Thermo Fisher Scientific A-11006), AlexaFluor 555 *goat* anti-*mouse* IgG, (Thermo Fisher Scientific A-21422), AlexaFluor 555 *donkey* anti-*mouse* IgG (Thermo Fisher Scientific A-31570), AlexaFluor 555 *goat* anti-*rabbit* IgG, (Thermo Fisher Scientific A-21428), AlexaFluor 594 *donkey* anti-*mouse* IgG (Thermo Fisher Scientific A-21203), AlexaFluor 633 *goat* anti-*mouse* IgG, (Thermo Fisher Scientific A-21050), AlexaFluor 633 *goat* anti- *rabbit* IgG, (Thermo Fisher Scientific A-21070), AlexaFluor 647 *chicken* anti-*mouse* IgG, (Thermo Fisher Scientific A-21463), AlexaFluor 647 *chicken* anti-*rat* IgG, (Thermo Fisher Scientific A-21472), AlexaFluor 647 *chicken* anti-*rabbit* IgG, (Thermo Fisher Scientific A-21443), and for immunoblot (1:10,000 dilution): *goat* anti- *mouse* IgG-HRP conjugated (Bio-Rad 1706516) and *goat* anti-*rabbit* IgG-HRP conjugated (Bio-Rad 1706519). Actin was stained using Phalloidin-Alexa647 (Cell Signaling Technology 8940) and DNA using 2.5μg/ml Hoechst 33342 (Thermo Fisher Scientific H3570), 2 μg/ml 4’,6-diamidino-2-phenylindole (DAPI, Sigma-Aldrich D9542) or DRAQ5 (BioStatus DR50200).

### Immunostaining for flow cytometry

Cells were pelleted between each following steps for 5 min at 1,000 *g*. After assay, cells were resuspended in ice-cold PBS, fixed in 4 % PFA (Thermo Fisher Scientific 28908) at room temperature for 20 min, washed twice for at least 30 min with PBS with 50 mM NH4Cl to quench free PFA, and stored at 4 °C, or used immediately for immunostaining. Depending on antibodies, cells were immunostained for 90 min at room temperature (with gentle mixing) with primary antibodies diluted in PBS with 0.05-0.1% saponin (Sigma-Aldrich 47036) or 0.1% Triton X-100 (Sigma-Aldrich 9036) and 5-10% heat inactivated Horse Serum or BSA, followed by secondary antibodies and 2.5μg/ml Hoechst 33342 or 2 μg/ml DAPI. Subsequent to secondary antibody staining, cells were washed three times for 5 min in PBS with 0.05% saponin, two times in PBS, passed through a nylon mesh filter to eliminate clumps and either used immediately or stored at 4 °C in the dark until flow cytometry analysis.

### Cell sorting and flow cytometry

Growing and quiescent cell populations were harvested with Accutase, their DNA live-stained with 2.5 μg/ml Hoechst 33342 for 30-45 min at 37°C, washed with ice-cold PBS^++^ and G1 and G0 cells with 2n DNA were sorted using a MoFlo XDP Cell Sorter (Beckman Coulter) and used for further experiments (mass spectrometry, immunoblotting or labelling for flow cytometry analysis).

Flow cytometry analysis of fixed and immunolabeled cell samples was performed on a BD Biosciences LSRII equipped with a 325 nm ultraviolet laser and 450/50 nm band-pass filter for Hoechst or DAPI, a 488 nm laser and 530/30 nm bandpass filter for Alex-aFluor488 or DiO, and a 633 nm laser and 660/20 nm bandpass filter for AlexaFluor633 or -647. For experiments requiring red fluorescent dyes (AlexaFluor555 or DiI), a BD Biosciences FACS AriaIII with 561nm excitation and 610/20nm emission bandpass filter was used instead.

Data were analysed using FlowJo 8.8.6 (FlowJo LLC). To compare signals in proliferating and quiescent cells, laser intensity was kept constant when analysing the two cell cycle states. In a height versus area dot plot of DNA stain fluorescence intensity (Hoechst or DAPI), a gate was created on single cells to eliminate cell aggregates from further analysis. A DNA profile histogram of DNA stain fluorescence intensity was then used to gate on the cell population with 2n DNA content (G1 or G0 populations). The average geometrical mean of the signal area was used for quantification. Signal of labelled cells was divided by signal of control cells labelled with unspecific IgG or blank cells for activity assays and expressed as x-fold signal intensity of background signal.

### Cell lysis and protein quantification

Quiescent RPE1 cells were harvested with Accutase; proliferating RPE1 cells grown in full cell culture medium were harvested with Accutase, labelled with Hoechst 33342 and sorted for the G1 (2n DNA) population by flow cytometry (MoFlo XDP cell sorter). Both cell population were lysed in Lysis Buffer (8M urea (Sigma-Aldrich U1250) containing phosphatase (Cell Signaling Technology 5870S) and protease inhibitors (Cell Signaling Technology 5871S), to a final protein concentration of *c.* 25 mg/mL. For quantification of protein levels per cell, 10^6^ growing or quiescent cells were sorted by flow cytometry, lysed in 100 μl of Lysis buffer (8M urea with phosphatase and protease inhibitors). Following lysis, samples were sonicated (five times 5-15 second ultrasound pulses with 30 seconds rest on ice, 7 μm amplitude) to fragment DNA.

### Protein quantification by Bradford assay

Total protein content per cell in proliferating and quiescent cell populations were measured using Coomassie Plus (Bradford) Protein Assay kit (Thermo Fisher Scientific 23236). The assay was performed according to manufacturer’s instructions. Briefly, a bovine serum albumin (BSA) standard was created according to manufacturer’s instructions diluting BSA ranging from 0 to 2000 μg/ml protein in Lysis Buffer (8M urea containing phosphatase and protease inhibitors). Protein lysates were diluted in Lysis Buffer to concentrations within the linear range of the BSA standard curve. Triplets of each sample were analysed in a 96- well microtiter plate measuring absorbance at 594nm using FLUOstar OPTIMA ABS absorbance microplate reader (BMG LABTECH). A linear regression function was fitted on the BSA absorbance data points in the linear range from 125-1,000 μg/ml. Protein concentration of samples was determined using the mean of technical triplicates which were within the linear range of the BSA standard curve.

### Protein quantification by SDS-PAGE and Coomassie staining

Sonicated cell lysates were boiled (95 °C for 10 min) in SDS sample buffer (50 mM Tris pH 6.8, 10% (v/v) SDS, 500 mM DTT, 0.05% (v/v) Bromophenol Blue and 45% (v/v) glycerol) and ran on 12% Bis-Tris Plus SDS gels (Thermo Fisher Scientific NW00120BOX). Gels were stained (1 hour, room temperature) with Coomassie staining solution (45% (v/v) Methanol, 10% (v/v) glacial acetic acid, 0.3% (v/v) Brilliant Blue G250) and destained for 1 hour with Coomassie destaining solution (40% (v/v) Ethanol and 10% (v/v) glacial acetic acid) and further destained in 1:1 (v/v) Coomassie destaining solution/dH2O overnight. Gels were scanned and protein levels were quantified by densitometry quantification of lanes in 8-bit images using ImageJ software and compared relative to each other.

### Co-immunoprecipitations

To generate G0 cells, RPE1 cells were seeded at a 1:10 ratio in six 150 mm round plates containing and cultured in full serum media, with media replaced every two days. After ten days, ensuring minimal mitotic activity, the media was switched to serum-free, again replacing every two days. On day 17, growing cells were seeded at a 1:10 ratio in eighteen 150 mm round plates in full serum media. For immunoprecipitation, antibody (Raf-B Antibody (F-7), sc-5284 Santa Cruz or IgG Isotype control, Cell Signalling 61656) and beads (Dynabeads Protein G) were incubated overnight at 4°C in a bead:antibody volume ratio of 2:1 (20 μl:10 μl) in 2ml of TBS. Tubes were placed on a rotating wheel overnight at 4°C. On day 19, both growing and G0 cells were washed twice in 1X PBS and drained to remove excess liquid. For G0 and growing cells, 700 μl per plate of IP buffer (50 mM Tris-HCl pH 7.5, 150 mM NaCl, 1% IGEPAL, 1X protease (cOmplete, Mini, EDTA-free Protease Inhibitor Cocktail, Roche) and phosphatase inhibitors (PhosStop, Roche)) were added, and cells were scraped and pooled into a 15 ml Falcon tube and incubated on ice for 30 minutes. Lysates were sonicated for 15 seconds at 40% amplitude and centrifuged at 14,000 x g for 10 minutes at 4°C. Supernatants were transferred to fresh tubes and pre-cleared with 30 μl of protein G Dynabeads alone for 2 hours at 4°C before centrifugation and collection of the supernatant. A protein concentration assay (Pierce BCA Protein Assay Kits) was performed to calculate the amount of lysate needed to equate to 5500 μg of protein per condition. 60 μg of each condition were taken to serve as input controls. Antibody-bound Dynabeads were washed with TBS, and appropriate volumes of lysate were added to each tube (BRAF G0, BRAF Growing, Isotype Ab G0, Isotype Ab Growing) and the resultant mixtures were rotated overnight at 4°C. Beads were were then washed five times in excess IP buffer at 4°C, before resuspended in 30 μl of 2X LDL gel-loading buffer + 5% βME. Samples were treated with nuclease (Benzonase Nuclease, Millipore) and heated at 95°C for 5 minutes, cooled, and centrifuged before loading 15 μl of supernatant per well, with 30 μg of input controls for G0 and Growing being loaded per well as appropriate. SDS-PAGE Gels were run at 120 V for 10 minutes and then at 150 V until completion. For transfer, PVDF membranes were activated in methanol, and gels were transferred in transfer buffer (48 mM Tris Base, 390 mM Glycine, 0.1% w/v SDS, 20% methanol) at 95 V in a cold room for 90 minutes for BRAF and pERK, and 120 minutes for IQGAP1. Membranes were blocked with fluorescent block (Immobilon Block Millipore) for 1 hour, incubated overnight with primary as appropriate (1:1000 in 5% BSA TBS/Tween-20) at 4°C, washed, and incubated with fluorescent secondary antibodies in 5% milk TBS/T for 1 hour at room temperature. Membranes were washed and imaged using LiCOR Odyssey CLx Imaging system.

### GTP-Ras and GTP-Rap1 detection

Active Ras Detection kit (Cell Signaling Technology 8821) was used following the manufacturer’s instructions. Briefly, growing or G0 cell lysates (1mg/mL) were treated *in vitro* with GTPγs or GDP to activate or inactivate Ras and then incubated with glutathione resin and GST-Raf1-RBD. GTPγS-treated lysate was also incubated without GST-Raf1-RBD in the presence of glutathione resin as a negative control. Active Rap1 Detection kit (Cell Signaling Technology 8818) was used following the manufacturer’s instructions. Briefly, growing or G0 cell lysates (1mg/mL) were treated *in vitro* with GTPγs or GDP to activate or inactivate Rap1 and then incubated with glutathione resin and GST-RalGDS-RBD. GTPγS-treated lysate was also incubated without GST-RalGDS-RBD in the presence of glutathione resin as a negative control. Immunoblot analysis of cell lysates or eluted samples were performed using a *mouse* monoclonal anti-Ras, a *rabbit* monoclonal anti-Rap1 or a *mouse* monoclonal anti-GST antibodies.

### SDS-PAGE and protein transfer

Sonicated cell lysates were boiled (95 °C for 10 min) in SDS sample buffer (50 mM Tris pH 6.8, 10% (v/v) SDS, 100 mM DTT, 0.05% (v/v) Bromophenol Blue and 45% (v/v) glycerol) and ran on 12% Bis-Tris Plus SDS acrylamide gels (Thermo Fisher Scientific NW00120). Depending on the molecular weights and abundances of the proteins of interest, 20-100μg protein were loaded per lane on 8 to 12 % Bis-Tris Plus SDS acrylamide, ran with MES or MOPS running buffers in a Bolt Mini Gel Tank (all Thermo Fisher Scientific) at 300 V and 75 mA for 30-45 min. Proteins were transferred onto PVDF or nitrocellulose membranes via semi-dry transfer blotter (Perkin Elmer NEF201001EA) with fixed voltage at 24 V and variable current for 20-45 min. Membranes were washed twice in TBST (50 mM Tris pH 7.4, 150 mM NaCl, 0.05 % (v/v) Tween20) and blocked in 5 % skimmed milk (Thermo Fisher Scientific LP0031) in TBST for one hour. Membranes were incubated for either 2 hours at room temperature or overnight at 4 °C, with primary antibodies diluted in TBST containing 5% horse serum, were washed incubated for 30 min with horseradish peroxidase (HRP)-conjugated secondary antibodies (BioRad) diluted in TBST containing 5 % horse serum, and finally washed another three times 10 min in TBST. Blots were developed with the ECL kit (Thermo Fischer Scientific or Merck Millipore) and x-ray films Amersham Hyperfilm ECL (GE Healthcare 289068367), developed in a Compact X4 film developer (Xograph Imaging Systems). In some experiments, membranes were stripped twice for 15 min each in pre-warmed Immunoblot stripping buffer 200 mM Glycine, 0.01% (v/v) SDS, 1% (v/v) Tween-20, pH 2.5), washed twice in PBS for 10 min and twice in TBST for 5 min before being re-probed with another primary antibody. Uncropped and unprocessed immunoblots are shown in Supplementary Figures S16 to 18.

### SILAC proteomics and phosphoproteomics

Exponentially growing or quiescent RPE1 cells were cultured in light or heavy Stable isotope labeling by amino acids in culture (SILAC) medium (DMEM/F12 SILAC (-K/-R), Thermo Fisher Scientific Pierce 88215), supplemented with 40 mg/L K8 (Sigma 608041), 84 mg/L R10 (Sigma 608033), 200 mg/L L-Proline (to avoid arginine-to-proline conversion^48^ ), 10% FBS (dialyzed three times 3 hours in 120 mM NaCl, 32 mM NaHCO_3_ at 3.5 mW cutoff), 5% Sodium bicarbonate w/v (Sigma Aldrich S8761), 1mM GlutaMAX-I (Thermo Fisher Scientific A1286001), 1X antibiotic-antimycotic (Thermo Fisher Scientific 15240062),) for at least five cell divisions to allow complete labelling. Light- and Heavy-SILAC quiescent and growing cell samples were harvested with Accutase, labelled with Hoechst 33342 and sorted for the G0 and G1 (2n DNA) populations by flow cytometry (MoFlo XDP cell sorter) and lysed in Lysis Buffer (8M urea, Sigma-Aldrich U1250) containing phosphatase (Cell Signaling Technology 5870S) and protease inhibitors (Cell Signaling Technology 5871S).

*SILAC proteomics.* G0 and G1 lysates from >3 biological repeats were combined, G0/G1 and G1/G0 (Heavy/Light) mixes (50μg of each) were separated by 4-12% SDS-PAGE ran in MES buffer, lanes cut in 20 horizontal bands and each band was further cut into 1 mm cube. Gel pieces were reduced with DTT, alkylated with IAA and digested with trypsin at 37°C overnight. Peptides were extracted from gel pieces and partially dried in a SpeedVac concentrator (Savant). These extracts were desalted with C18 stage tips (3M Empore) prior to mass spectrometry analysis.

*SILAC phosphoproteomics. Enzymatic digestion.* Protein samples (G0 and G1 lysates from >3 biological repeats were combined) in 8M urea, 20mM Hepes were reduced with 5 mM DTT at 56 °C for 30 min and alkylated with 10 mM iodoacetamide (IAA) in the dark at room temperature for 30 min. Lys_C (Promega) was then added to the samples and digested for 3 hr at 25 °C. Next, urea concentration was diluted to 2M and samples were incubated with trypsin (Promega) over night, at 25°C. Digestion was stopped by the addition of trifluoroacetic acid (TFA) to a final concentration of 1 % and then centrifuged at 4000*g* for 20min. The supernatants were desalted using Sep-Pak Plus Short tC18 cartridges (Waters). Bound peptides were eluted with 60% acetonitrile in 0.5% acetic acid and lyophilized.

*Strong Cation Exchange Chromatography.* Strong cation exchange (SCX) chromatography was performed using an Ultimate 3000 Nano/Capillary LC System (Dionex). Lyophilized peptides were resolubilized in 200 μL of solvent A (5 mM KHPO_4_ pH 2.65, 30% MeCN) and loaded onto a PolySULFOETHYL A column, 4.6 mm x 204 mm (PolyLC). The peptides were separated using a 45 min linear gradient of 0-23% solvent B (solvent A + 350 mM KCl) at a flow rate of 1 ml/min. Fifteen fractions, in 3 min intervals, were collected and lyophilized. Each fraction was resuspended in 0.1% TFA and desalted using home-made Poros R3 reversed phase (Applied Biosystems) column. The eluted peptides were lyophilized prior to TiO2 enrichment.

*TiO_2_ enrichment of Phosphopeptides.* Lyophilized peptides were resuspended in a solution of 2, 4- dihydroxybenzoic acid, 80% MeCN, 2% TFA (loading buffer) and incubated with TiO_2 beads_ (GL Science Inc. Japan) for 15 min. Beads were transferred to a C8 stage tip (P10) and washed sequentially with loading buffer and 80% MeCN, 2%TFA. Phosphopeptides were eluted with ammonia solution, pH10.5, followed by 50% MeCN, 0.05% TFA. The combined eluates were partially dried down and desalted with a C18 Stage tip (3M Empore).

*Mass Spectrometry Data Acquisition.* Liquid chromatography was performed on a fully automated Ultimate U3000 Nano LC System (Dionex) fitted with a 100µm x 2cm PepMap100 C_18_ nano trap column and a 75μm×25cm reverse phase PepMap100 C_18_ nano column (Dionex). Samples were separated using a binary gradient consisting of buffer A (2% MeCN, 0.1% formic acid) and buffer B (90% MeCN, 0.1% formic acid). The HPLC system was coupled to an LTQ Orbitrap Velos mass spectrometer (Thermo Scientific) equipped with a nanospray ion source. The mass spectrometer was operated in standard data dependent mode, performed survey full-scan (m/z = 320-1600) in the Orbitrap analyzer, followed by MS^2^ acquisitions of the 20 most intense ions in the LTQ ion trap and multistage activation was used for phosphoproteomics only. Dynamic exclusion was enabled with a repeat count of 1, exclusion list of 500 and exclusion duration of 60s.

*Mass Spectrometry analysis*. MS Raw file spectra was analyzed using MaxQuant search engine suite (version 1.6.1.0^49^, using default search settings for a SILAC labelled phosphoproteomics workflow and a minimum label ratio count of 2, unless otherwise noted were searched against the UniProtKB/SwissProt human proteome databases (03/04/2013), concatenated with the default Maxquant contaminant database. Carbamidomethylation of Cystein was set as a fixed modification, and oxidation of Methionine and acetylation of the protein amino- terminus and phodphorylation of STY were allowed as variable modifications. The resulting peptide-to-spectrum matches (PSMs) were filtered by MaxQuant at 1% FDR. We performed a standard Perseus workflow to evaluate quality and identify statistically robust differentially expressed (DE) proteins and phosphorylation site on log2 transformed data calculated by MaxQuant.

### Label-free mass spectrometry sample preparation by in-gel digestion

Samples were separated by SDS-PAGE on a 4-12% gel and stained with Instant Blue (Expedeon) staining solution. Sample lanes were cut into 10 sections, then further cut into 1mm^3^ pieces and washed in destaining solution (40% ethanol, 10% glacial acetic acid in water). Proteins were reduced in 10 mM DTT, then alkylated with 20 mM iodoacetamide in 50 mM ammonium bicarbonate buffers. Gel pieces were washed then immersed in a 10ng/µl Trypsin buffer in 50 mM ammonium bicarbonate and digested for 16-18 hours at 37°C. Gel pieces were incubated in Elution buffer (1% formic acid, 2% acetonitrile in LC-MS grade water (Thermo Fisher Scientific)) and dried in a SpeedVac. Peptides resuspended in 0.5% acetic acid in water and desalted on C18-Stagetips, then dried in a SpeedVac. Peptides were resuspended in LC-MS running buffer (3% acetonitrile, 0.1% Formic acid in water) prior to analysis by LC-MS and spiked with *E. coli* ClpB peptides (Waters, UK) such that 50 fmol of spiked-in peptide standard was introduced per injection.

### Liquid chromatography and label-free mass spectrometry data acquisition

LC-MS grade solvents were used for all chromatographic steps. Separation of peptides was performed using a Waters NanoAcquity Ultra-Performance Liquid Chromatography system. Peptides were reconstituted in 97:3 H2O:acetonitrile + 0.1% formic acid. The mobile phase was: A) H2O + 0.1% formic acid and B) Acetonitrile + 0.1% formic acid. Desalting of the samples was performed online using a reversed-phase C18 trapping column (180 µm internal diameter, 20 mm length, 5 µm particle size; Waters). Peptides were separated by a linear gradient (0.3 μl/min, 35°C column temperature; 97-60% Buffer A over 60 min) using an Acquity UPLC M-Class Peptide BEH C18 column (130Å pore size, 75 μm internal diameter, 250 mm length, 1.7 μm particle size, Waters, UK). [Glu1]-fibrinopeptide B (GFP, Waters, UK) was used as lockmass at 100 fmol/µl. Lockmass solution was delivered from an auxiliary pump operating at 0.5 µl/min to a reference sprayer sampled every 60 seconds. The nanoLC was coupled online through a nanoflow sprayer to a QToF hybrid mass spectrometer (Synapt G2-Si; Waters, UK). Accurate mass measurements were made using a data-independent mode of acquisition (HDMS^E^). Each sample was analyzed in technical duplicate.

### Database searching and MS analysis

Raw data was analyzed using Progenesis v4.0 (Waters, UK). Data were queried against a Homo sapiens FASTA protein database (UniProt proteome:UP000005640) concatenated with a list of common contaminants obtained from the Global Proteome Machine (ftp://ftp.thegpm.org/fasta/cRAP) and *E. coli* ClpB, which acted as a standard for label-free absolute protein quantitation^50^. Carbamidomethyl-C was specified as a fixed modification and Oxidation (M) and Phosphorylation of STY were specified as variable modifications. A maximum of 2 missed cleavages were tolerated in the analysis to account for incomplete digestion.

For peptide identification 3 corresponding fragment ions were set as a minimum criterion whereas for protein identification a minimum of 7 fragment ions were required. Protein false discovery rate was set at 1%. Samples were normalized to peptide abundance and fractions were combined *in-silico* in Progenesis to obtain absolute protein abundances for differential expression analysis.

### Proteomics Functional and interaction analysis

Downstream proteomic and phosphoproteomic analysis were undertaken by importing to Cytoscape (v3.7.1) for protein interaction network analysis using the STRING database plugin with interactions filtered based on confidence of the interaction with a medium confidence (score of ≥ 0.4). Overrepresentation enrichments analysis was employed with Fisher’s exact tests to calculate p value and the Benjamini–Hochberg procedure to calculate false discovery rates (FDR). FDR cutoffs of < 0.05 were utilized and the analysis of the genes identified to be up- and downregulated was first performed using the Bingo plugin (3.0.4). The Cytoscape plugin PhosphoPath was used to visualize and identify regulated phosphor-regulated pathways using protein-protein interactions from the biogrid database (BioGrid) and kinase-substrate interactions (PhosphositePlus) with pathway networks determined by (Wikipathways).

### Immunostaining for microscopy

Growing cells were seeded 2 days prior to the experiments at 1.5 x10^4^ cells/cm^2^ and quiescent cells were seeded 17 days prior to the experiments at 1.4 x10^4^ cells/cm^2^ either in glass-bottom 96- well microplates (Greiner Bio-One 655892) or on glass coverslips (0.13-0.16 mm (Academy) previously washed with acetone and 100 % ethanol and placed in 24-well plates. After assay, cells were fixed with 4% PFA (Thermo Fisher Scientific 28908) at 37 °C for 20 min, washed 3 times with PBS and 1 time for at least 30 min with PBS with 50 mM NH4Cl to quench free PFA. Depending on antibodies, cells were immunostained for 30-90 min at room temperature or overnight at 37 °C with primary antibodies diluted in PBS with 0.05-0.1% saponin (Sigma-Aldrich 47036) or 0.1% Triton X-100 (Sigma-Aldrich 9036) and 5-10% heat inactivated Horse Serum or BSA, followed by secondary antibodies and 3.3 μM Draq5 (Thermo Fisher Scientific 62251) or 2 μg/ml 4’,6-diamidino-2-phenylindole (DAPI) (Sigma-Aldrich D9542). Subsequent to secondary antibody staining, cells were washed three times for 5 min in PBS with 0.05% saponin, two times in PBS and two times in miliQ water. Water was exchanged for an immunomount Mowiol solution containing antifading agent 1,4-diazabicyclo[2.2.2]octane (DABCO) (GeneTex GTX30930) and the microplates were either imaged immediately or stored at room temperature in the dark over night before long-term storage at 4 °C. Cover slips were mounted on glass slides (Thermo Scientific) using immunomount Mowiol solution containing DABCO, kept at room temperature in the dark over night to solidify and then stored at 4 °C.

### Pig intestine sample preparation and immunostaining

Within 2 hours of death, porcine ileum samples were washed three times in PBS containing penicillin/streptomycin. Tubular sections of approximately 1 cm were cut and fixed at room temperature for 1 hour with 4% PFA (w/v), followed by three washes with 1xPBS. The samples were cryoprotected in 30% sucrose (w/v) solution for a minimum of 24 hours or until the tissue had sunk in the solution. Subsequently, the tissue was rinsed with 1X PBS, allowed to reach room temperature, and transferred to plastic cryomolds covered with OCT. The tissue was snap-frozen with liquid nitrogen in an isopentane bath and stored at -80°C prior to sectioning. Sectioning was performed using a Leica CM1950 cryotome to obtain 10 µm thick sections, and placed on to Suprfrost adhesion microscope slides, and stored at -80°C prior to immunohistochemistry. The samples were allowed to defrost for 15-20 minutes at room temperature, then hydrated with three 1X PBS washes. The sections were permeabilized with 0.25% Triton-X100 in 1X PBS for 5 minutes at room temperature and washed three times with 1X PBS. Tissue sections were then blocked for 1 hour with 0.1% Saponin and 10% horse serum in 1X PBS. After the blocking step, the sections were rinsed three times with 1X PBS and incubated overnight at 4°C with primary antibodies diluted in 0.1% Saponin and 10% horse serum in 1X PBS. The next day, the sections were thoroughly washed three times with 1X PBS and incubated for 1 hour at room temperature in the dark with secondary antibodies and DAPI, diluted in 0.1% Saponin and 10% horse serum in 1X PBS. Following the 1-hour incubation, the sections were washed three times with PBS. Tissue sections were incubated with 0.05% 20X TrueBlack Lipofuscin Autofluorescence Quencher (Biotium, 23007) in 70% ethanol for one minute and washed thoroughly three times with 1X PBS. The tissue sections were mounted using Fluoromount Aqueous Mounting Medium (Sigma Aldrich, F4680), then allowed to dry overnight in the dark at 4°C.

### Confocal microscopy

Confocal microscopy was performed on a LSM700 confocal microscope (Zeiss) with a 40 X Plan Apochromat oil objective (numerical aperture, NA = 1.3) or a 63 X Plan Apochromat oil objective (NA = 1.4) equipped with 405 nm (5mW), 488 nm (10 mW), 555 nm (10 mW) and 639 nm (5 mW) lasers. Emission light path was adjusted on the Zeiss ZEN imaging software for 488 nm and 555 nm excitation to eliminate autofluorescence in yellow wavelengths. Images were acquired at 1,024x1,024 pixel size with zoom factor set to 1.0-1.5. To compare signals in proliferating and quiescent cells, laser intensity and gain were kept constant when imaging the two conditions for a protein of interest.

### High-throughput automated widefield microscopy

Automated high-throughput image acquisition was performed on an ImageXpress Micro XL Widefield High-Content Screening System (Molecular Devices) equipped with a Nikon ELWD 20X S Plan Fluor air objective (NA = 0.45) using MetaXpress 5.3 software. Fluorophores were excited with a Lumencor Sola solid-state white-light engine which wavelengths were narrowed down through the following excitation filters and emission was detected with the following band-pass filter cubes: Hoechst33342 or DAPI were imaged using 377/50 and 477/60 nm Excitation and Emission filters, AlexaFluor488 or DiO using 482/35 and 536/40 nm Excitation and Emission filters, AlexaFluor555, Dil or Rhodamine using 543/22 and 593/40 nm Excitation and Emission filters, AlexaFluor594 using 565/40 and 624/40 nm Excitation and Emission filters, and AlexaFluor633 or AlexaFluor647 using 628/40 and 692/40 nm Excitation and Emission filters. Emitted light was captured by a 4.66 megapixel complementary metal-oxide semiconductor (CMOS) camera. Images were acquired as 16-bit grayscale images at 2,160x2,160 pixels with one image per acquired channel.

The workflow for setting up automated imaging of a 96 well-plate was as followed: (1), calibration of the plates to allow for plate and well bottom autofocus setup (using a dedicated 690 nm 20 mW laser); (2) selection of the ELWD 20X NA=0.45 S Plan objective; (3) selection of the number and location of wells (up to 96) and intra-well sites and fields of view (6-18 per well) to be imaged selection; (4) Acquisition loop: i) Laser-based focusing on plate and well bottom was enabled for each image to be acquired, ii) selection of the channels of wavelengths for excitation and emission (up to 4), iii) selection of the channel serving as focus offset for each other channel and iv) determined for each channel of image-based focus and exposure times, ranging from 1-2,000ms; and (5) Application of focus journal after each image was acquired To account for unevenness in plate design or differences in focus planes. The journal encoded a z-stack (for example 4 images with 2 μm spacing between each image, set manually) from which the image with the best focus was saved as final acquisition.

The workflow for setting up automated imaging of histological slides was similar (1) Using the Slide Scanning journal within the Molecular Devices ImageXpress control software, calibration of the slides to allow for correctly autofocused images was setup (using a dedicated 690 nm 20 mW laser); (2) A slide pre-scan at 4x magnification in the DAPI channel was undertaken before rectangular regions of interest of a 21x21 non overlapping fields of view array created to contain each of the three sections on the slide. selection of the ELWD 20X NA=0.45 S Plan objective followed by (3) manual selection of the of the number and location of wells (up to 96) and intra-well sites and fields of view (6-18 per well) to be imaged selection; (4) Acquisition loop: i) Laser-based focusing on plate and well bottom was enabled for each image to be acquired, ii) selection of the channels of wavelengths for excitation and emission (up to 4), iii) selection of the channel serving as focus offset for each other channel and iv) determined for each channel of image-based focus and exposure times, ranging from 1-2,000ms; and (5) Application of focus journal after each image was acquired To account for unevenness in plate design or differences in focus planes.

The journal encoded a z-stack (for example 4 images with 2 μm spacing between each image, set manually) from which the image with the best focus was saved as final acquisition.

### *In vitro* automated image analysis

Image analysis was conducted using a protocol established in CellProfiler image analysis software. A set of image analysis algorithms were constructed to measure the properties of interest within the RPE1 cell culture labeled with DAPI, and other targets of interest, as required. Each image set, corresponding to one field of view or site and comprising three or four fluorescent channels as appropriate, were analyzed independently using this pipeline. Between 4 and 12 fields of view per well were analysed with experiments repeated in triplicate unless indicated otherwise. The analysis cell segmentation pipeline comprises an illumination correction function calculated for each channel using a median filter (300 x 300 pixels) to correct for illumination variations across each 96-well plate. Quality control filtration of regions of clustered fluorescent signal, associated with concentrated DAPI fluorescence with diameter between 155 and 1500 pixel units were then masked from each image set. Cells nuclei of each cell in the field of view were segmented based on arbitrary fluorescence intensity associated to DAPI DNA labelling, a median size (20 to 60 pixels), and shape (circular). Next, cells bodies were segmented by propagating from the nuclei using an arbitrary fluorescence intensity corresponding to the red or green, fluorescent channel, depending on appropriate cytoplasmic signal and an Otsu three class thresholding method. Finally, cytoplasmic regions of cells were segmented by masking the nuclear associated region from the cell body. Fluorescence intensity, shape and fluorescent ratio between cytoplasm, cell body and nuclear regions per cell were then measured. Interexperimental normalization per labelling was undertaken by dividing each data point by the associated median fluorescent signal of the labelling in the experiment. Data per condition were determined by first aggregating cell signals or the median signal per cell per field of view, as indicated, and then over replicate wells and experiments. Fractional results of the three subcellular compartments measured are expressed as a percentage of their values normalized to a total a control sample and the minimum number of RPE1 cells per condition per experiment quantified was greater than 1,000 and upwards to 25,000 cells per condition as appropriate. A generalized Cell Profiler pipeline associated to this analysis in this study is available on the cell profiler website and in the supporting martial accompanying this study.

### *In vivo* automated image analysis

Samples from three different pigs were analysed, with three sequential sections from one pig being placed on an imaging slide. Two were treated as technical repeats and positively labelled as above with Dapi, pERK, (an ECM of interest) and LGR5-Alexa Fluor 647, and whereas the third section was treated as a background control and labelled with only DAPI. Output images per section were processed through the entire image analysis pipeline based on the following approach. Firstly, to establish a baseline for the background signal, the overall fluorescent intensity per sample was measured. Next, the interstitial tissue regions of interest were segmented based on elevated DAPI fluorescent intensity, closeness and the minimum and maximum size of individual objects in pixels (min-400 to max-1,000,000) using an adaptive two class Otsu thresholding strategy, with holes in the segmented objects being filled in. Anything falling within this size range or that was dimmer than the median object florescence was excluded, and a series of masks were generated in each of the four channel images to isolate the interstitial tissue for further analysis. To eliminate artifacts such as bubbles arising from mounting media, dust, and debris, another mask based on size and fluorescent intensity was applied across all four channels, as these artifacts tend to be larger than cells and exhibit higher fluorescence intensity across all channels. Once the preprocessing steps mentioned above were completed, the images were fully processed through the existing pipeline. LGR5 positive cells were identified in a size range of 4 to 40 pixels and that that were above a minimum fluorescent intensity level determined by the overall background fluorescent signal associated with the background image. ECM positive concentrations and pERK positive cells were independently identified and segmented in their respective channels using a similar approach with identical size ranges and modulated minimum fluorescence as appropriate. Next, identified segmented object intensities and sizes were measured in all channels, before being exported to a database for further processing in R-studio.

Initially, data frames associated LGR5, ECM, DAPI and pERK identified objects from all experiments were generated in R-studio 4.3.3 for quality control, normalisation, and analysis. Quality control and artefact detection was undertaken by calculating the median objects segmented per FOV, identifying those FOVs that showed segmented objects that deviated significantly from this value. In practical terms this equated to FOVs with more than 50 objects detected. These FOV images and their segmentation overlays were then manually inspected and discarded or retained based on labelling artefact detection across four channels assessed. Secondly, the numbers of objects detected and segmented within the positively labelled sections were compared to those of the DAPI labelled control section to ensure fluorescent thresholding for segmentation were accurate, yielding minimal objects identified in the blank control. Thresholding values were altered iteratively within a slide set analysis to ensure labelled object detection and segmentation with high accuracy.

Next, downstream normalisation in R-studio was undertaken using the following transformations:

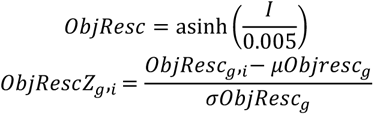

- Obj_resc: The result of the asinh transformation applied to the intensity data across each of the four channels measured for each object class segmented (ECM or LGR5 or pERK)
- *I*: The intensity measurements of Segmented Objects (ECM or LGR5 or pERK).
- asinh: The inverse hyperbolic sine function.
- *g*: The group defined by the labelling.
- *i*: The index of the data point within group *g*.
- ObjResc_g,i_: The the asinh rescaled intensity value the *i*-th data point within group *g*.
- μObjResc*g*: The mean of ObjResc within group *g*.
- σObjResc*g*: The standard deviation of ObjResc within group *g*.
- ObjRescZ: The result of the Z-score data transformation of Obj_resc

To evaluate data Lgr5 cells associated with laminin ECM concentration, Lgr5 cell objects were subset against a minimum ECM signal corresponding to a Z-score greater than 0 for all ECM targets examined. Violin plots represent the Z-scored fluorescent intensity signal of pERK labelling from the aggregated subset Lgr5 positive objects from at least 6 specimen sections equally spread across three pigs imaged in 21x21 FOV grids per section. A generalized Cell Profiler pipeline and R script associated to this analysis from this study is available on the cell profiler website and in the supporting martial accompanying this study as appropriate.

### Statistics and reproducibility

Gaussian distribution of samples was tested with D’Agostino & Pearson omnibus normality test. Samples following a Gaussian distribution were tested for statistical significance with Student’s unpaired two-tailed t-test (two sample groups), one-way ANOVA and Dunett’s test for multiple comparisons (more than two sample groups) or two-way ANOVA with Tukey’s test for multiple comparisons (more than two sample groups with two parameters per group). If samples did not follow a Gaussian distribution, an unpaired t-test with Welsh’s correction was used to compare two sample groups and Kruskal-Wallis test with Dunn’s multiple comparisons test for more than two sample groups. Significance of mean comparison coefficients was annotated as follows: **** P < 0.0001, *** P < 0.001, ** P < 0.01, * P < 0.05, *ns*, not significant.

## DATA AVAILABILITY

The source data for statistical analysis of all the measurements used in this study will be provided after acceptance and before publication in the Source Data File. Complete proteomics and phosphoproteomics (LC-MS/MS) data is included as Supplementary Table 1 and 2 and will be deposited in ProteomeXchange Consortium via the PRIDE partner repository after acceptance and before publication. All uncropped gels and immunoblots are provided in Supplementary Figures S16 to 18. All other data that support the findings of this study will be made available from the corresponding authors on reasonable request after publication.

## SUPPLEMENTARY FIGURE LEGENDS

**Supplementary Figure S1.**
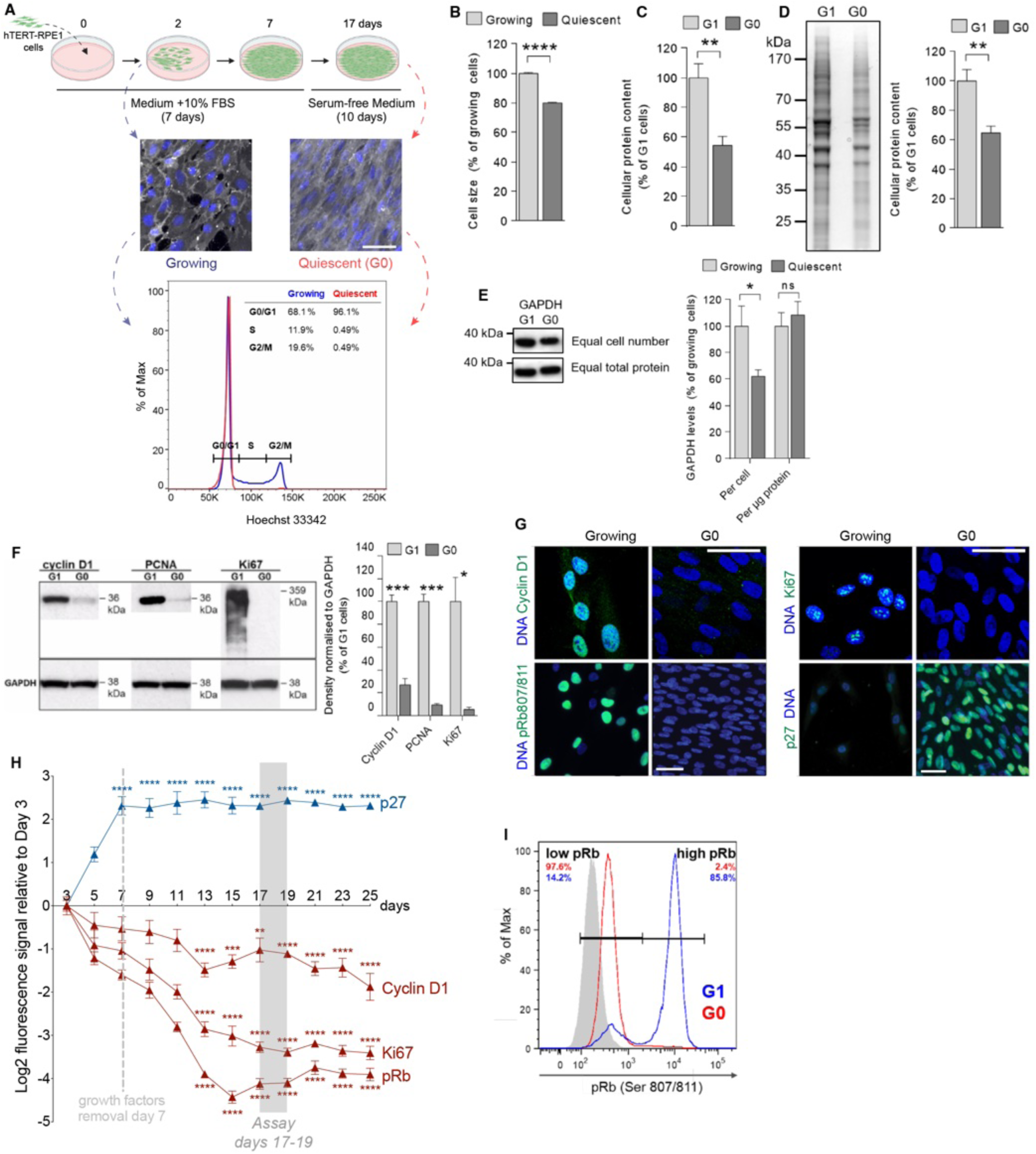
Establishment and characterization of an *in vitro* model of cellular quiescence. (A) Top, schematic of in vitro cell quiescence protocol: exponentially growing human hTERT-RPE1 cells were plated at low density and grown for 7 days in full serum medium (refreshed every 2 days), at which point they formed homogeneous monolayers. From Day 7, cells were maintained in serum-free medium (refreshed every 2 days), and used on Day 17 for experiments. Middle, representative images of Growing cells (Day2) or Quiescent (Day 17) cells, immunostaining for a plasma membrane marker (anti-CD44, white) and DNA (blue). Bottom, Representative histogram showing the DNA profile of ethanol-fixed growing (blue) and quiescent (red) RPE1 cells stained for DNA with Hoechst33342 and analyzed by flow cytometry. Gates represent the cut-off lines for cell cycle phases G0/G1, S and G2/M, which are quantified in the upper right corner. Percentage of max (% of Max) indicates the number of cells relative to the peak fraction of cells. A minimum of 50,000 cells per condition were analyzed. (B) Mean diameter of growing and quiescent cells in suspension measured with a haemocytometer. N=15 (growing cells) and N=11 (quiescent cells) biological replicates. (C) Protein concentration measurement using Bradford assay. G0 and FACS-sorted G1 cells were lysed in 8M urea, with volumes adjusted to cell numbers. N=4 biological replicates. (D) Left, representative lanes of Coomassie blue-stained SDS- PAGE gel with lysates from G0 and FACS-sorted G1 cells lysed in equal volumes of 8M urea per cell number. Right, color intensity per lane was quantified by densitometry measurement and normalized to G1 cells. N=5 biological replicates. (E) Left, representative immunoblots of total lysates from FACS-sorted G1 and G0 RPE1 cells probed against GAPDH. The upper blot shows loading normalized to cell number and the lower blot shows loading normalized to total protein levels. Right, GAPDH levels per band quantified by densitometry measurement. N=5 (per cell) or N=6 (per μg protein) biological replicates. (F) Left, representative immunoblots of total lysates from G0 and FACS-sorted G1 cells probed against Cyclin D1, PCNA and Ki67. GAPDH served as loading control. Right, Densitometry quantification of 3 independent immunoblots as shown in (Left). (G) Representative fluorescence microcopy images of Growing or G0 cells immunostaining for endogenous (all green) Cyclin D1, Ki67, phosphorylated Ser807/811 Retinoblastoma protein (pRb) and p27^kip1^ (p27) and DNA (blue). (H) Mean fluorescence (Log2) form 3 independent experiments of the indicated endogenous immunostainings Cyclin D1, Ki67, pRb and p27 of quiescent cell cultures measured every 2 days over 25 days. (I) Histogram of pRb in G1 and G0 cells analyzed by flow cytometry. After elimination of cell doublets and dead cells, the population with 2n DNA content was used for further analysis. Gates used for pRb quantification are displayed. Percentages of G1 (blue) and G0 (red) cell populations with low and high pRb are listed in the left and right upper corner, respectively. Blank signal is shown in grey. A minimum of 50,000 cells per condition were analyzed. In A and G, the scale bars represent 50 μm. In B, C, D, and F, bars show the mean and error bars represent SEM. *P* values calculated by Student’s unpaired two-tailed t-tests. *\*P<0.05, **P<0.001, ***P<0.001, ****P<0.0001*

**Supplementary Figure S2.**
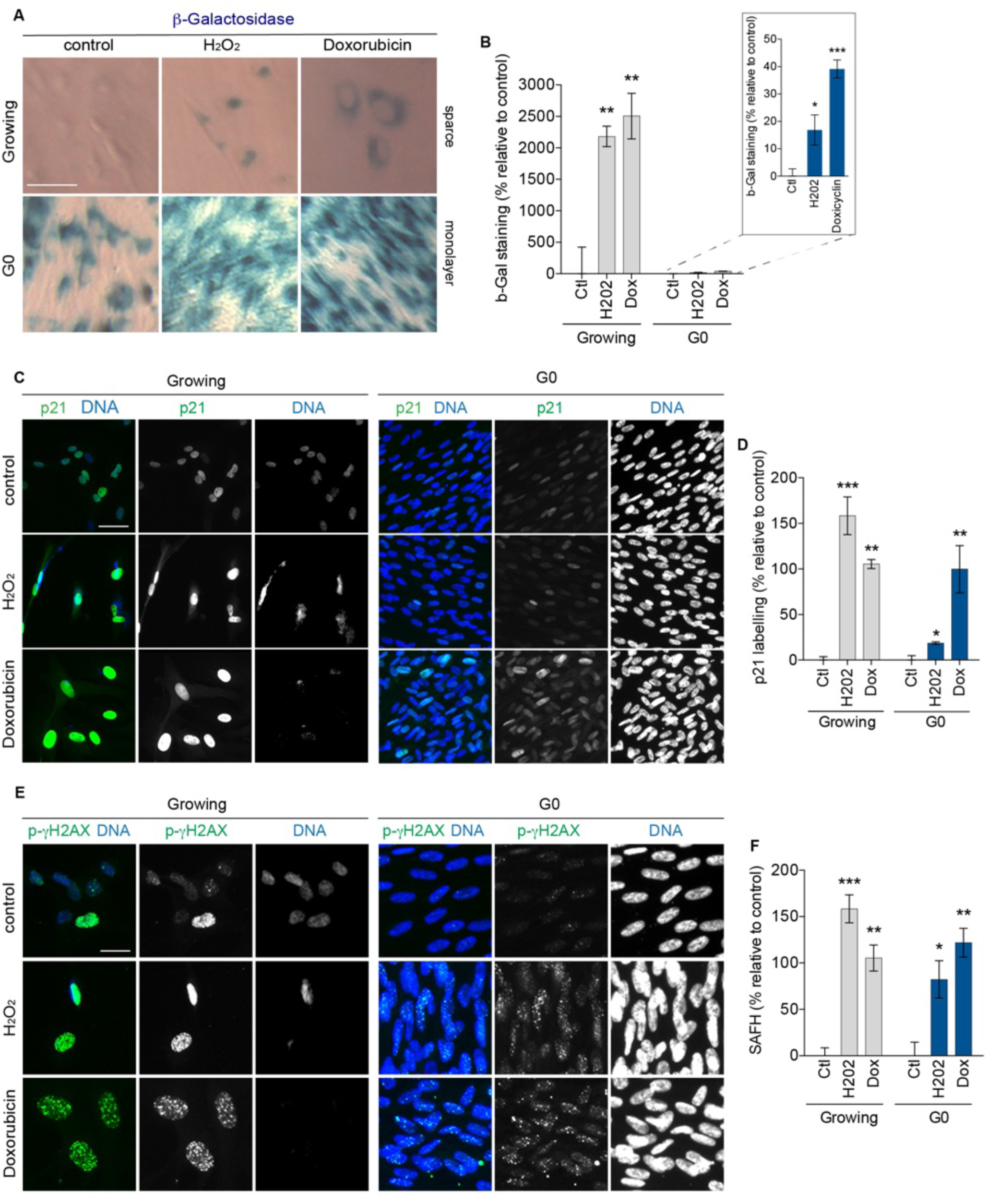
Cells from *in vitro* model of cellular quiescence are not senescent. (A) Representative brightfield microscopy images of Growing and quiescent (G0) cells, treated with 500 μM H_2_O_2_ for 2 hours or 50 nM Doxorubicin for 4 days and stained with β-Galactosidase Staining kit (blue). (B) Mean intensity β-Galactosidase staining (normalized to control) in Growing or G0 cells treated as in (A). (C) Representative fluorescent microscopy images of Growing and quiescent (G0) cells, treated with 500 μM H_2_O_2_ for 2 hours or 50 nM Doxorubicin for 4 days and immunostained for endogenous p21^WAF1/CIP1^ (‘p21’, green) and DNA (blue). (D) Mean fluorescence of p21 levels, normalized to control, in cells treated as in (C). (E) Representative fluorescent microscopy images of Growing and quiescent (G0) cells, treated with 500 μM H_2_O_2_ for 2 hours or 50 nM Doxorubicin for 4 days and immunostained for endogenous nuclear phosphoryated Ser139 Histone H2A.X (‘p-γH2AX’, green) and DNA (blue). (F) Mean fluorescence of p-γH2AX levels, normalized to control, in cells treated as in (E). In A, C and E, the scale bars represent 50 μm.

**Supplementary Figure S3.**
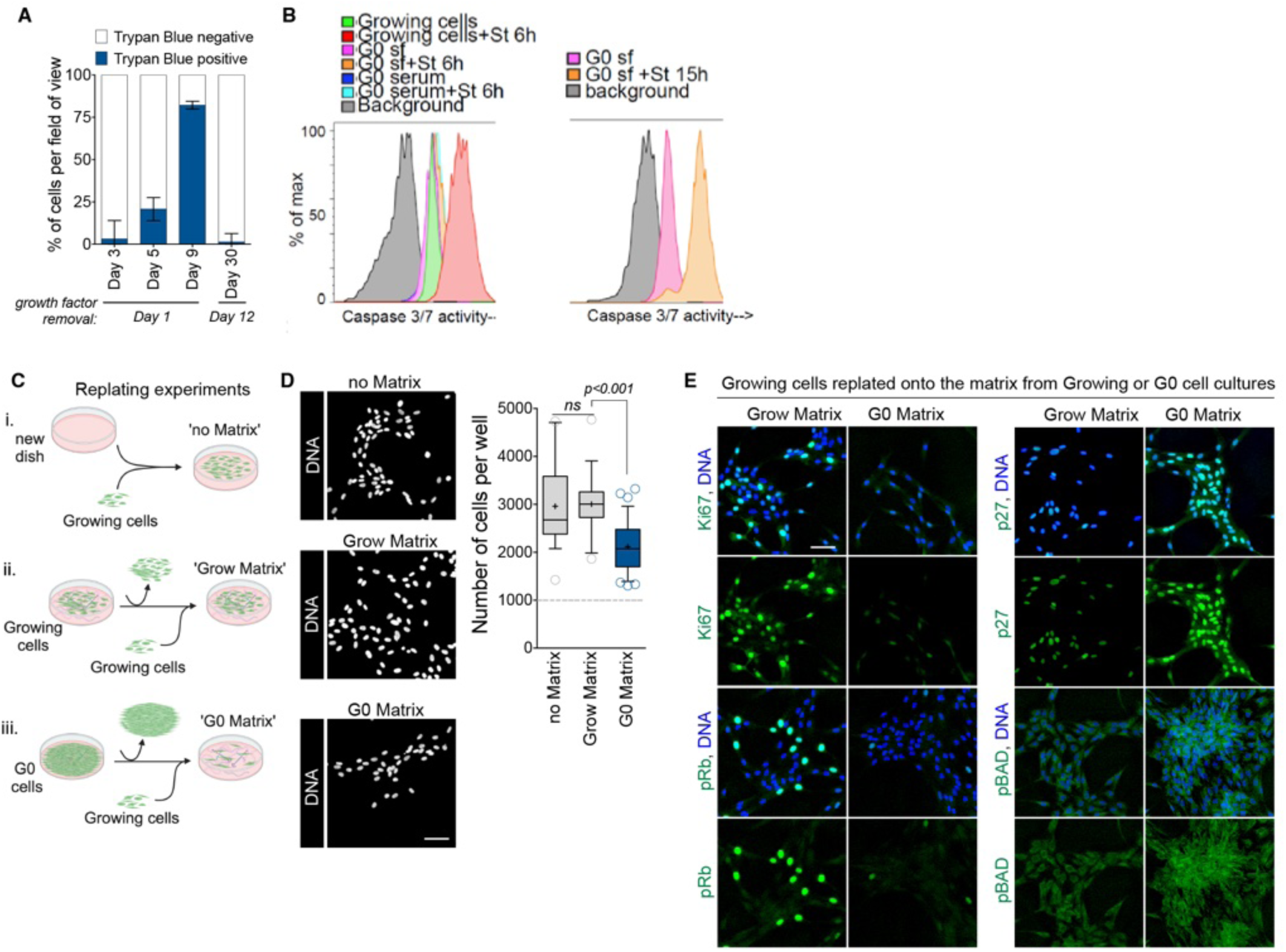
Cell cycle exit and long-term survival of quiescent cells originate from their ECM. (A) Percentage of cells per field of view, grown for the indicated times and either switched to serum-free medium adfter Day 1 or after Day 12 and positive (blue) or negative (white) for Trypan Blue. (B) Left, histogram of Caspase 3/7 activity analyzed by flow cytometry in Growing cells challenged with 2 μM staurosporine for 6h (red) or not (green), or G0 cells maintained either in serum-free medium (sf) and challenged with staurosporine (orange) or not (pink) or in full-serum medium (serum) and challenged with staurosporine (cyan) or not (blue). Blank signal (background) is shown in grey. A minimum of 50,000 cells per condition were analyzed. Right, histogram of Caspase 3/7 activity analyzed by flow cytometry in G0 cells maintained in serum-free medium (sf) and challenged with 2 μM staurosporine for 15h (orange) or not (pink). Blank signal (background) is shown in grey. A minimum of 50,000 cells per condition were analyzed. (C) Schematic of replating experiments: i) ‘no Matrix’ was growing cells plated onto new tissue culture dishes, ii) ‘Grow Matrix’ was the ECM naturally secreted by growing cells previously cultured for 4 days, and removed by accutase, before the replating of cells from unrelated, exponentially growing, samples, iii) ‘G0 Matrix’ was the ECM naturally secreted by quiescent cells previously cultured for 17 days, and removed by accutase, before the replating of cells from unrelated, exponentially growing, samples. (D) Left, representative fluorescence microscopy images of the DNA of Growing cells replated onto dishes previously coated with ‘Grow Matrix’, or ‘G0 Matrix’ or not (‘no Matrix’), as defined in (d) and grown for 6 days in serum-free medium. Right, number of cells per well, as indicated. The dotted line shows the number of cells initially seeded (1x10^3^ cells per well). (E) Representative fluorescence microscopy images of Growing cells replated onto dishes naturally coated with ‘Grow Matrix’ or ‘G0 Matrix’, as defined in (D) and immunostaining for endogenous Ki67, p27^kip1^ (p27) phosphorylated Ser807/811 Retinoblastoma protein (pRb), or pBAD (all green), and DNA (blue). In D and E, the scale bars represent 50 μm. In D, crosses show the mean, horizontal lines show the median and whiskers show the 10 to 90^th^ percentiles of values. Outliers are shown as circles. *P* values calculated by one-way ANOVA (D).

**Supplementary Figure S4.**
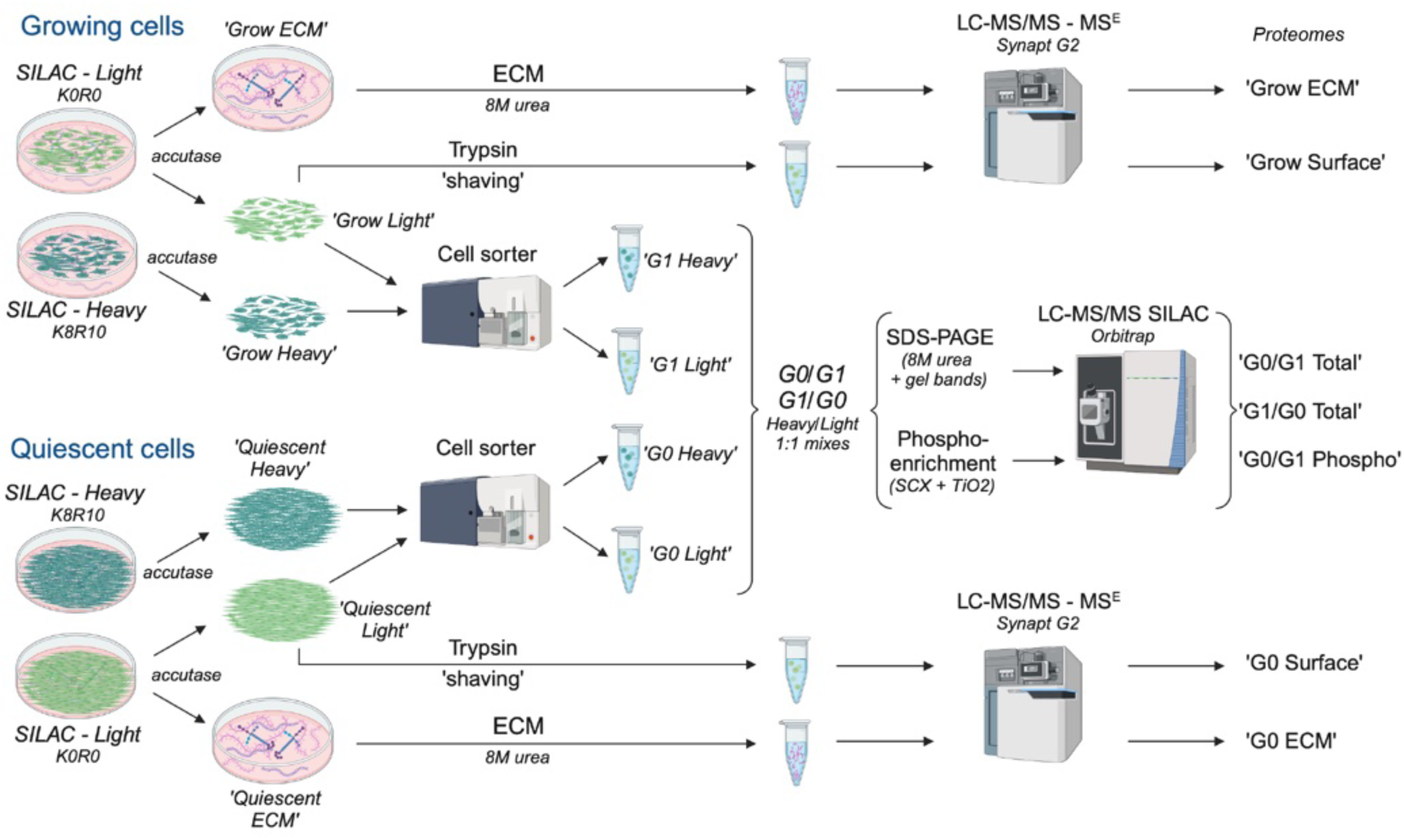
Cell surface, ECM, Total protein and Phosphorylation Mass Spectrometry pipelines. Schematic of Surface, ECM and Total proteomes and phosphoproteomes performed in this study. For Surface and ECM proteomes, growing or quiescent cells were detached using accutase, then their cell surface proteins were ‘shaved’ using Trypsin and collected (‘Grow Surface’ and ‘G0 Surface’). ECM and proteins left on tissue culture dishes were collected and lysed in 8M urea (‘Grow ECM’ and ‘G0 ECM’). Proteins were identified and quantified using label-free quantitative mass spectrometry (LC-HDMS^E^). For Total proteomes and phosphoproteomes, growing or quiescent cells grown in Light (K0R0) or Heavy (K8R10) SILAC medium (*see STAR Methods*) were detached using accutase, FACS-sorted to isolate 2n DNA population (‘G1 Heavy’, ‘G1 Light’, ‘G0 Heavy’ and ‘G0 Light’ fractions), lysed in 8M urea and mixed 1:1 before gel band fractionation or Phospho-enrichment (*see STAR Methods*). Proteins and phosphorylations were identified and quantified using an Orbitrap mass spectrometer.

**Supplementary Figure S5.**
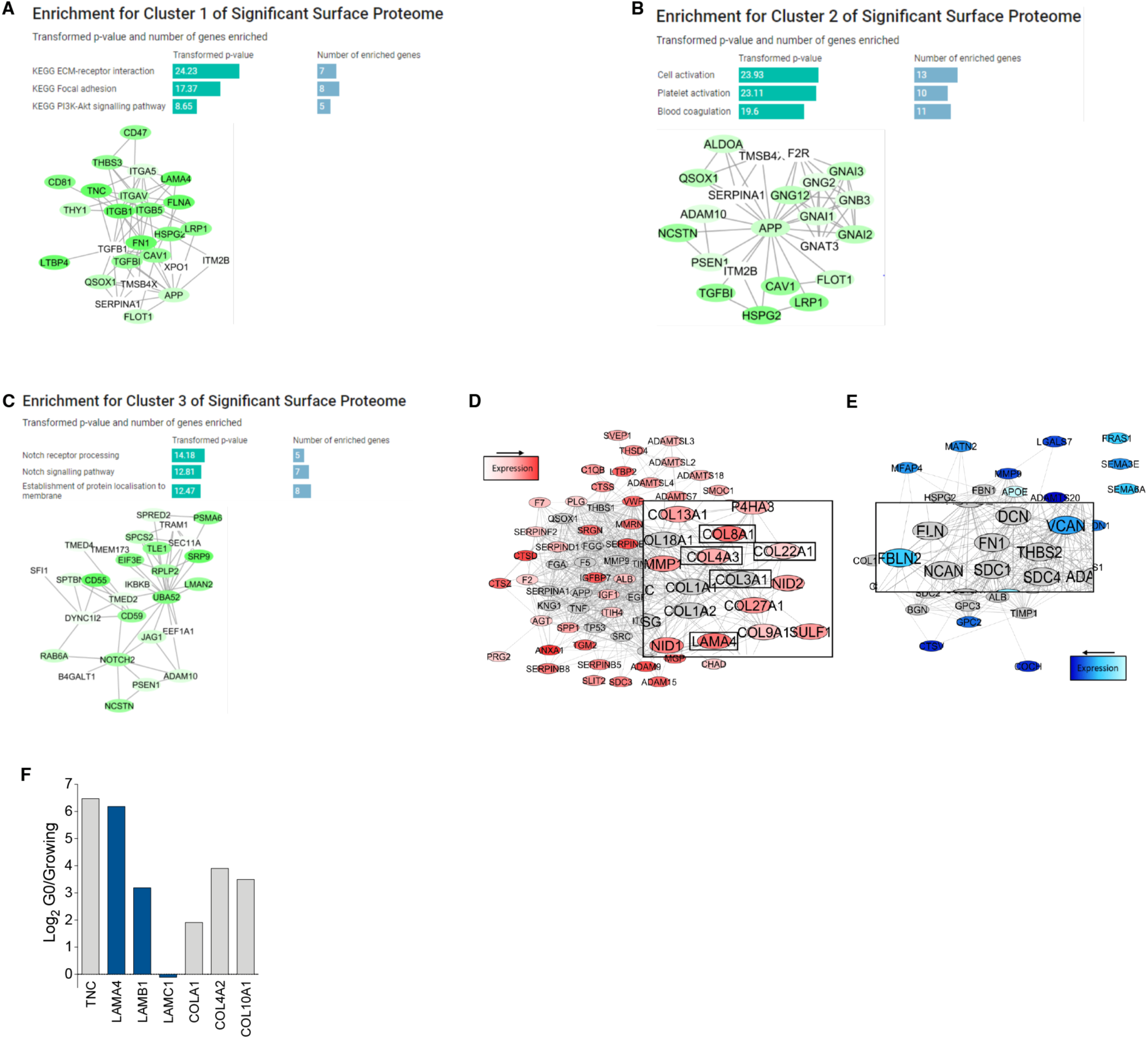
Cell Surface Proteome analysis - upregulated proteins during G0. (A-C) Clusters of proteins significantly enriched from the surface proteome from (Fig. 1G) containing 35, 22 and 28 proteins, respectively. The proteins are color-coded based on the expression data with the darker green colors having a higher expression in G0 compared to their expression in G1. The enrichment results from these clusters are represented in the bar charts with the *p-value* of each KEGG term transformed and the number of proteins within the clusters involved in each enrichment stated. (D) ECM proteins upregulated during quiescence. (E) ECM proteins downregulated during quiescence. (F) Fold changes (Log2) in selected protein between G0 and Growing Surface and ECM fractions detected as in (Fig. 1G).

**Supplementary Figure S6.**
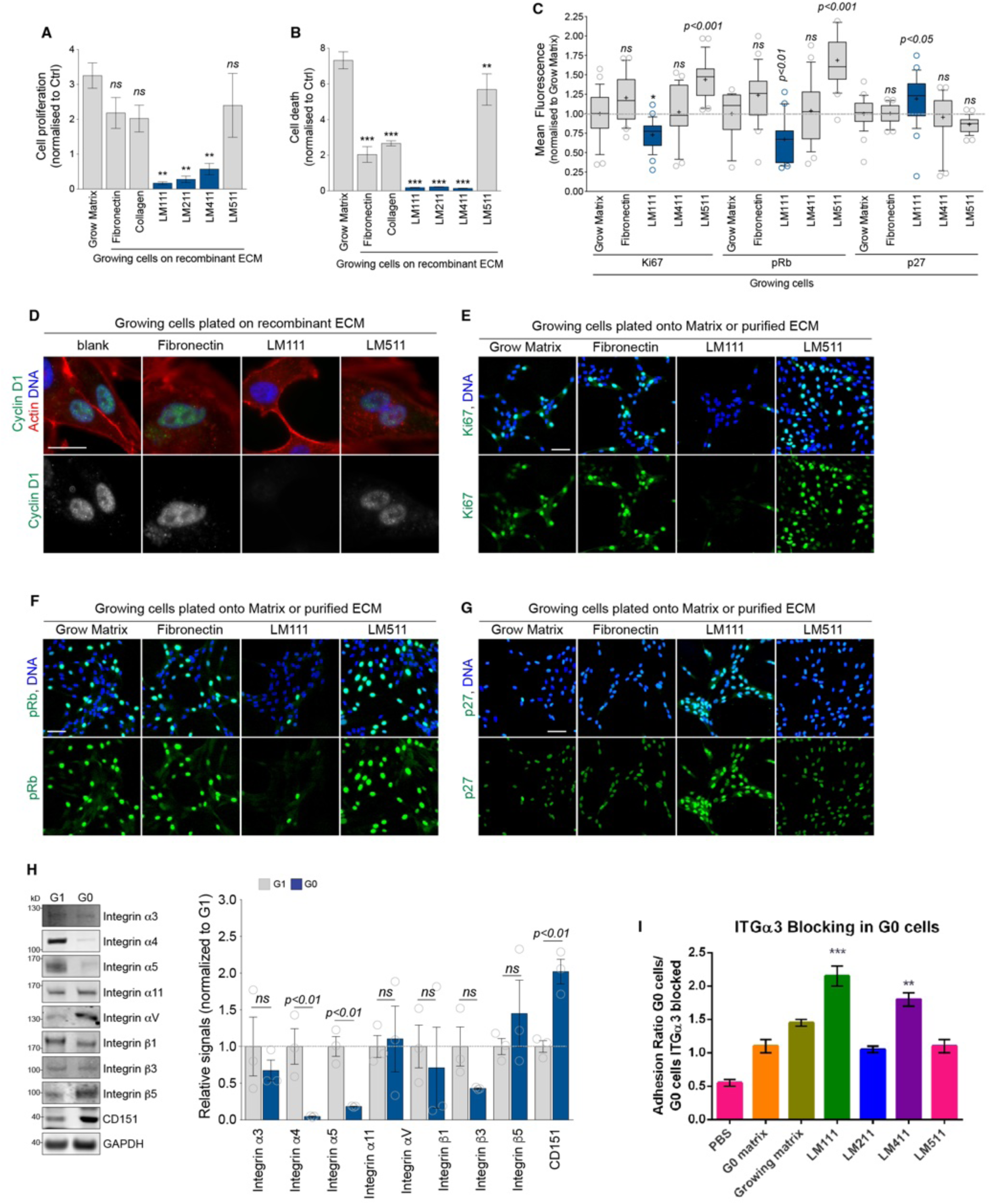
Laminin 111 is sufficient to trigger cell cycle exit of growing cells. (A) Cell proliferation index (normalized to no coating control) of growing cells replated onto Grow Matrix or the indicated recombinant ECM and measured using a xCELLigence RTCA DP E-plate system. (B) Cell death index (normalized to no coating control) of sub confluent growing cells seeded in full serum for 12 hr before being cultured for a further 10 days in SF media and analyzed as in (A). (C) Mean fluorescence of endogenous Ki67, pRb or p27 in growing cells replated onto Grow Matrix, purified Fibronectin or recombinant LM111, LM411 or LM511 and grown for 3 days in full-serum medium. Part of this panel was used in Figure 1k. (D) Representative fluorescence microscopy images of growing cells replated onto Grow Matrix, purified Fibronectin or recombinant LM111 or LM511 and grown for 6 days in serum-free medium, and immunostained for endogenous Cyclin D1 (green), actin (red) and DNA (blue). (E) Representative fluorescence microscopy images of growing cells replated onto Grow Matrix, purified Fibronectin or recombinant LM111 or LM511 and grown for 6 days in serum-free medium, and immunostained for endogenous Ki67 (green) and DNA (blue). (F) Representative fluorescence microscopy images of growing cells replated onto Grow Matrix, purified Fibronectin or recombinant LM111 or LM511 and grown for 6 days in serum-free medium, and immunostained for endogenous phosphorylated Ser807/811 Retinoblastoma protein (pRb, green) and DNA (blue). (G) Representative fluorescence microscopy images of growing cells replated onto Grow Matrix, purified Fibronectin or recombinant LM111 or LM511 and grown for 6 days in serum-free medium, and immunostained for endogenous p27^kip1^ (p27, green) and DNA (blue). (H) Left, representative immunoblots showing the levels in Integrin Integrin α3, α4, α5, α11, αV, β1, β3, β5 or CD151 in G1 or G0 cell extracts (isolated by cell sorting, *see STAR Methods*). Uncropped gels are displayed in Supplementary Fig. S16-18. Right, relative signals normalized to GAPDH, of 3 independent experiments as in (Left). (G) ITGα3 blocking in G0 cells and quantifying their adhesion to various laminin subtypes, PBS, G0 and growing cell naturally deposited ECM. Results represent mean ± SEM (biological N=3). One-way ANOVA, Bonferroni test ****p≤ 0.0001, **p≤ 0.01, Scale bar 20 (C) and 50 (D-F) μm.

**Supplementary Figure S7.**
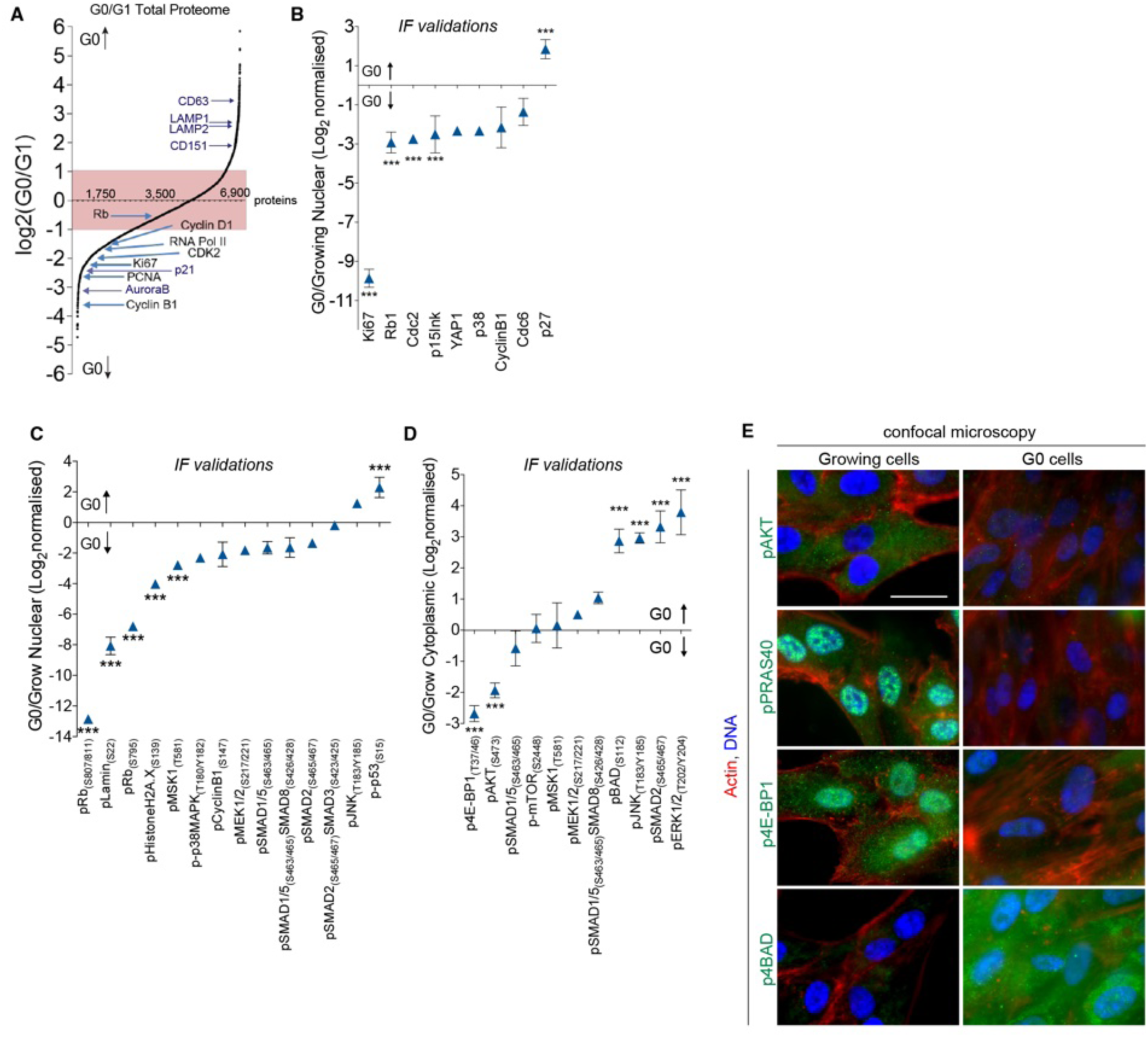
Total proteome ad phosphoproteome validations. (A) S’ plot showing the relative abundance in 6,982 proteins measured by SILAC mass spectrometry, and ranked by Log2 normalized ratio (G0/G1). The pink shaded area highlights the range between -1 to +1 ratio, beyond which variations were considered significant (*see STAR Methods*). (B) Log2 normalized ratio (G0/Growing) of the mean fluorescence in the nucleus of G0 or Growing cells immunostaining for endogenous Ki67, Rb, Cdc2, p15^ink^, YAP, p38, Cyclin B1, Cdc6 and p27. (C) Log2 normalized ratio (G0/Growing) of the mean fluorescence of in the nucleus G0 or Growing cells immunostaining for endogenous phosphitylations, as indicated. (D) Log2 normalized ratio (G0/Growing) of the mean fluorescence of in the nucleus G0 or Growing cells immunostaining for endogenous phosphitylations, as indicated.

**Supplementary Figure S8.**
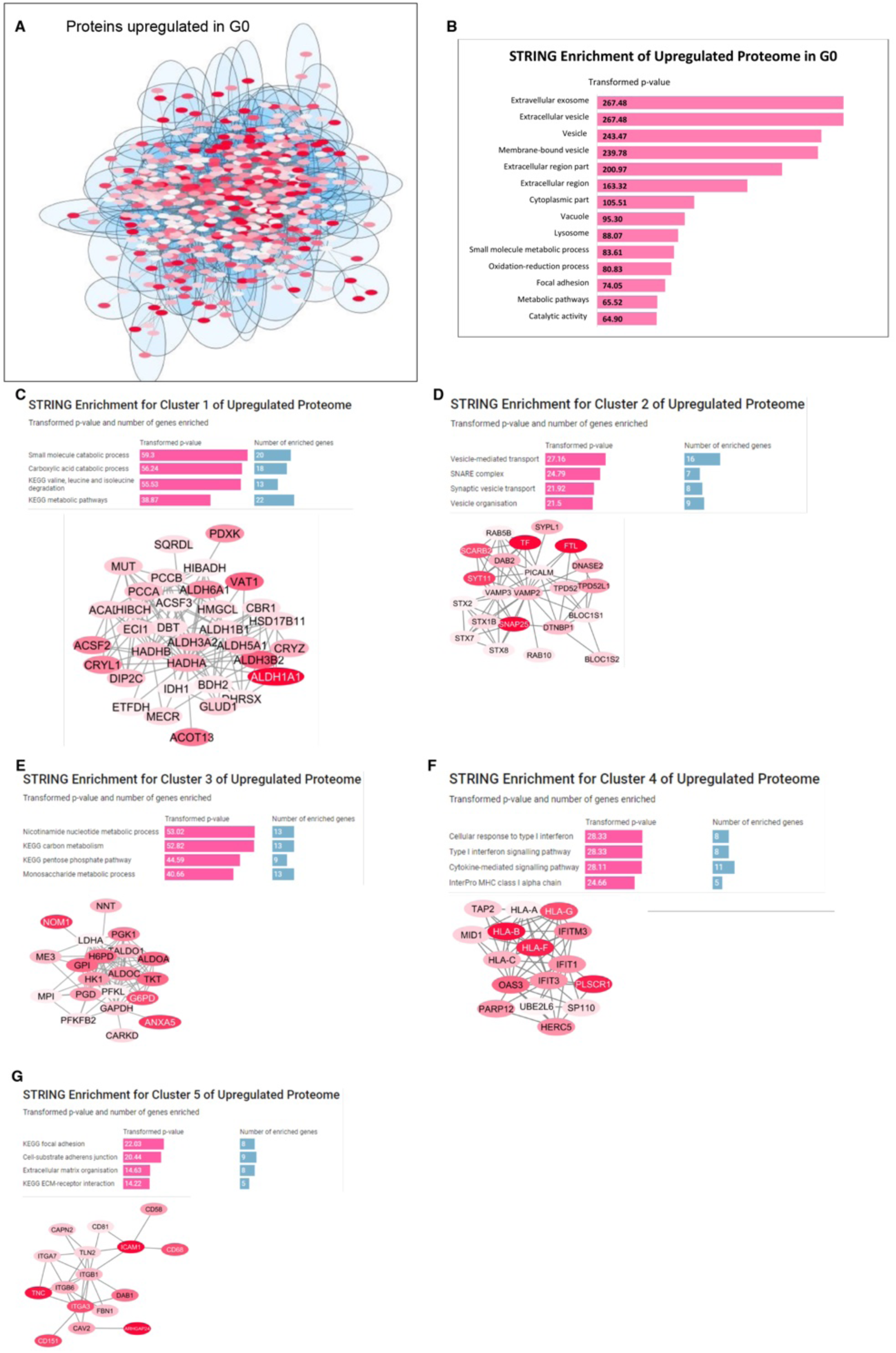
Total Proteome analysis - upregulated proteins during G0. (A) Overall interaction network generated with Cytoscape using the STRING database between 790 proteins significantly upregulated from the Total Proteome. Blue circles show 119 clusters detected by AutoAnnotate, which clusters proteins with a shared STRING term. Nodes are color-coded based on expression values with the deeper red colors indicating higher expression in G0. (B) STRING enrichment of the 790 proteins significantly upregulated from the Total Proteome the predicted the following GO and KEGG pathways to be the most enriched. The *p*-value of each enrichment was natural log transformed. (C-G) The five largest clusters from (A) containing 34, 22, 20, 16 and 16 proteins, respectively. The proteins are color-coded based on the expression data with the darker red colors having a higher expression in G0 compared to their expression in G1. The enrichment results from these clusters are represented in the bar charts with the *p-value* of each KEGG term transformed and the number of proteins within the clusters involved in each enrichment stated.

**Supplementary Figure S9.**
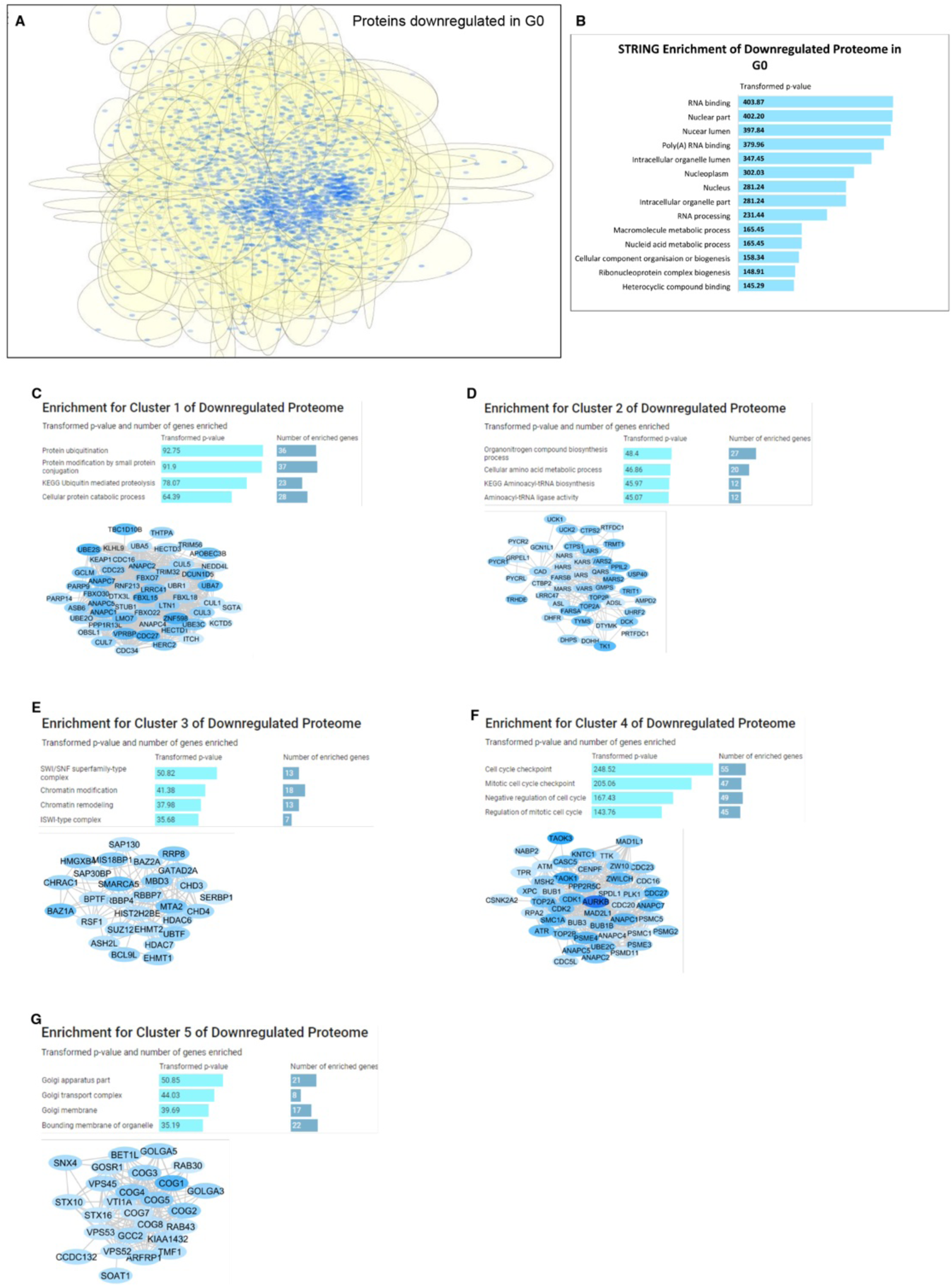
Total Proteome analysis - dowregulated proteins during G0. (A) Overall interaction network generated with Cytoscape using the STRING database between 2,000 proteins significantly downregulated from the Total Proteome. Yellow circles show 239 clusters detected by AutoAnnotate, which clusters proteins with a shared STRING term. Nodes are color-coded based on expression values with the deeper blue colors indicating higher expression in G0. (B) STRING enrichment of the 2,000 proteins significantly downregulated from the Total Proteome the predicted the following GO and KEGG pathways to be the most enriched. The *p*-value of each enrichment was natural log transformed. (C-G) Five clusters from (A) containing 53, 45, 28, 55 and 26 proteins, respectively. The proteins are color-coded based on the expression data with the darker blue colors having a lower expression in G0 compared to their expression in G1. The enrichment results from these clusters are represented in the bar charts with the *p-value* of each KEGG term transformed and the number of proteins within the clusters involved in each enrichment stated.

**Supplementary Figure S10.**
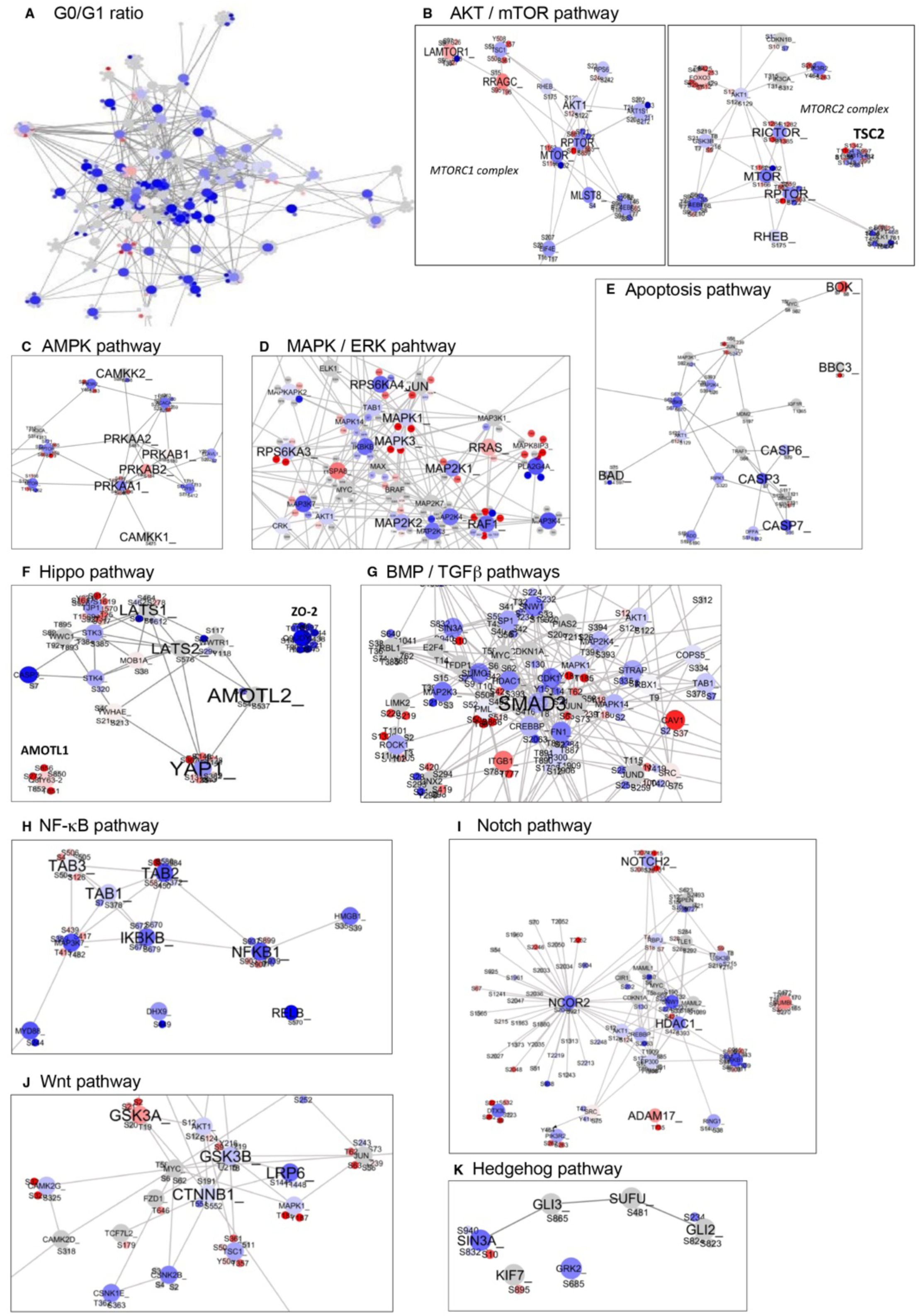
Phosphoproteome analysis - changes in major signaling pathways during G0. (A-K) PhosphoPath snapshots of the broad G0:G1 phosphoproteme (A), or focused on the following signaling pathways: AKT/mTOR (B), AMPK (C), MAPK/ERK (D), Apoptosis (E), Hippo (F), BMP/TGFβ (G), NFκB (H), Notch (I), Wnt (J) and Hedgehog (K) pathways. Blue indicates proteins and phosphorylation that were reduced during G0 (negative Log2 ratio) and red elevated during G0 (positive Log2 ratio).

**Supplementary Figure S11.**
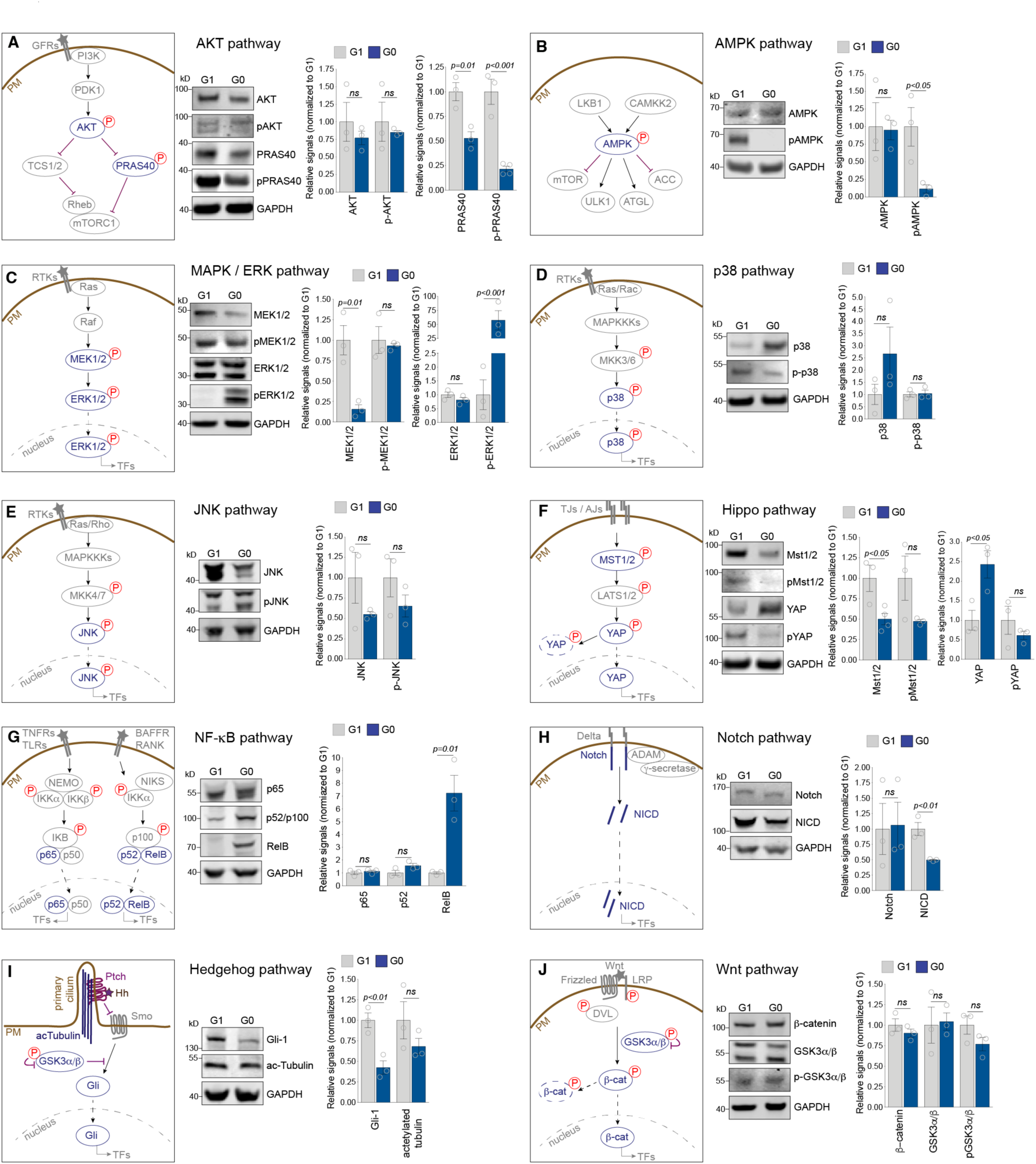
Activity of 10 major signaling pathways during G0. (A) Left, Schematic of the AKT signaling pathway. Middle, representative immunoblots showing the levels in phosphorylated and total AKT and PRAS40 in G1 or G0 cell extracts (isolated by cell sorting, *see STAR Methods*). Uncropped gels are displayed in Supplementary Fig. S16-18. Right, relative signals normalized to GAPDH, of 3 independent experiments as in *(Middle)*. (B) Left, Schematic of the AMPK signaling pathway. Middle, representative immunoblots showing the levels in phosphorylated and total AMPK in G1 or G0 cell extracts (isolated by cell sorting, *see STAR Methods*). Uncropped gels are displayed in Supplementary Fig. S16-18. Right, relative signals normalized to GAPDH, of 3 independent experiments as in *(Middle)*. (C) Left, Schematic of the ERK signaling pathway. Middle, representative immunoblots showing the levels in phosphorylated and total MEK1/2 and ERK1/2 in G1 or G0 cell extracts (isolated by cell sorting, *see STAR Methods*). Uncropped gels are displayed in Supplementary Fig. S16-18. Right, relative signals normalized to GAPDH, of 3 independent experiments as in *(Middle)*. (D) Left, Schematic of the p38 signaling pathway. Middle, representative immunoblots showing the levels in phosphorylated and total p38 in G1 or G0 cell extracts (isolated by cell sorting, *see STAR Methods*). Uncropped gels are displayed in Supplementary Fig. S16-18. Right, relative signals normalized to GAPDH, of 3 independent experiments as in *(Middle)*. (E) Left, Schematic of the JNK signaling pathway. Middle, representative immunoblots showing the levels in phosphorylated and total JNK in G1 or G0 cell extracts (isolated by cell sorting, *see STAR Methods*). Uncropped gels are displayed in Supplementary Fig. S16-18. Right, relative signals normalized to GAPDH, of 3 independent experiments as in *(Middle)*. (F) Left, Schematic of the Hippo signaling pathway. Middle, representative immunoblots showing the levels in phosphorylated and total Mst1/2 and YAP in G1 or G0 cell extracts (isolated by cell sorting, *see STAR Methods*). Uncropped gels are displayed in Supplementary Fig. S16-18. Right, relative signals normalized to GAPDH, of 3 independent experiments as in *(Middle)*. (G) Left, Schematic of the NF-κB signaling pathway. Middle, representative immunoblots showing the levels in total p65, p52/p100 and RelB in G1 or G0 cell extracts (isolated by cell sorting, *see STAR Methods*). Uncropped gels are displayed in Supplementary Fig. S16-18. Right, relative signals normalized to GAPDH, of 3 independent experiments as in *(Middle)*. (H) Left, Schematic of the Notch signaling pathway. Middle, representative immunoblots showing the levels in total Notch and NICD in G1 or G0 cell extracts (isolated by cell sorting, *see STAR Methods*). Uncropped gels are displayed in Supplementary Fig. S16-18. Right, relative signals normalized to GAPDH, of 3 independent experiments as in *(Middle)*. (I) Left, Schematic of the Hedgehog signaling pathway. Middle, representative immunoblots showing the levels in total Gli-1 and actetylated tubulin in G1 or G0 cell extracts (isolated by cell sorting, *see STAR Methods*). Uncropped gels are displayed in Supplementary Fig. S16-18. Right, relative signals normalized to GAPDH, of 3 independent experiments as in *(Middle)*. (J) Left, Schematic of the Wnt signaling pathway. Middle, representative immunoblots showing the levels in total β-catenin and phosphorylated and total GSK3α/β in G1 or G0 cell extracts (isolated by cell sorting, *see STAR Methods*). Uncropped gels are displayed in Supplementary Fig. S16-18. Right, relative signals normalized to GAPDH, of 3 independent experiments as in *(Middle)*.

**Supplementary Figure S12.**
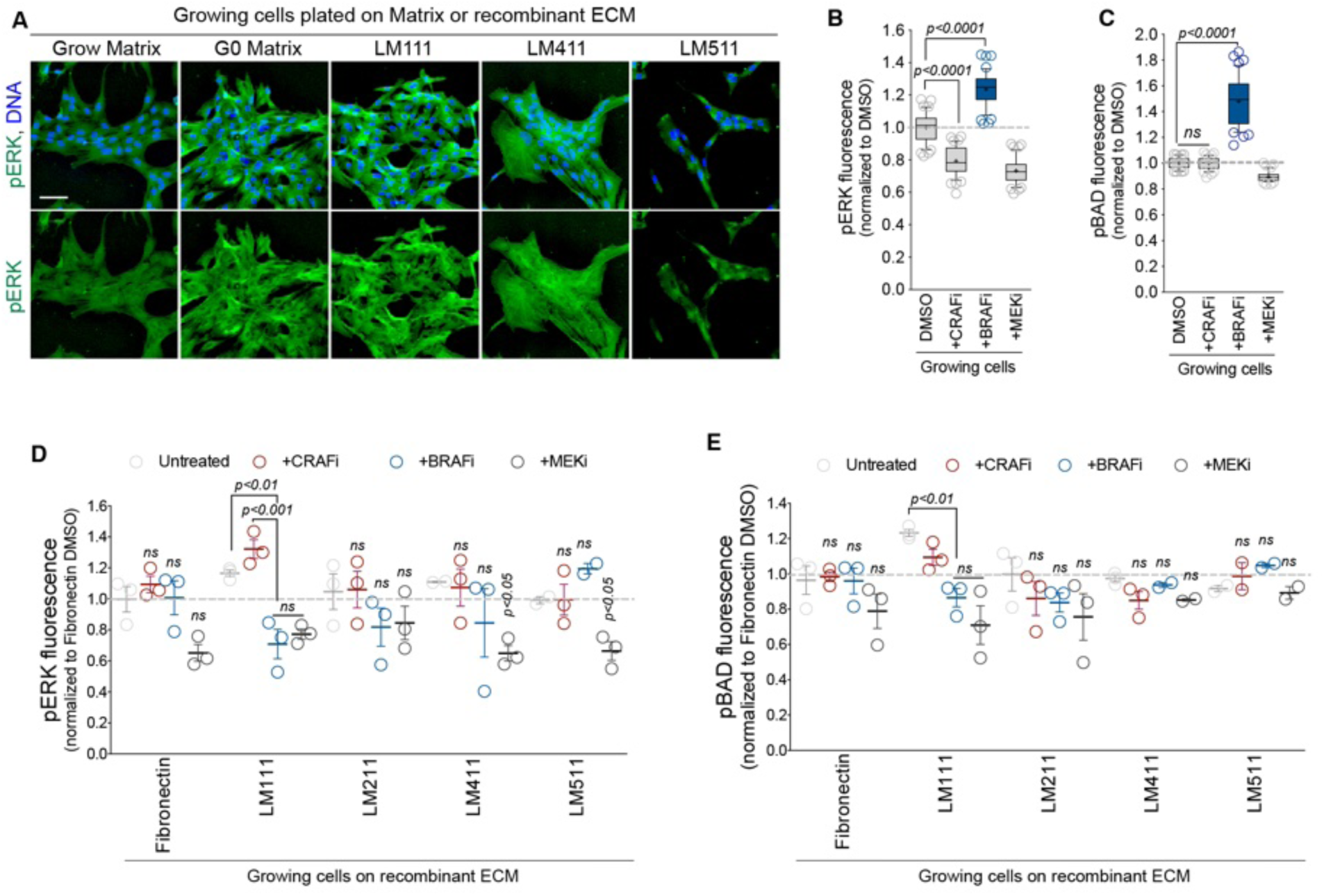
LM111 was sufficient to induce pERK and pBAD elevation through the activation of BRAF. (A) Representative fluorescence microscopy images of growing cells replated onto Grow or G0 Matrix or recombinant LM111, LM411 or LM511 and grown for 6 days in serum-free medium, and immunostained for endogenous pERK (green) and DNA (blue). Scale bar 50 μm. Part of this panel was used in Figure 2H. (B) Mean fluorescence of pERK expressed as percentage changes relative to untreated control samples, in growing cells treated with 2 μM of PD0325901 (MEKi), 2.5 μM of GDC-0879 (BRAFi), or 2 μM of ZM336372 (CRAFi) for 6 hours and immunostained for endogenous pERK. (C) Mean fluorescence of pBAD expressed as percentage changes relative to untreated control samples, in growing cells treated with 2 μM of PD0325901 (MEKi), 2.5 μM of GDC-0879 (BRAFi), or 2 μM of ZM336372 (CRAFi) for 6 hours and immunostained for endogenous pBAD. (D) Mean fluorescence of pERK expressed as percentage changes relative to untreated control samples, from 3 independent experiments done as in (B) and using purified Fibronectin or recombinant LM111, LM211, LM411 or LM511, as indicated. Part of this panel was used in Figure 3G. (E) Mean fluorescence of pBAD expressed as percentage changes relative to untreated control samples, from 3 independent experiments done as in (C) and using purified Fibronectin or recombinant LM111, LM211, LM411 or LM511, as indicated. Part of this panel was used in Figure 3G.

**Supplementary Figure S13.**
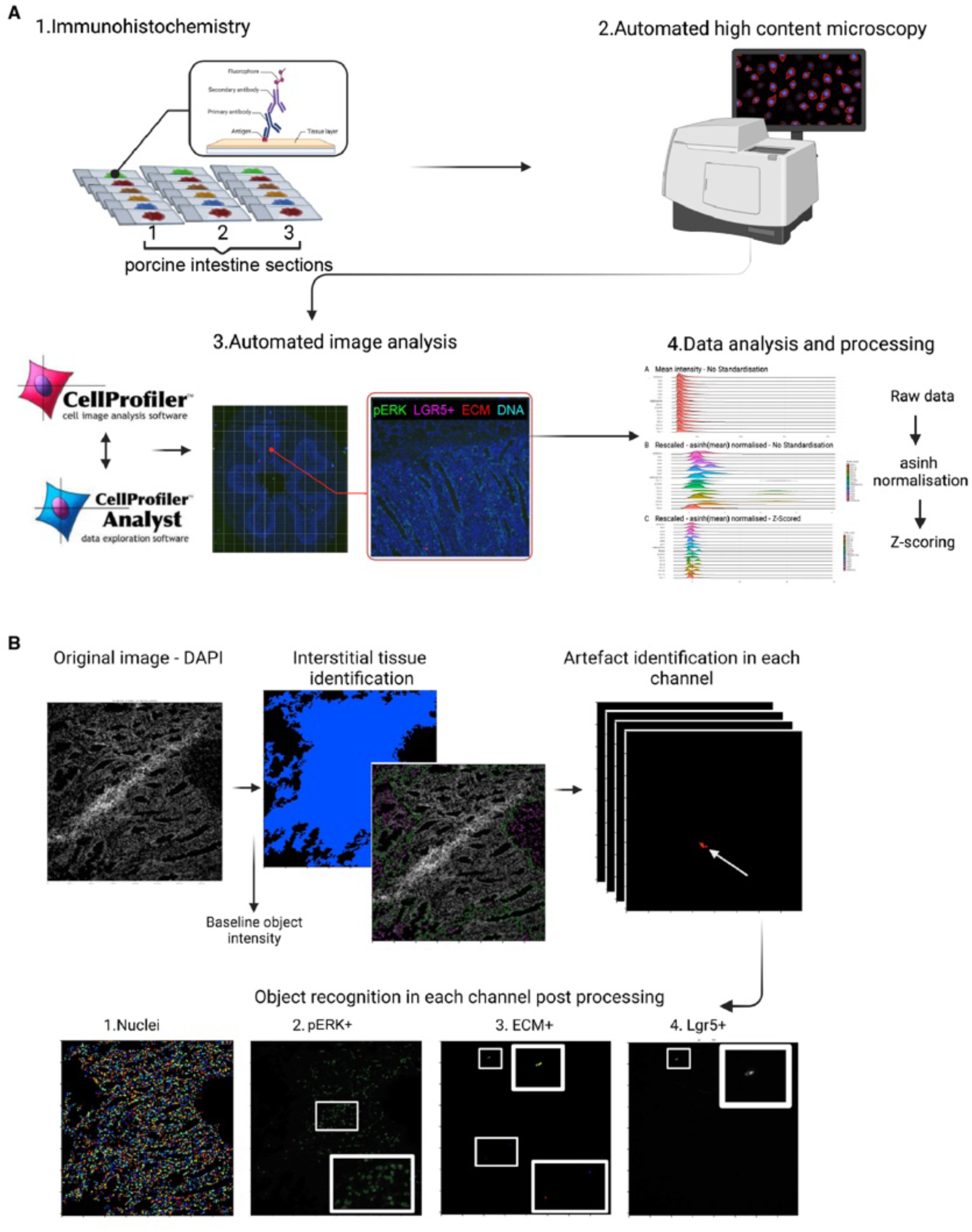
Histology immunostaining, imaging and automated analysis pipeline. (A) Histology immunostaining, imaging and automated analysis pipeline (*see STAR Methods*). (B) Explicit outline of the object recognition pipeline used in CellProfiler (*see STAR Methods*).

**Supplementary Figure S14.**
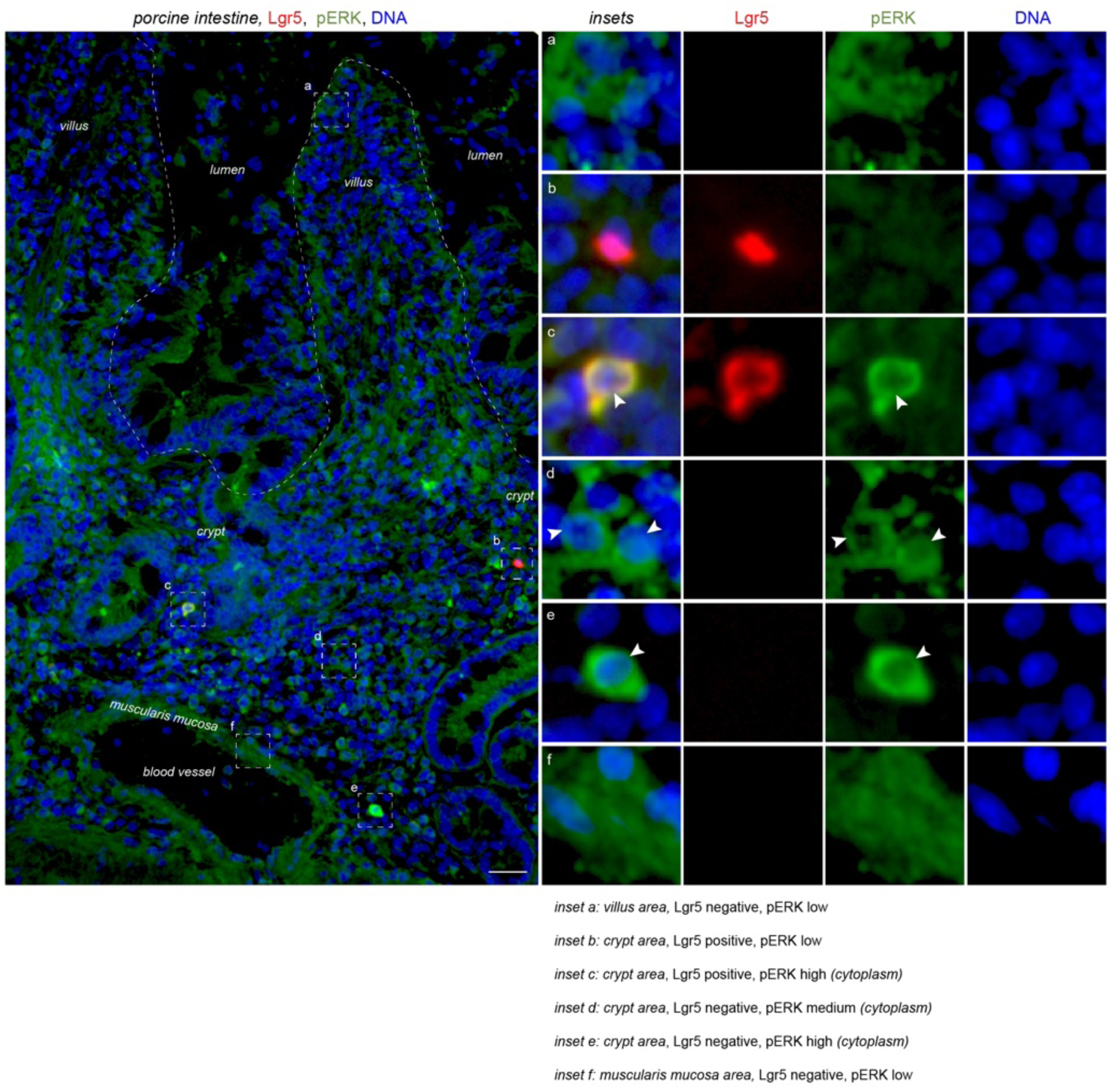
*in vivo* evidence of cytoplasmic pERK in quiescent cells. Representative fluorescence microcopy images of porcine intestine sections immunostained for endogenous pERK (green), stem-cell marker Lgr5 (red) and DNA (blue). Scale bar 50 μm.

**Supplementary Figure S15.**
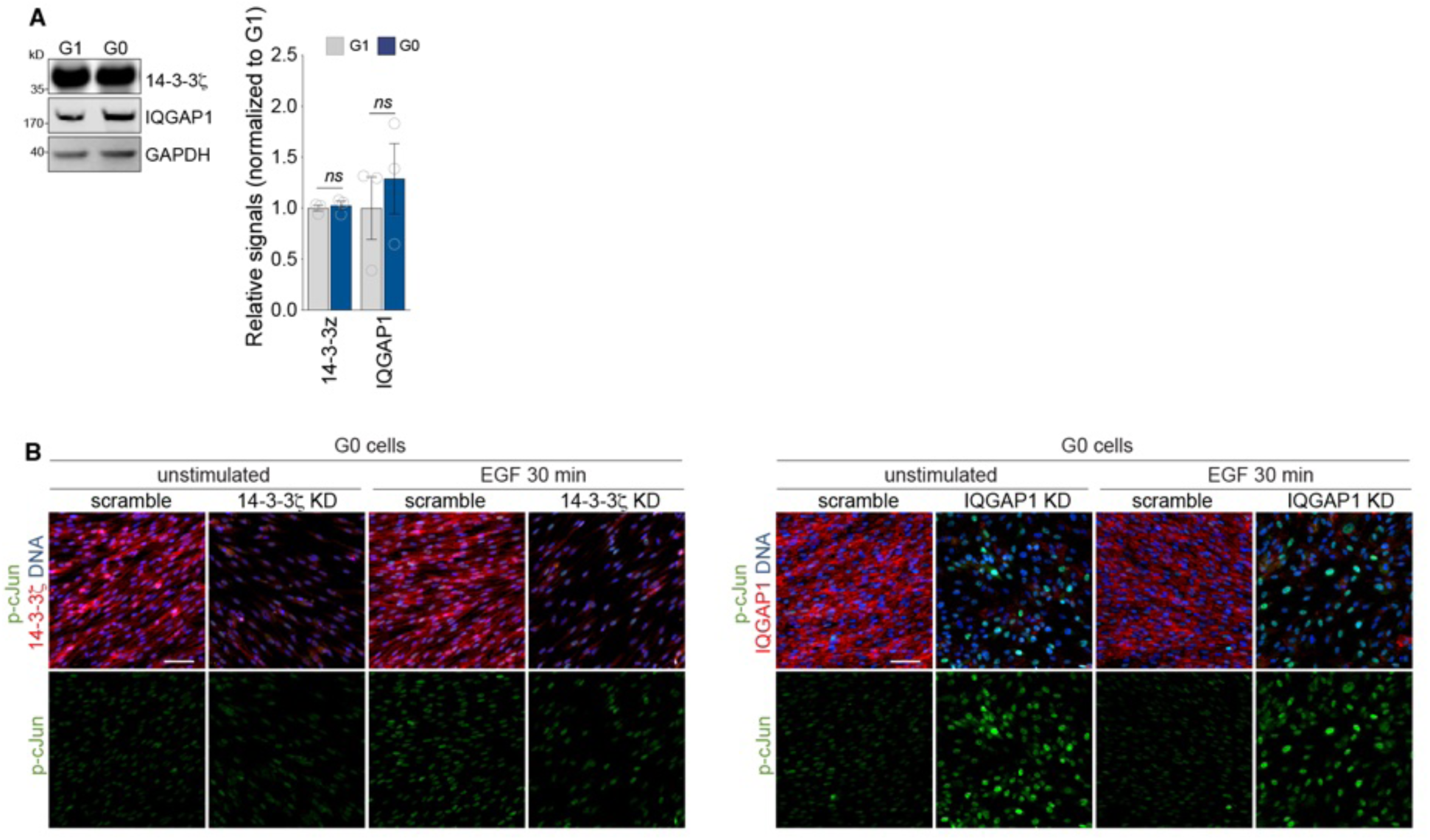
IQGAP1 depletion triggers cJun phosphorylation in G0 cells. (A) Left, representative immunoblots showing the levels in 14-3-3ζ and IQGAP1 in G1 or G0 cell extracts (isolated by cell sorting, *see STAR Methods*). Uncropped gels are displayed in Supplementary Fig. S16-18. Right, relative signals normalized to GAPDH, of 3 independent experiments as in (Left). (B) Representative fluorescence microcopy images of G0 cells depleted of 14-3-3ζ (14-3-3ζ KD) or IQGAP1 (IQGAP1 KD), or treated with control siRNA oligos (scramble), and stimulated with 50ng/mL EGF for 30 min or not (unstimulated), and immunostained for endogenous p-cJun (green), 14-3-3ζ or IQGAP1 (red) and DNA (blue). Part of this panel was used in Figure 5H. Scale bar 50 μm.

**Supplementary Figure S16.**
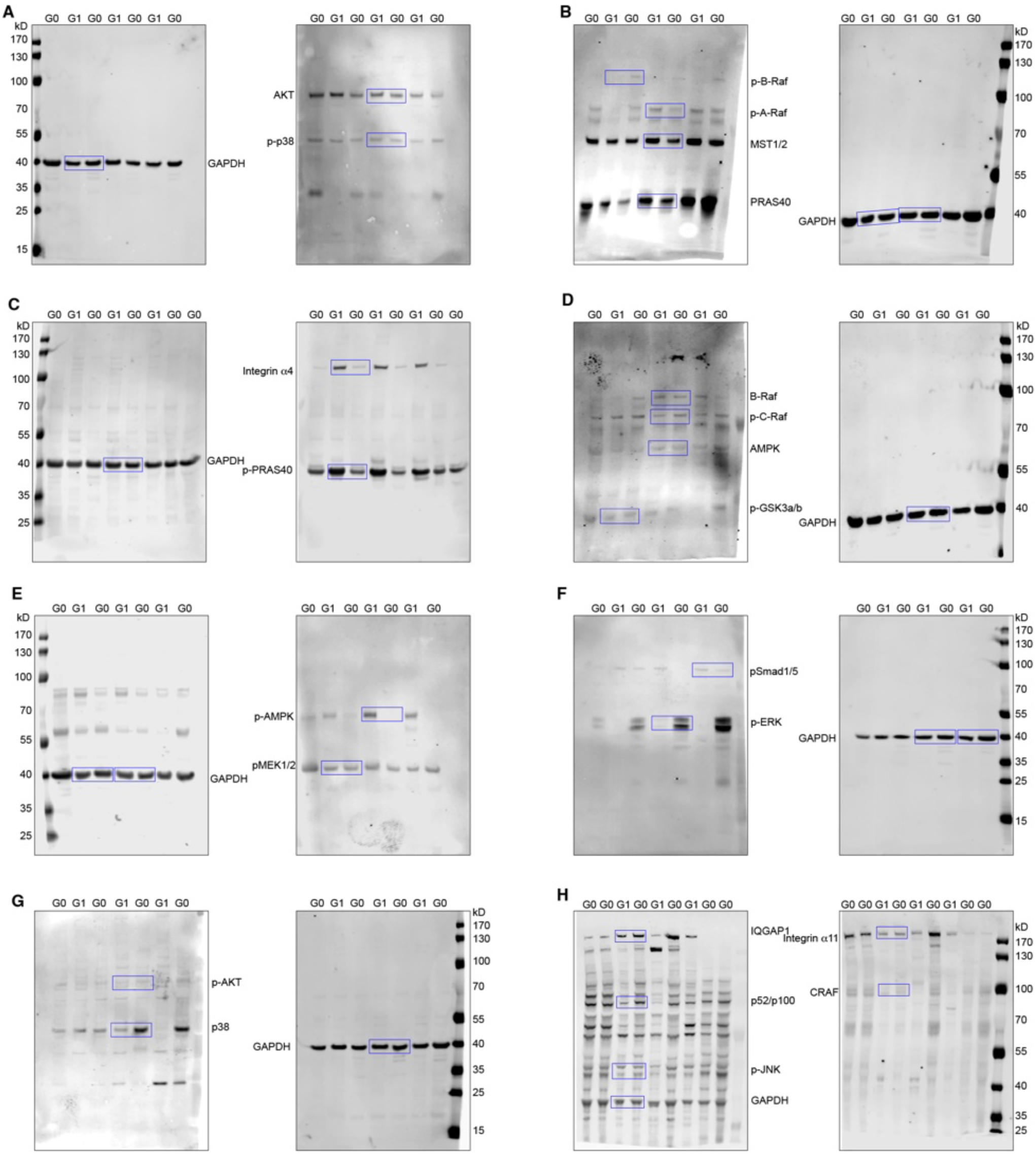
Uncropped gels - part I.

**Supplementary Figure S17.**
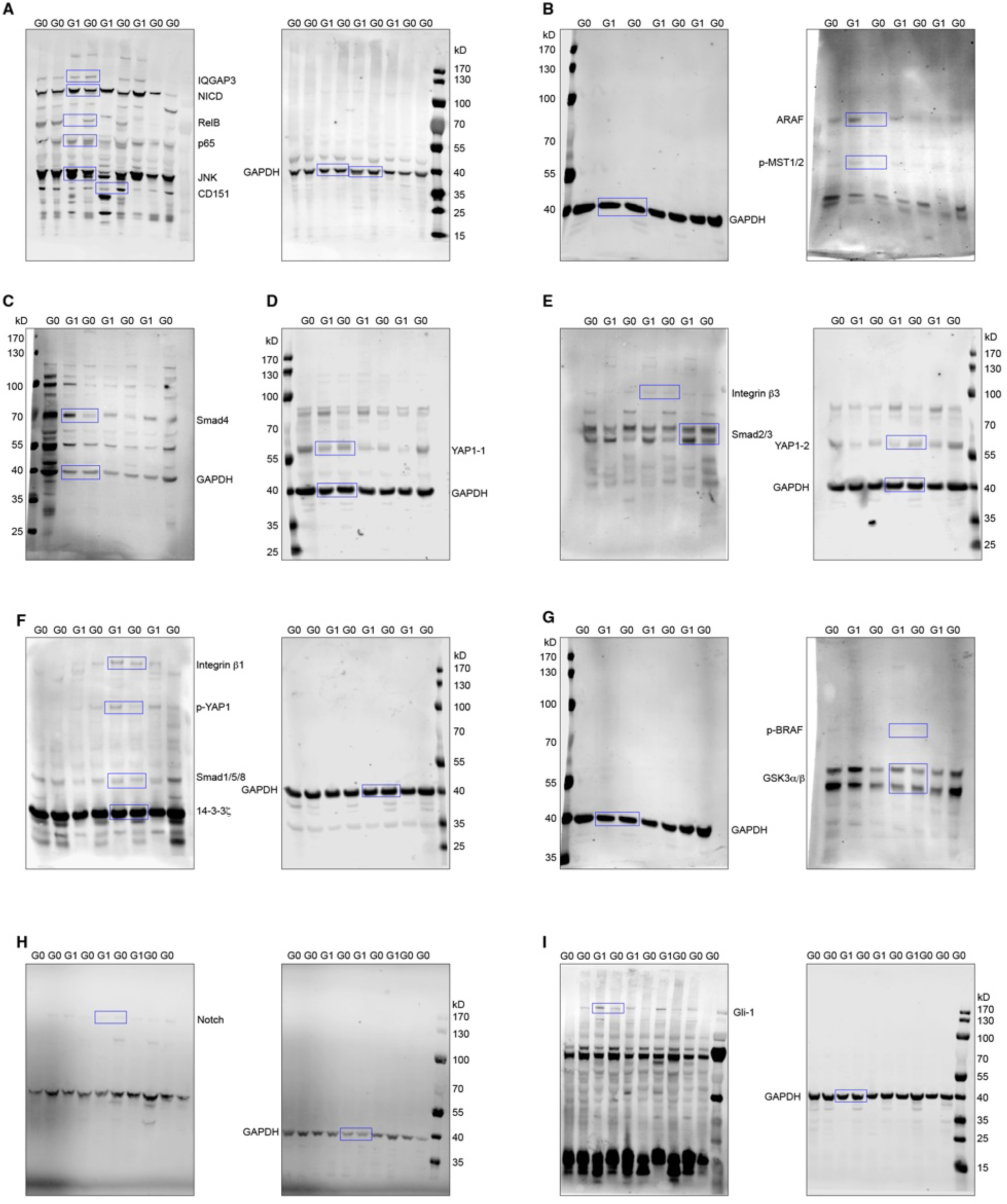
Uncropped gels - part II.

**Supplementary Figure S18.**
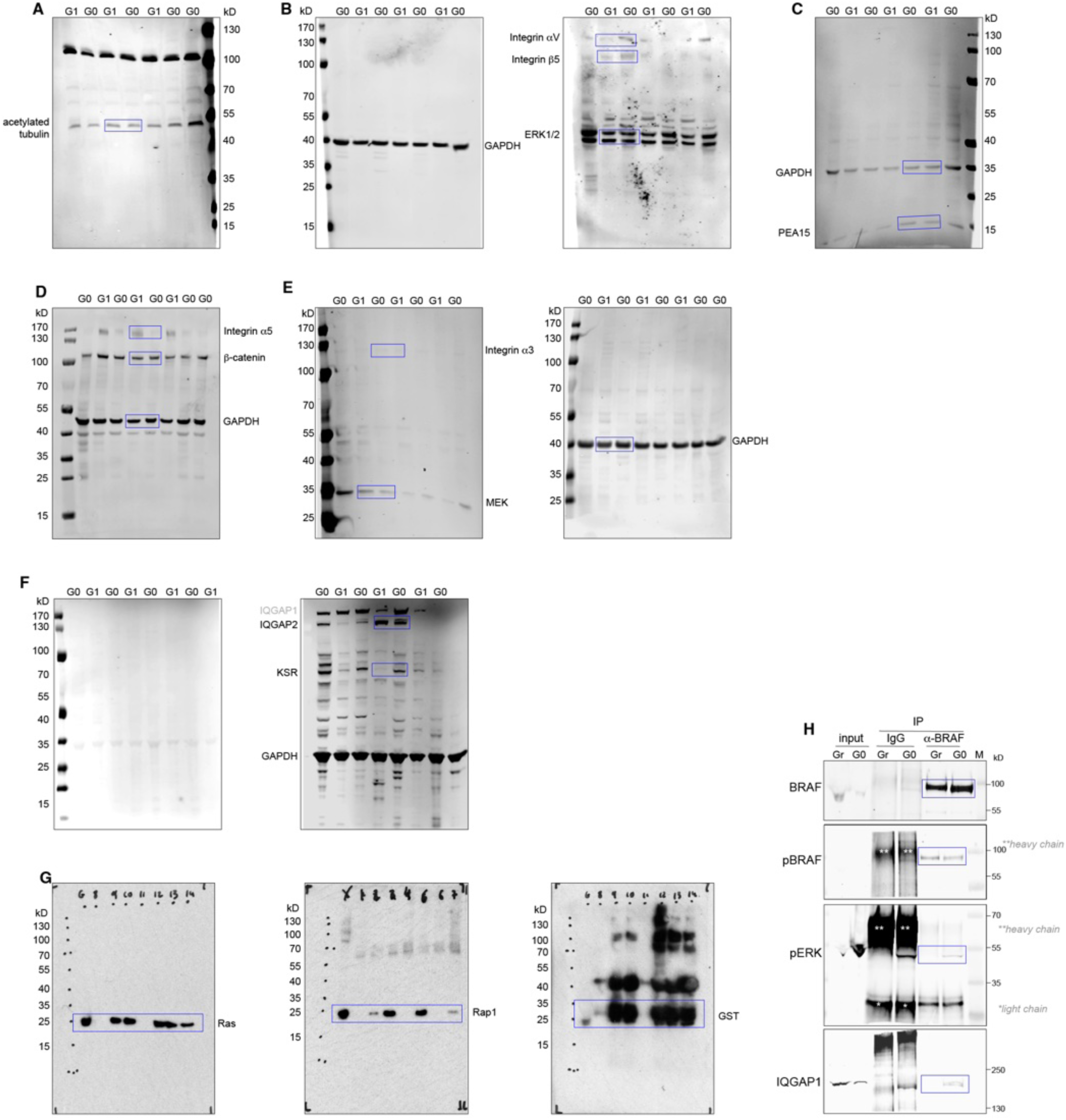
Uncropped gels - part III.

**Supplementary Table 1.** G0 vs Growing Surface and ECM proteomics raw data.

**Supplementary Table 2.** G0 vs G1 SILAC Total Proteome and Phosphoproteome raw data.

